# BESFA: Bioinformatics based Evolutionary, Structural & Functional Analysis of Prostrate, Placenta, Ovary, Testis, and Embryo (POTE) Paralogs

**DOI:** 10.1101/2021.12.20.473416

**Authors:** Sahar Qazi, Bimal Prasad Jit, Abhishek Das, Muthukumarasamy Karthikeyan, Amit Saxena, M.D Ray, Angel Rajan Singh, Khalid Raza, B. Jayaram, Ashok Sharma

**Author notes:** Corresponding Author: Ashok Sharma, PhD, Additional Professor, Laboratory of Chromatin and Cancer Epigenetics, Room No-3029, Department of Biochemistry, All India Institute of Medical Sciences, Ansari Nagar, New Delhi-110029, Bharat (India)., E. mail. Equally contributed (Sahar Qazi, Bimal Prasad Jit).

## Abstract

The POTE family comprises 14 paralogues and is primarily expressed in Prostrate, Placenta, Ovary, Testis, Embryo (POTE), and cancerous cells. The prospective function of the POTE protein family under physiological conditions is less understood. We systematically analyzed their cellular localization and molecular docking analysis to elucidate POTE proteins’ structure, function, and Adaptive Divergence. Our result discerns that group three POTE paralogs (POTEE, POTEF, POTEI, POTEJ, and POTEKP (a pseudogene)) exhibits significant variation among other members could be because of their Adaptive Divergence. Furthermore, our molecular docking studies on POTE protein revealed the highest binding affinity with NCI-approved anticancer compounds. Additionally, POTEE, POTEF, POTEI, and POTEJ were subject to an explicit molecular dynamic simulation for 50ns. MM-GBSA and other essential electrostatics were calculated that showcased that only POTEE and POTEF have absolute binding affinities with minimum energy exploitation. Thus, this study’s outcomes are expected to drive cancer research to successful utilization of POTE genes family as a new biomarker, which could pave the way for the discovery of new therapies.

## Introduction

Cancer testis/Cancer Germline (CT/CG) antigens are observed mostly in adult testis/germline tissues and various cancer tumors, which are highly immunogenic and can be seen as potential candidates for cancer biomarkers and therapeutics. POTE (Prostrate, Placenta, Ovary, Testis, and Embryo) is a primate-specific class of proteins, first discovered by an *in-silico* screening approach using the Expressed Sequence Tags (EST) database^1^. POTE has 14 family members and is grouped into classes based on their similarities. POTE family members are localized on eight different chromosomes: 2, 8, 13, 14, 15, 18, 21, and 22, respectively ^1^,^2^. The POTE proteins are composed of mainly three types of repeats, ANK motifs (33 amino acids), cysteine-rich region (CRR; 37 amino acids), and alpha-helical region of varying length in these paralogs. All the paralogs of the POTE family code for a different number of repeats have revealed that during gene evolution, several members of the POTE gene family comprise an actin-retroposon, which is present at the C-terminal of their ancestral paralogs^1,3^.

POTED (POTE-21) is located on chromosome 21. The first-ever POTE gene discovered codes for a protein of 66kDa composed of three CRR, 5 ANK, and many helical regions^1^. POTEH (POTE-22) and POTE-G (POTE-2C) are two genes highly similar to POTED, the only exception being that the latter does not have alpha-helical motifs in their sequences. POTEH (POTE-22) viz. located on chromosome 22, codes for a 34 kDa protein containing two ANK and four CRR conserved regions (motifs). It has been observed that when these three POTE protein sequences (POTED, POTEH, and POTEG) are aligned, there is approximately 73% similarity showing their high homology^4^. This high homology indicates that these proteins may be similar in their functionality, which would make it easier to illustrate their significance in many diseases such as cancer. When comparing dexterous studies focused on the functionality and specific regions based on evolutionary protein analysis, POTE paralogs have limited literature recapitulation. Harking back to Bera et al. (2006); Barger et al. (2018), the research group suggests that the POTE family is primate-specific and belongs to a cancer-testis antigen (CTA) family and is likely to play a pivotal role in primate biological dynamics. Their analysis paved the way for looking at the POTE members in the direction of cancer-testis antigens (CTAs). Recently, Barger et al. (2018) discerned that POTE groups 1 and 2 encapsulate that POTEA, POTEB, POTEB2, POTEC, and POTED a normal tissue expression that is relevant to cancer-testis antigens (CTAs). However, group 3 paralogs, POTEE, POTEF, POTEG, POTEH, POTEI, POTEJ, POTEKP, and POTEM, have a function in normal tissues and is not considered as cancer-testis antigens (CTAs). This study also somehow clubs our POTE paralogs based on their specificity. When comparing the two-benchmark studies^2,5^, it is evident that a thin layer of vague specificity surrounds POTE paralogs. Henceforth, it is essential to determine the POTE paralogs’ evolutionary spectra, which can help determine their structural and functional aspects.

In this study, we have used intelligence strategies computationally to compute the POTE protein family members’ evolutionary relationship so that a correlation can be deduced between the evolutionary divergence of the POTE proteins and their functionality, accordingly. This study aims to identify whether POTE family members have been exposed to Darwinian selection in the process of evolution. Furthermore, structural predictions, molecular docking of the POTE paralogs to anticancer drugs, and molecular refinement with molecular mechanics/generalized Born surface area (MMGBSA) were computed to understand the POTE stability target receptors. Additionally, these POTE paralogs were subjected to a functionality assessment wherein each paralog’s function was deduced in biological, chemical, and molecular aspects.

## Results

### Sequence Similarity & Alignment

We identified a recent functional divergence in 14 POTE paralogs, as shown in Table 1. Because the duplications were identified through similarity of full-length protein sequences, this method detected functional proteins instead of non-functional ones, where an exception was present only for POTEKP. Our study has successfully identified the homogeneity of the POTE paralogs, not only within themselves but also with other species. We used SMART BLAST to identify our POTE proteins’ similarity on the evolutionary basis and found that POTEE, POTEF, POTEI, and POTEJ are orthologous to C. elegans, D. discoideum AX4, Thale cress, S. cerevisiae S288C. POTEA and POTEH are orthologous to the house mouse. POTEKP is orthologous to zebrafish, C.elegans, fruit fly, Dictyostelium discoideum AX4. There are no orthologous genes of POTEB, POTEB2, POTEB3, POTEC, POTED, POTED, POTEM, and POTEG. The pictorial representations of SMART BLAST results have been shown in Fig 1. It lucidly indicates the fact that POTE paralogs may have a recent divergence from other species to humans. This also reiterates that some of the POTE family members are clusters of orthologous groups (COGs) sharing high similarity with another genus. We can hypothesize that if they have sequence similarities with other species, then it is possible that their functions can be derived from these orthologous. It is not an apocryphal tenet that a protein sequence and structure can describe its appropriate functioning and throw light on its dynamic mechanistic pathways, providing insights to many specialized domains that can be fruitful in drug designing developments for various concerned diseases.

**Table 1:**
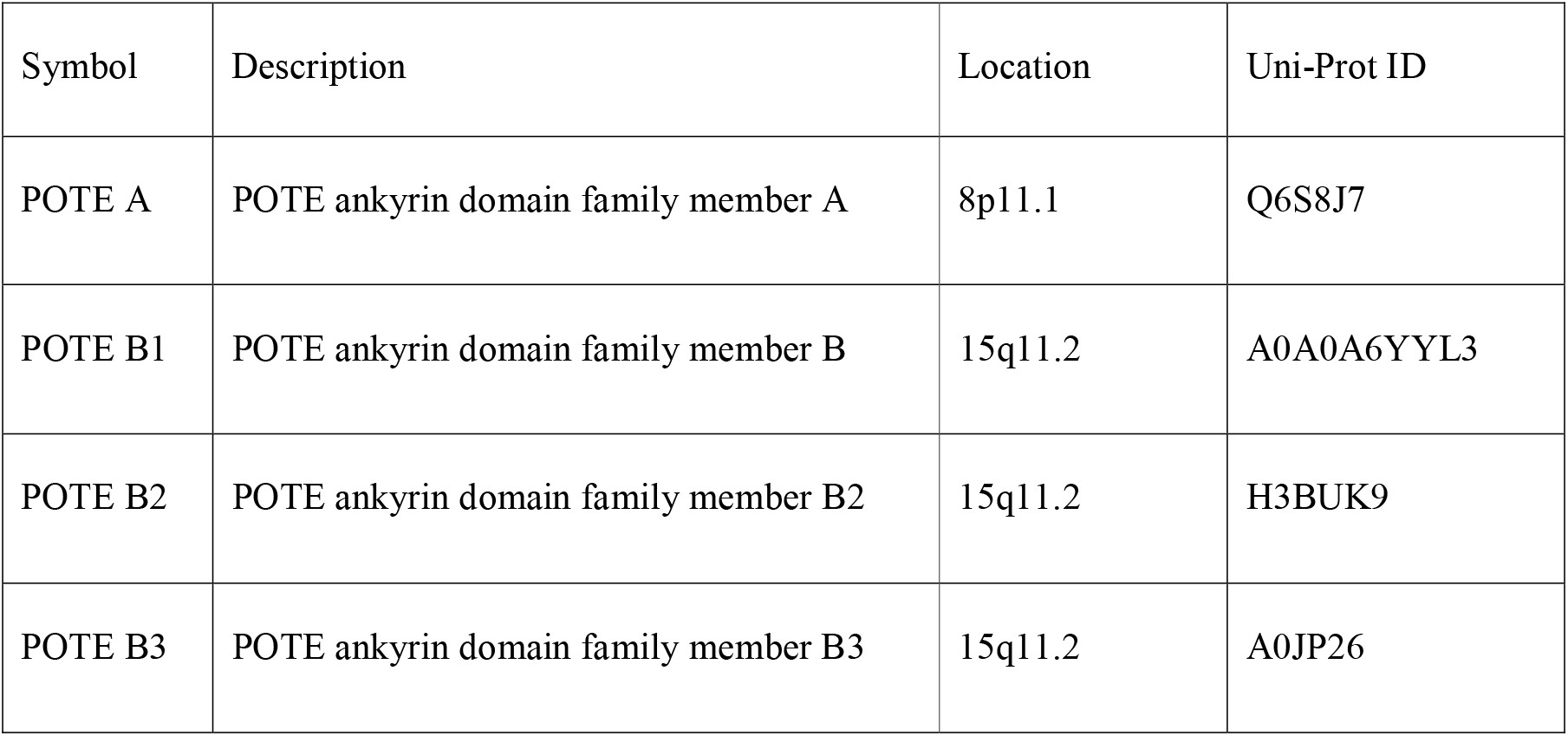
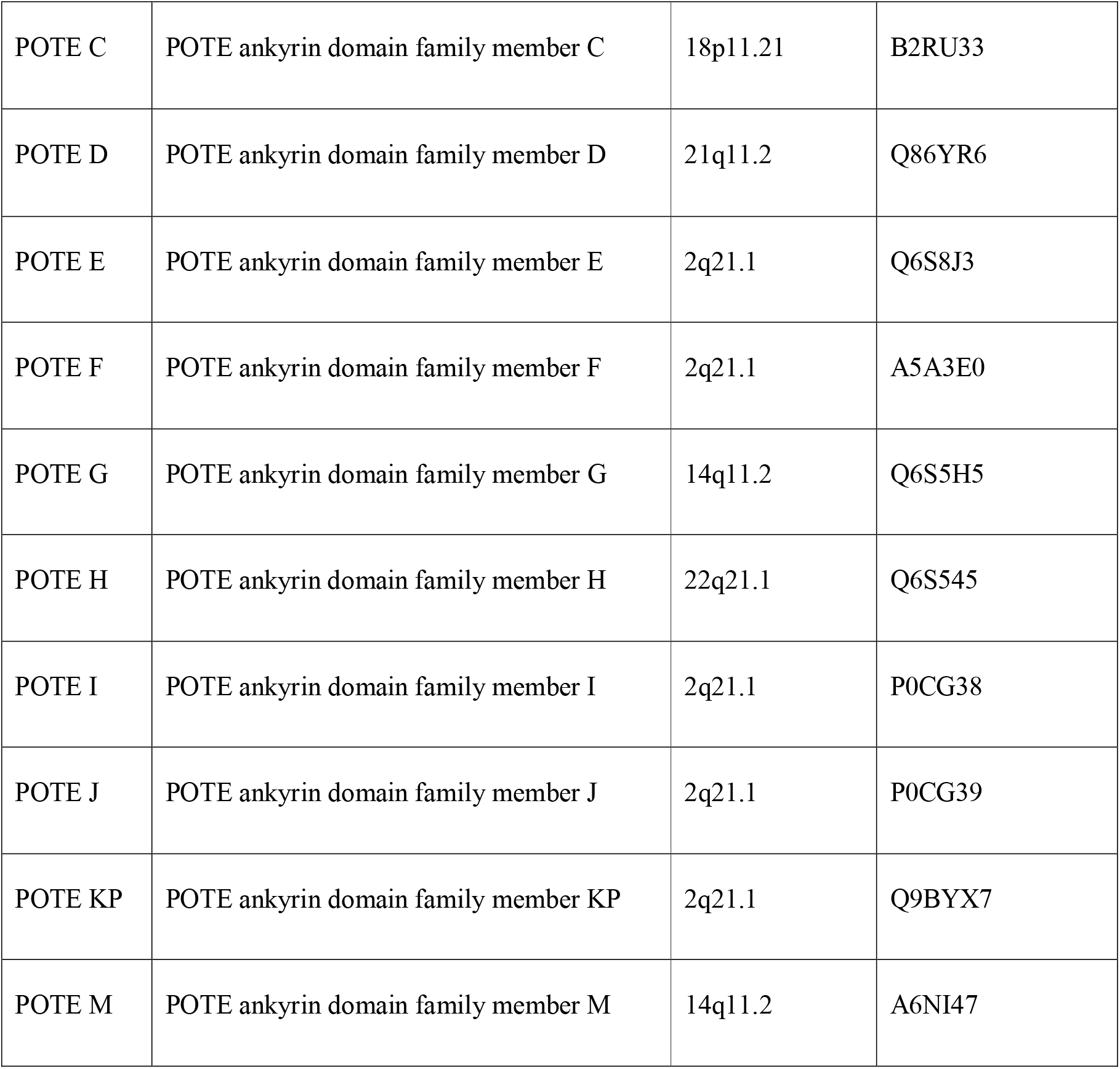
POTE gene family and UniProt accession IDs of POTE paralogs.

**Figure 1(A).**
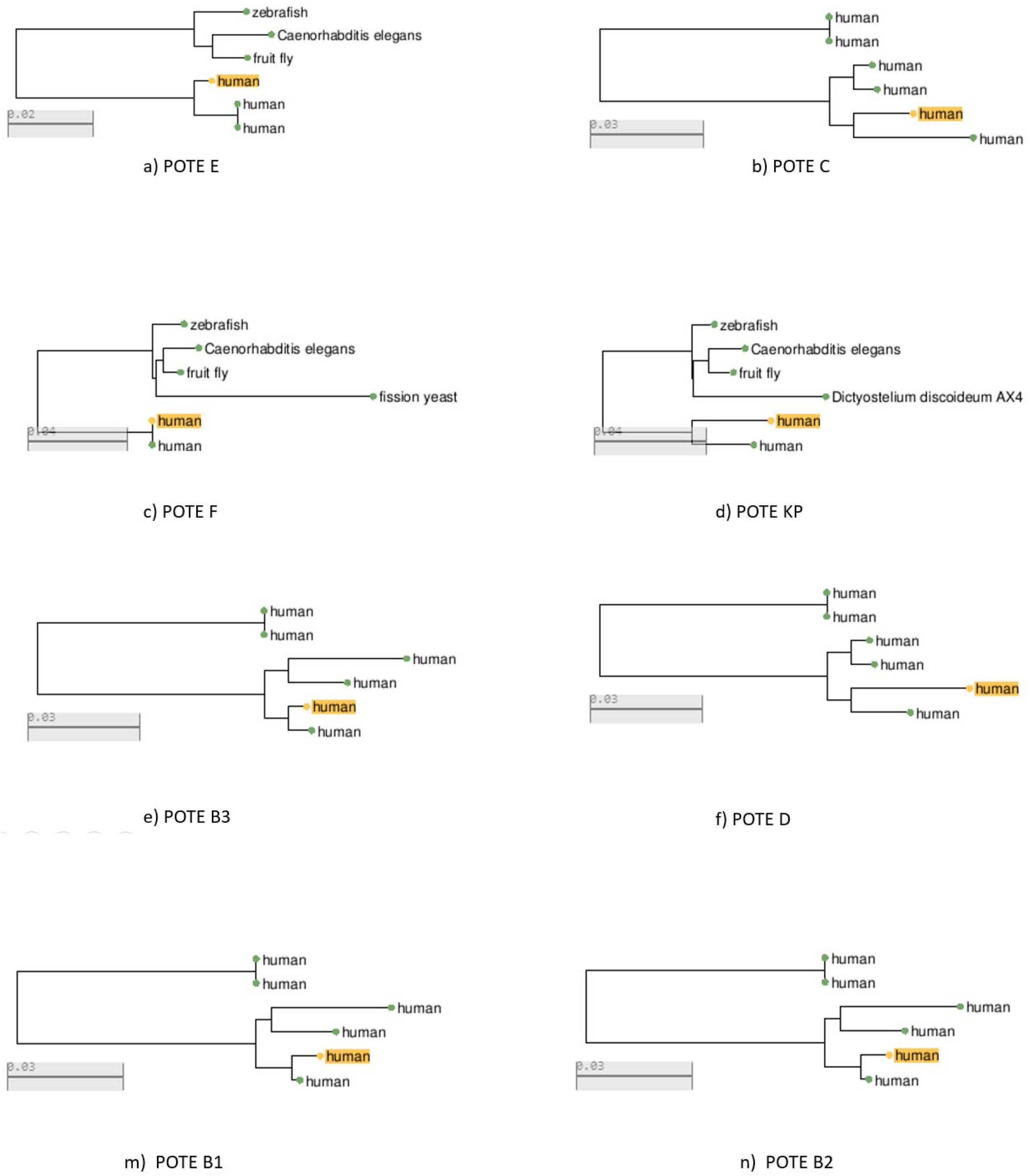
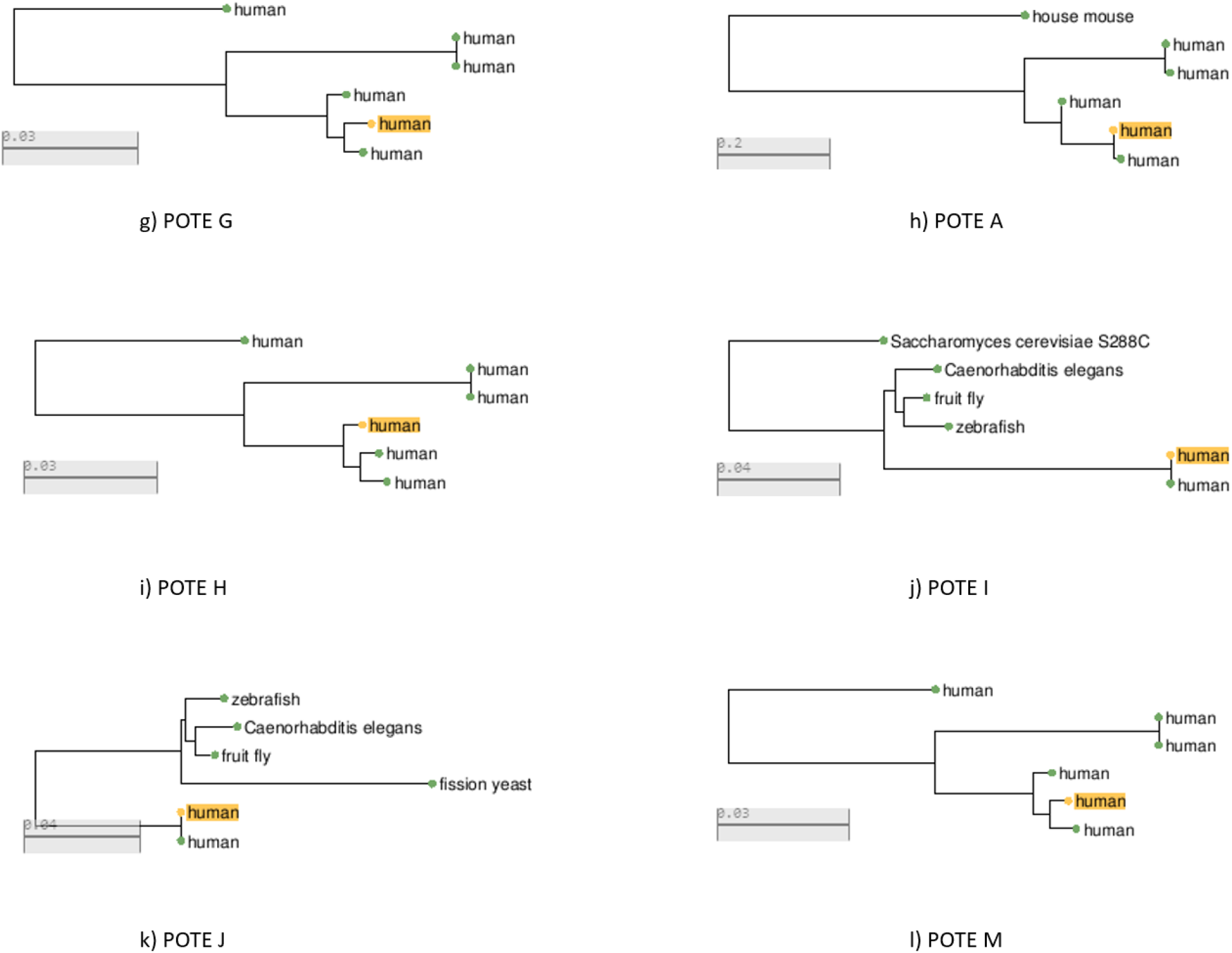
SMART BLAST results of POTE paralogs suggesting homology with other species.

**Figure 1(B).**
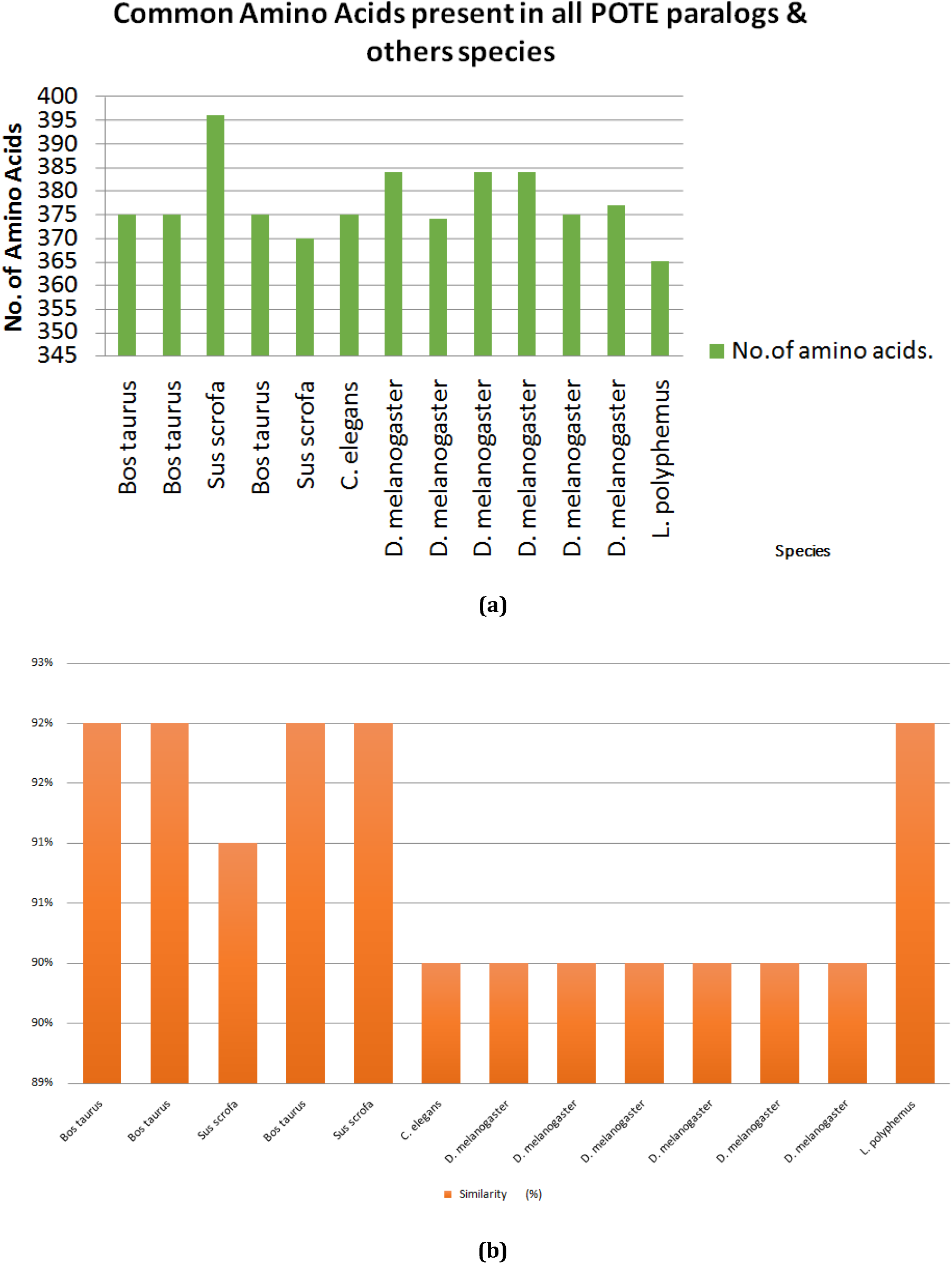
Position specific similarity search for POTE paralogs. a) Graphical representation of the common of amino acids, b) Sequence similarity (%) of various species with POTE paralogs by PSI-BLAST.

**Figure 1(C).**
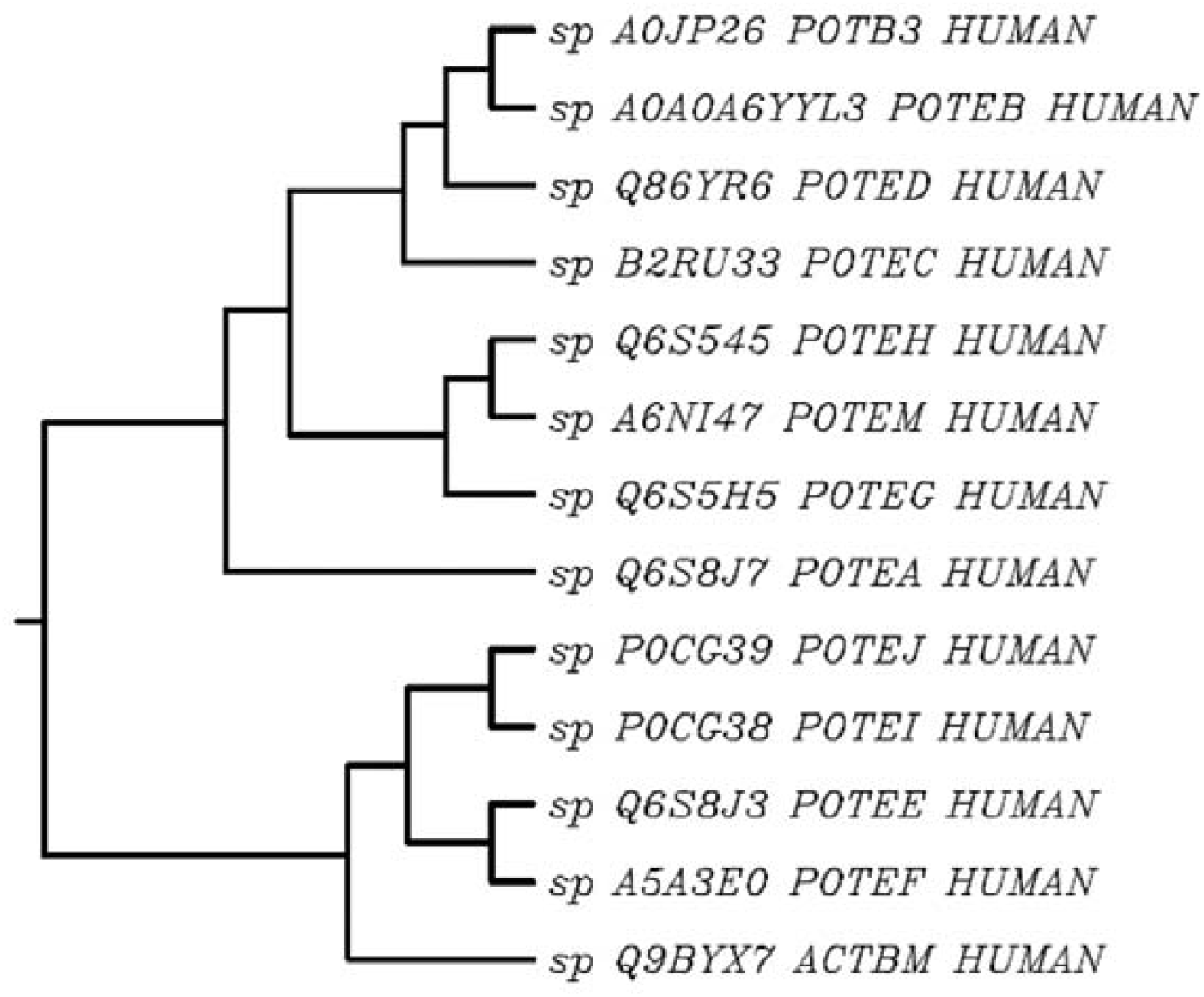
Cladogram of POTE paralogs by CLUSTAL W.

**Figure 1(D).**
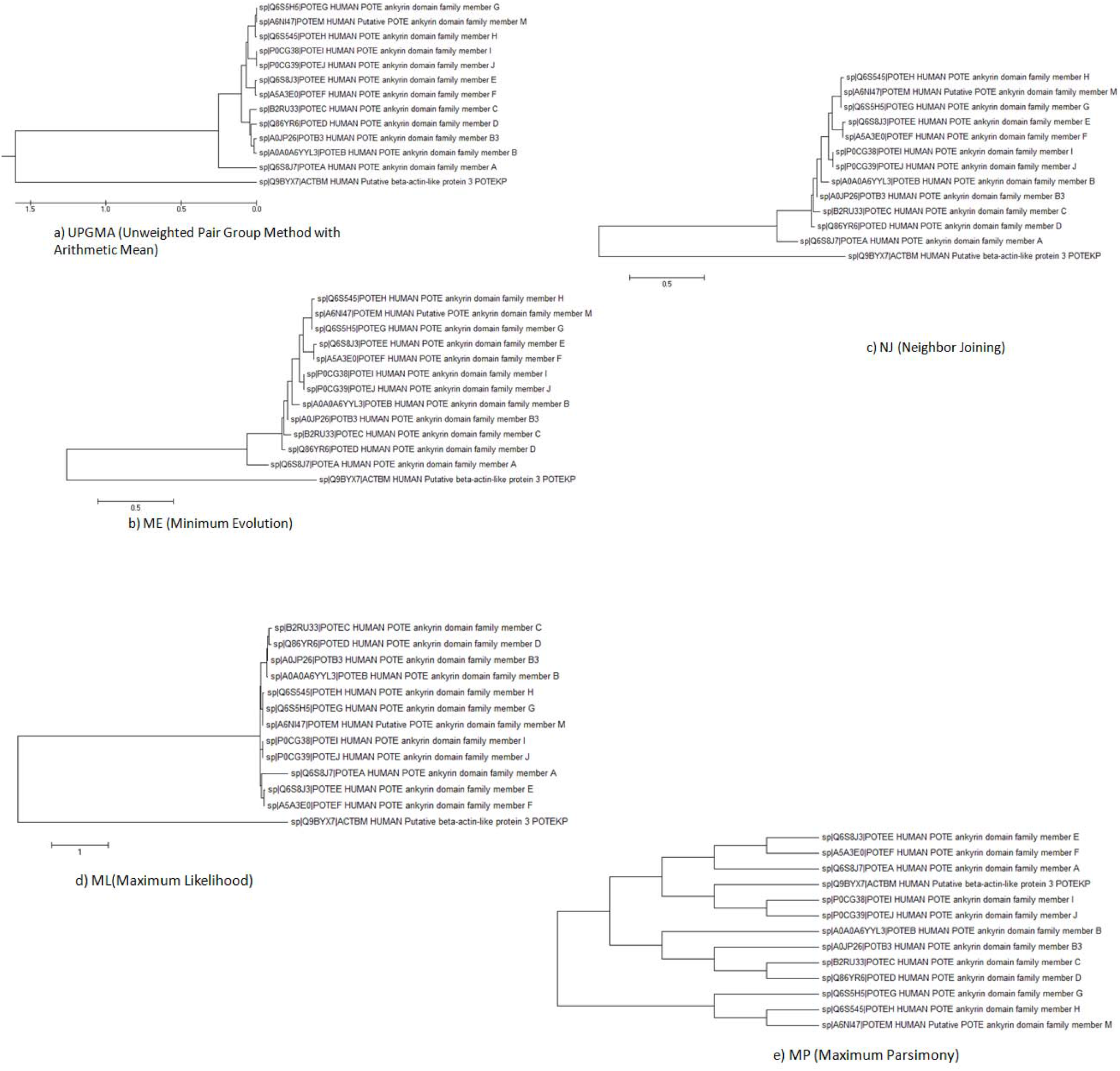
Phylogenetic trees of POTE paralogs using a). UPGMA, b). Minimum Evolution, c) Neighbor Joining, d). Maximum Likelihood and e). Maximum Parsimony.

PSI-BLAST is a statistically driven protein similarity search method that hunts regions of similarity between the query sequence and landmark database sequences and generates gapped alignments. The PSI-BLAST program is more sensitive than BLAST because it can find distantly related sequences missed in a BLAST search. It can repeatedly search the target landmark databases such as as-nr (non-redundant) using multiple alignments of high-scoring sequences found in each iteration to produce a new PSSM for the next round. The program iterates until no new sequences are found or if the threshold is achieved. The PSI-BLAST results show similar domains in the query sequence and similar sequence hits retrieved by the program (Table 2). The results retrieved by PSI-BLAST indicate the fact that all the 14 POTE paralogs have high similarity with *Bos taurus* (3U4L_A, 2OAN_A, 2BTF_A), *Sus scrofa* (5NW4_V & 5AFT_H), *Drosophila melanogaster* (4JHD_B, 4JHD_A, 4RWT_A, 2HF3_A, 3EKS_A & 4M63_C), *C. elegans* (1D4X_A) and *Limulus polyphemus* (3B63_L) and a total number of amino acids in these species as retrieved by PSI-BLAST (Fig. 2).

**Table 2.** Highest sequence similarity of POTE paralogs with various organisms other than *Homo sapiens* with accession numbers have been provided by PSI-BLAST having significant expectation values (E-value).

| Organism Name | Accession Id | Query Coverage | E-Value | Similarity (%) | No.of amino acids. |
| --- | --- | --- | --- | --- | --- |
| *Bos taurus* | 3U4L_A | 34% | 0.0 | 92 % | 375 |
| *Bos taurus* | 2OAN_A | 34% | 0.0 | 92% | 375 |
| *Bos taurus* | 2BTF_A | 34% | 0.0 | 92% | 375 |
| *Sus scrofa* | 5AFT_H | 34% | 0.0 | 92% | 370 |
| *Sus scrofa* | 5NW4_V | 34% | 0.0 | 91% | 396 |
| *C.elegans* | 1D4X_A | 34% | 0.0 | 90% | 375 |
| *D. melanogaster* | 4RWT-A | 34% | 0.0 | 90% | 384 |
| *D. melanogaster* | 2HF3_A | 34% | 0.0 | 90% | 374 |
| *D. melanogaster* | 4JHD_A | 34% | 0.0 | 90% | 384 |
| *D. melanogaster* | 4JHD_B | 34% | 0.0 | 90% | 384 |
| *D. melanogaster* | 3EKS_A | 34% | 0.0 | 90% | 375 |
| *D. melanogaster* | 4M63_C | 34% | 0.0 | 90% | 377 |
| *L. polyphemus* | 3B63_L | 34% | 0.0 | 92% | 365 |

**Figure 2(A).**
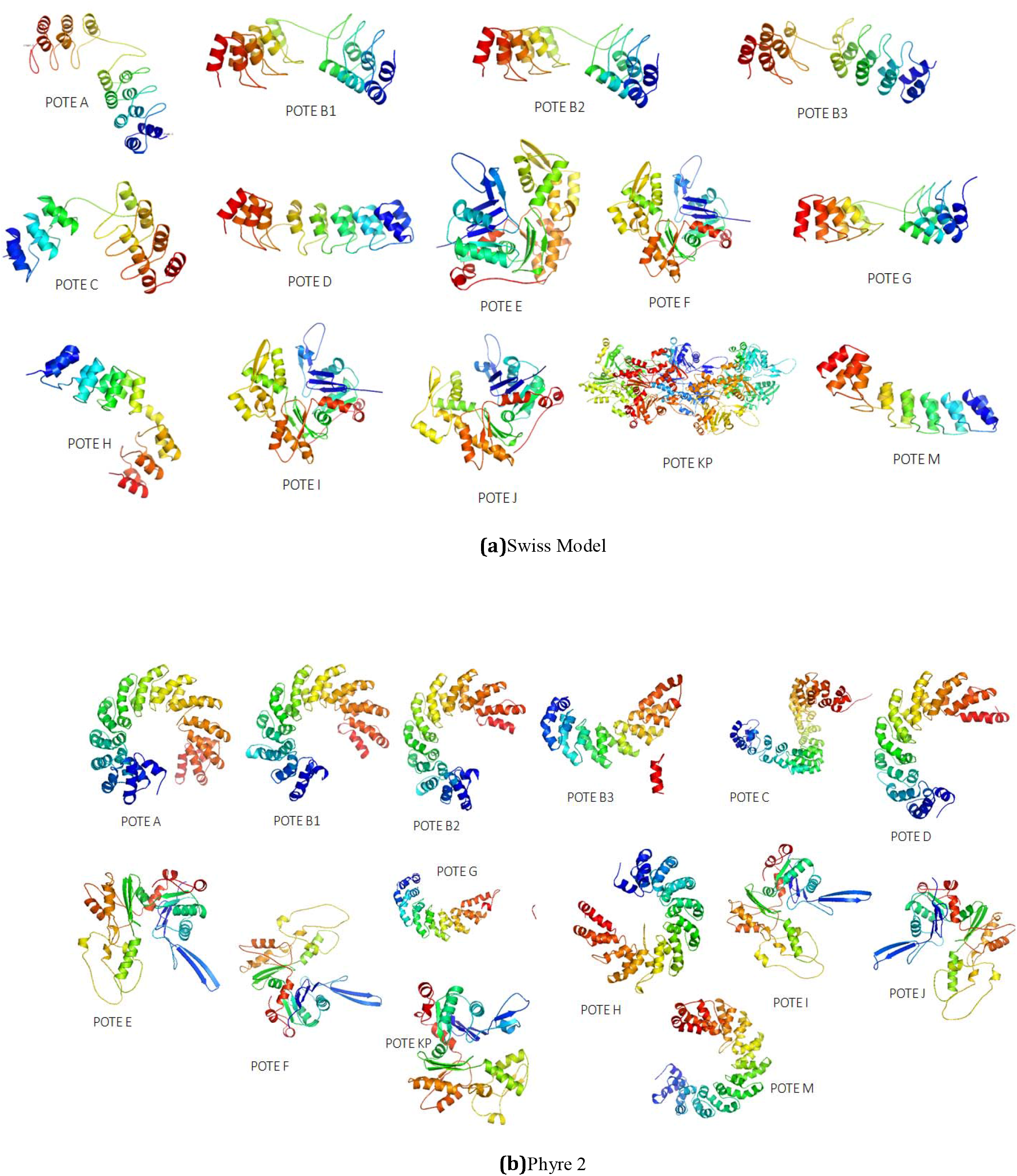
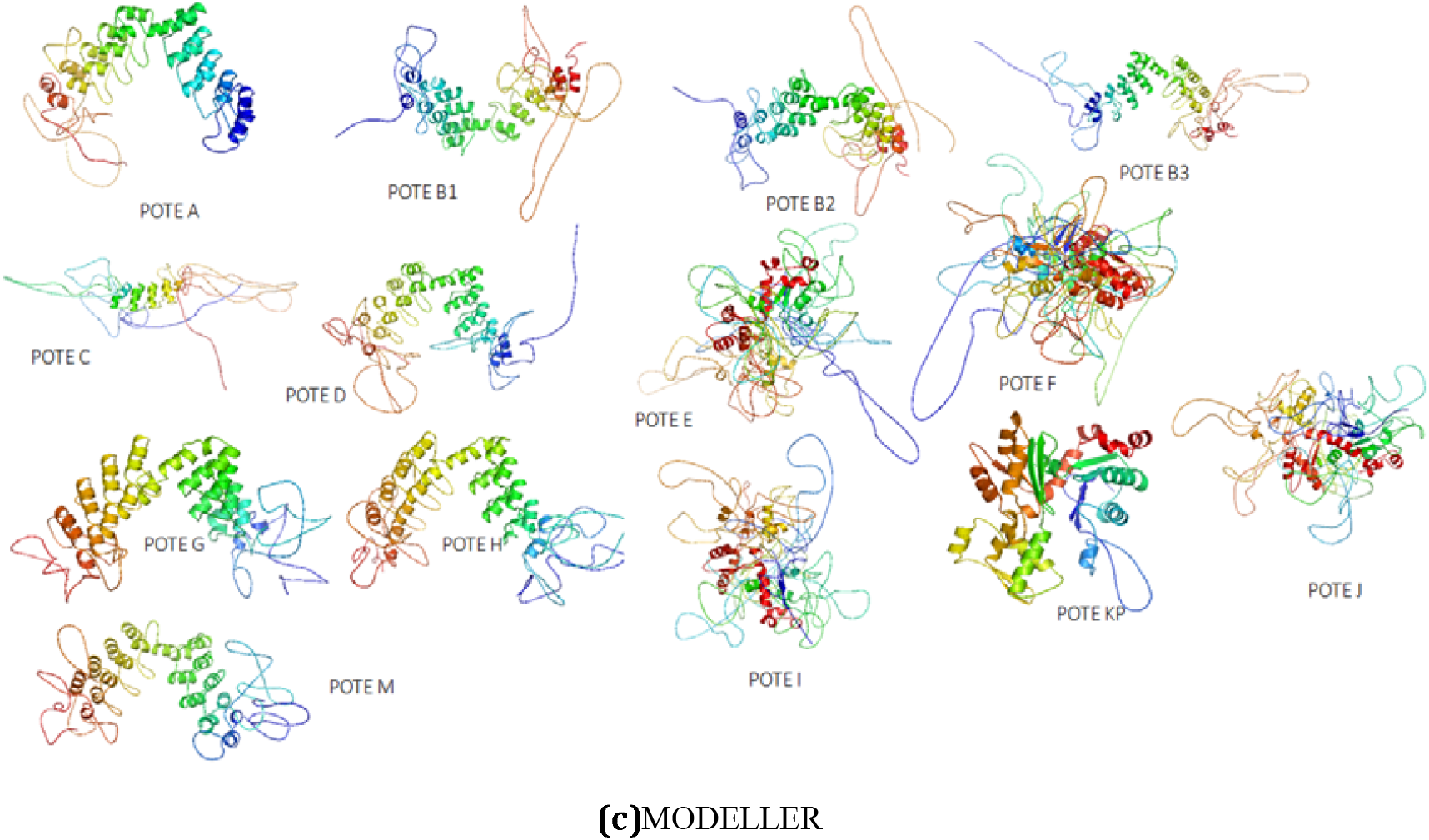
Tertiary models developed by using a). Swiss Model, (b) Phyre2, and c) MODELLER.

**Figure 2(B).**
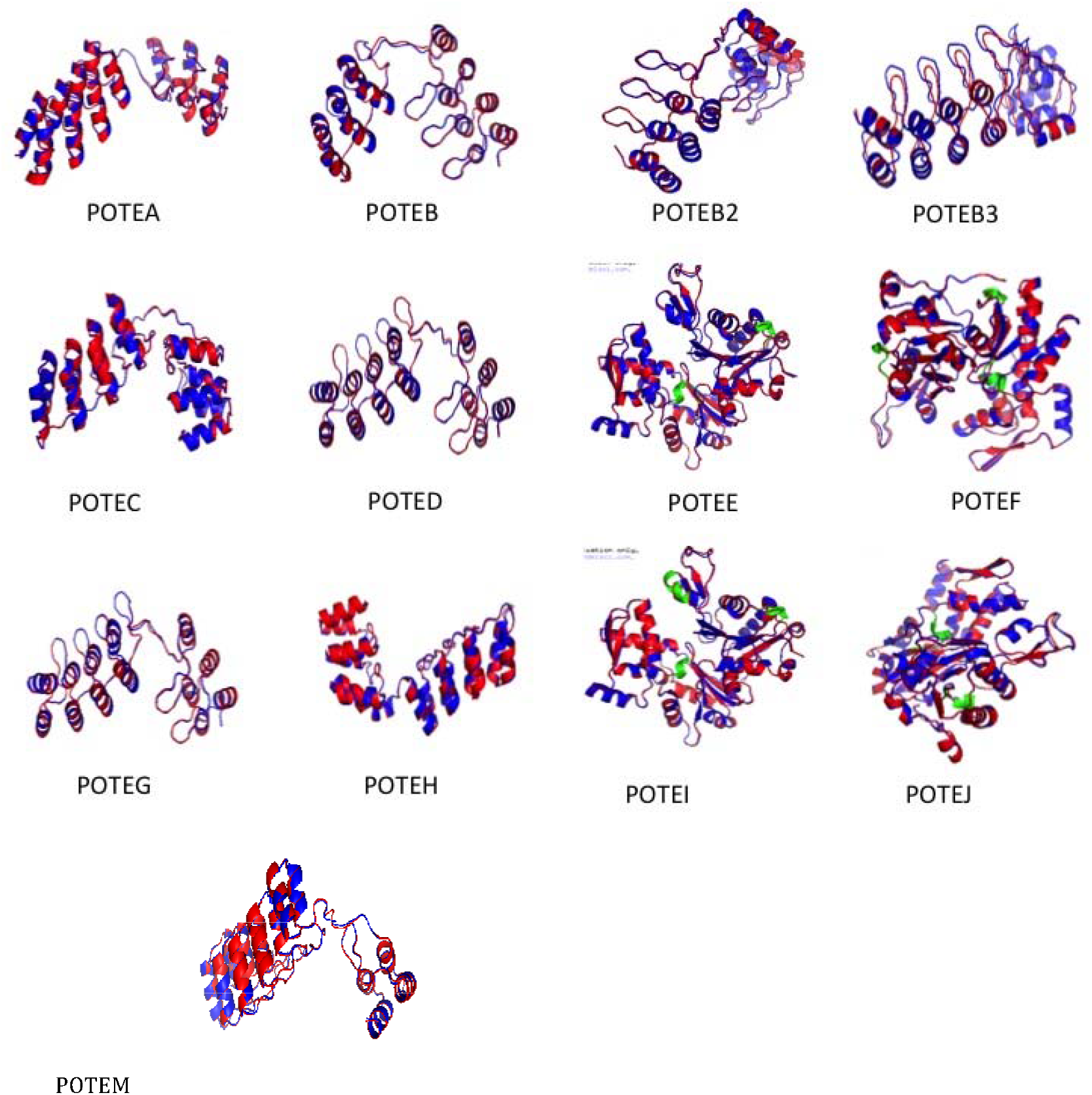
The structures superimposed of pre refinement (red) and post refinement (blue) is shown in detailed manner.

PSI-BLAST may successfully determine some subtle relationships that surpass the standard database in similarity searches but is dependent on the amino acid pattern, viz. conserved within the protein family of interest. We chose only the highest hits retrieved by the PSI-BLAST algorithm, and in each case; it reports a simple but structurally and functionally relevant relationship between humans and other species such as-*D. melanogaster, C. elegans, L. Polyphemus, B. Taurus, and S. scrofa* as these are optimal hits highly significant with an E-value of 0. The alignments suggest that these relationships have clear family members, henceforth hinting for further research analysis on the reliability and correlation of these protein hits with our POTE protein sequences query. POTE proteins, which have been presumed to share an evolutionary relationship, descend from a common root or origin. Thus, a multiple sequence alignment using Omega, Muscle, ClustralW & MEGA 5.02 was executed to infer the sequence homology and phylogenetic analysis. The results discern mutational events such as point mutations occurring at different locations as different characters in a single alignment column and insertion or deletion mutations (indels or gaps), which appear as hyphens in one or more of the sequences alignment. Since POTEB and POTEB2 are the same and equivalent in length, they have been tagged simply as POTEB while keeping POTEB3 separately, which is lengthier.

ClustalW, along with the alignment, also predicts phylogeny, shown in Supplemental Figure S2. The phylogenetic analysis executed by CLUSTALW discerns that these proteins have diverged at a slow pace and clubbed into five specific groups shows that Fig 3. Although POTEA is quite close to POTEB, POTEB3, POTEC, POTED, POTEH, POTEM, and POTEG, it has diverged and separated itself from the rest, hinting at its mono-existence. The reason being that POTEA has a varying structural skeleton; thus, in group I. POTEB, POTEB2, POTEB3, POTEC, and POTED are closer to one another because of highly similar structural framework and hence makes group II. POTEH, POTEM, and POTEG make group III as they are clubbed together because of the same ancestral root and similarity. POTEE, POTEF, POTEI, and POTEJ, however, make a visibly distinct group IV arising from a common root but segregating to sub clubs of POTEE and POTEF and POTEI and POTEJ, referring again to their similar framework. POTEKP is a non-functional protein that rises from a pseudogene and is bereft and an outlier with the lowest similarity with the other paralogs.

Phylogenetic analysis using Molecular Evolutionary Genetics Analysis (MEGA) provided a better overview of POTE’s alignment and phylogenetic by giving an edge to its slow divergence and conserved regions. Multiple sequence alignment by MEGA showcases the gapped alignments and stringently conserved regions in all the 14 paralogs as shown in Supplemental Figure S3 by marking them in similar color and letter. At the same time, the mismatches have been highlighted with a different color and letter. Phylogenetic tree construction discerns that POTEE, POTEF, POTEI, and POTEJ are clubbed together, hinting at their common divergence from other species during evolution, while POTEB, POTEB2, POTEB3, POTEC, POTED, POTEG, POTEH, and POTEM are placed relatively nearer to each other. POTEKP and POTEA are placed straightforwardly as outliers and nearer to one another, showing that their ancestral origin might be linked, as shown in Fig 4. The phylogeny was deduced using UPGMA, Neighbor-Joining, Minimum Evolution, Maximum Parsimony, and Maximum Likelihood algorithms, respectively, which are based on distance and cladistics approaches, respectively. All the five phylogenetic trees were obtained to hint at the concept of adaptive evolution in POTE paralogs.

### Secondary Structure Predictions

The secondary structures of all the POTE paralogs were developed by using PSIPRED software. The secondary structure prediction result is shown in Supplemental Figure S4, which indicates that the percentage of β-strands is much more significant in POTEE, POTEF, POTEJ, POTEI, and POTEKP paralogs than the percentage of β-strands in other paralogs, which makes our evolutionary analysis more subjective as a concept for adaptive divergence.

### Structural Predictions& Molecular Dynamic Simulations (MDS)

Based on homology and protein threading strategies, POTE paralogs protein structures were generated using the Swiss Model, MODELLER, and Phyre2. The templates are taken for the Swiss Model, and query coverage, e-value, and sequence identity have been mentioned in Supplemental Table 1. It was observed that all the protein paralogs comprised mainly of helices and supercoiled regions. Only POTEE, POTEF, POTEJ, and POTEI were the only paralogs, which contained β-strands along with the helices and coils in their tertiary structures, which matches the evolutionary analysis executed. POTEE, POTEF, POTEI, and POTEJ are clubbed together; thus, they have similar structures. POTEKP also contains β-strands. The POTE family’s tertiary models and structures were developed using the Swiss Model, Phyre 2, and MODELLER, as shown in Fig 5. We refined the models created using the Swiss Model since these tertiary models had a better overall quality than Phyre2 and MODELLER, predicted by Protein Quality (ProQ), Protein Structure Analysis (ProSA), and RAMPAGE. The resolution change has been represented in Table 3.

**Figure 5.**
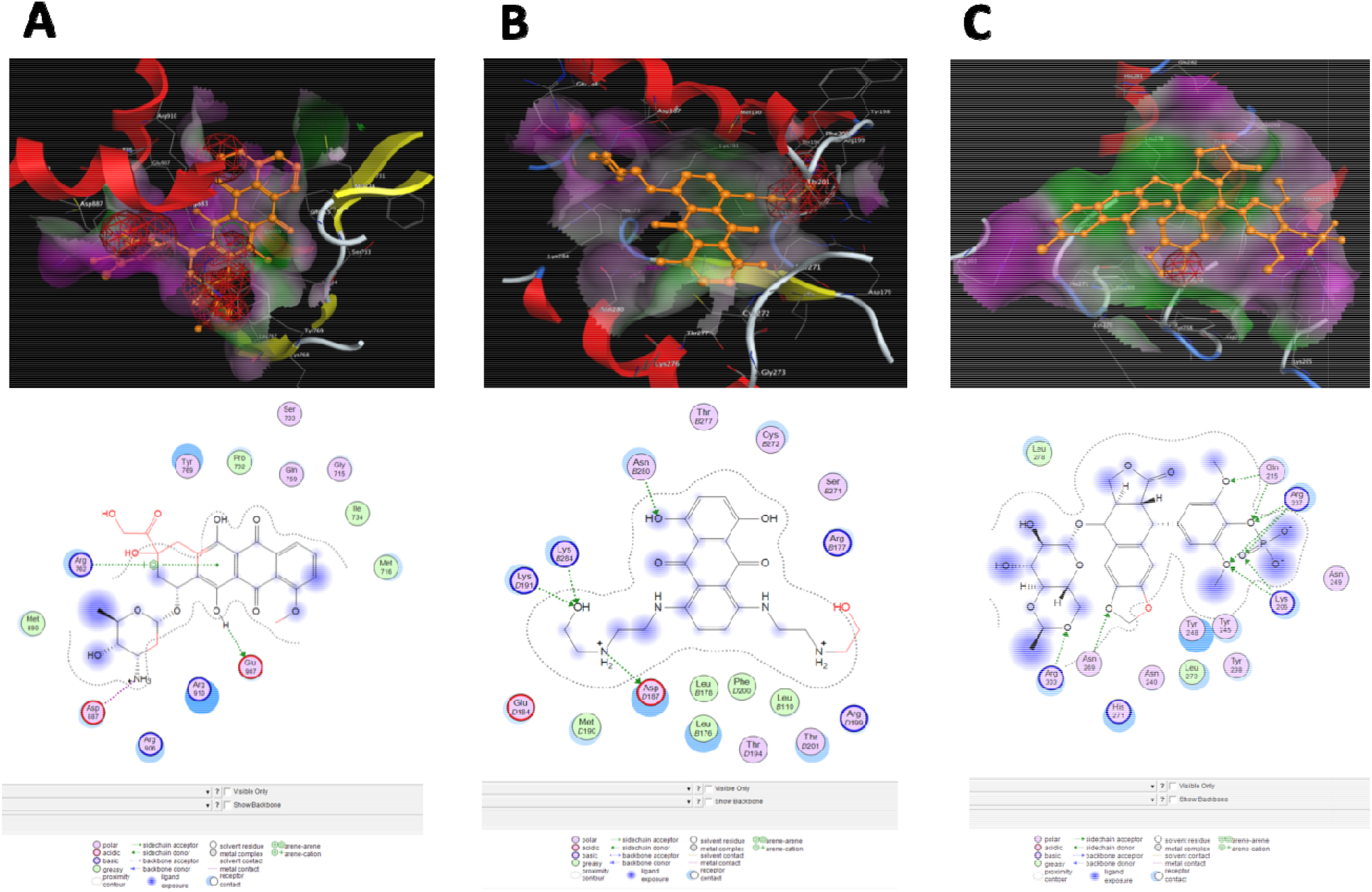
Molecular Docking Analysis of POTE with Anticancer Agents. (A). Binding conformation of Doxorubicin in an active pocket of POTEF protein. **(B)**. Binding conformation of Mitoxantrone in an active pocket of POTEK protein. **(C)**. Binding conformation of Etoposide in an active pocket of POTEM protein.

**Table 3.** Root Mean Square Deviation (RMSD) of both Predicted and Refined Structure.

| S.No. | MODEL ID | Initial Cα RMSD | GalaxyRefine |
| --- | --- | --- | --- |
| 1 | POTEA | 12.28 | 12.008 |
| 2 | POTEB | 4.8 | 4.6 |
| 3 | POTEB2 | 4.8 | 4.6 |
| 4 | POTEB3 | 9.40 | 9.077 |
| 5 | POTEC | 10.3 | 10.163 |
| 6 | POTED | 11.861 | 11.6 |
| 7 | POTEE | 0.554 | 0.41 |
| 8 | POTEF | 0.556 | 0.260 |
| 9 | POTEG | 11.811 | 11.6 |
| 10 | POTEH | 12.357 | 12.01 |
| 11 | POTEI | 0.691 | 0.54 |
| 12 | POTEJ | 11.32 | 11.01 |
| 13 | POTKP | 1.578 | NA |
| 14 | POTEM | 5.89 | 5.32 |

Refined structures also predict helices in the predicted structures, as shown in Fig 6. Refinement of the predicted structures was done first at 10ns to develop an idea about the accessible surface area, fluctuations present in the POTE paralogs’ initial structures, and the number of hydrogen bonds formed by each paralog. The obtained results have been portrayed in Supplemental Figure S5. As the models get compact during the refinement procedure, we thus discern that ROG is reduced after stabilization in all the models, as shown in Supplemental Figure S6. Further, we calculated the number of hydrogen bonds, which subsequently increased, showing that hydrogen bonds’ formation is enhanced across the trajectory shown in Figure S7. By exploring the Accessible Surface Area (ASA), Radius of gyration (ROG), hydrogen bonds, and Energy potential on our preliminary models, stabilized forms of all POTE paralogs were observed. However, after the initial 10ns refinement, it was noted that only POTEE, POTEF, POTEI, and POTEJ have short regions of remodeled gaps, which are highlighted in green Fig 6.

**Figure 6.**
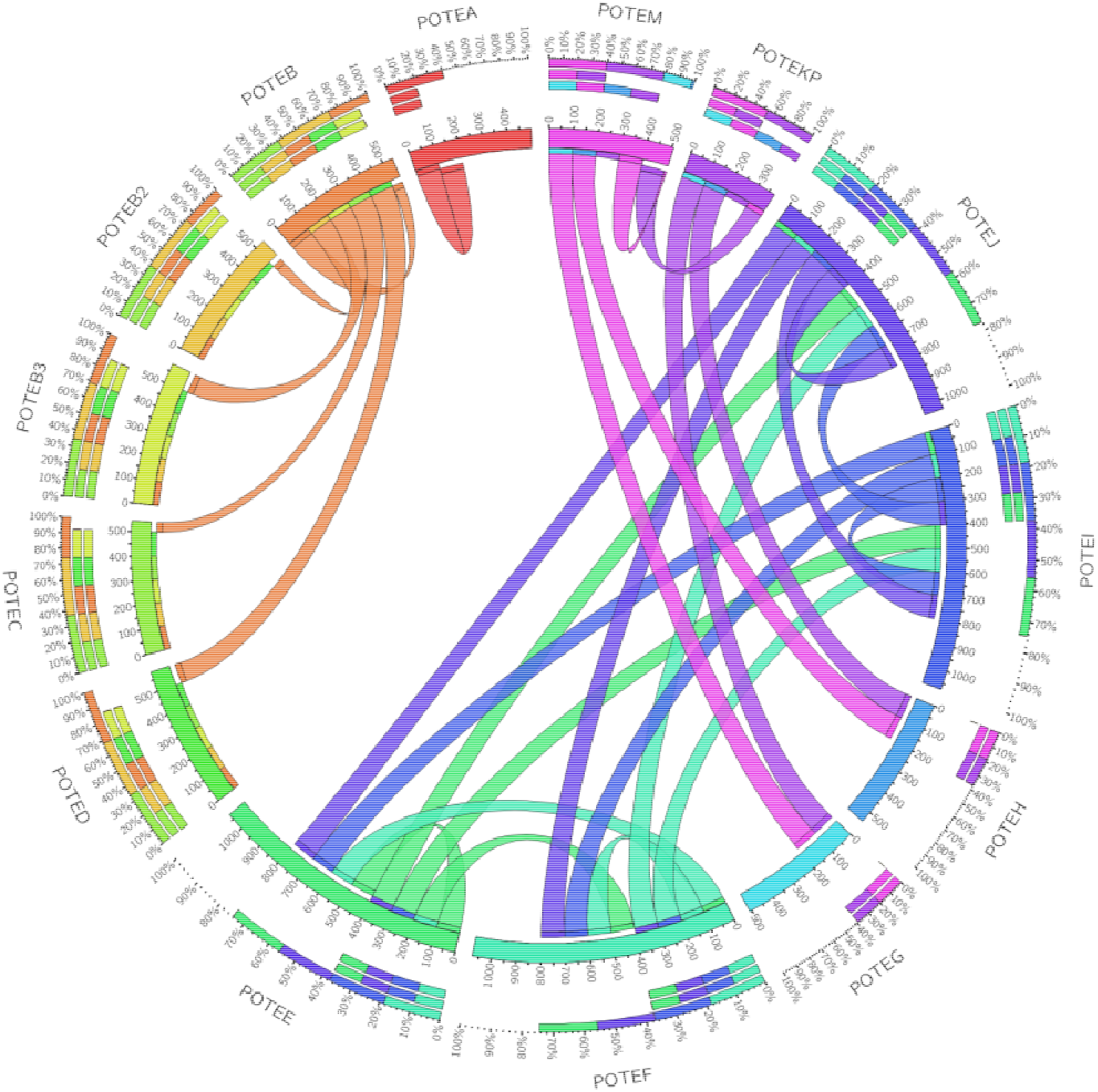
Circos plot: This plot is made using the length of each POTE paralog (mentioned in plot) and the relationship draw on the basis of evolutionary, structural and function, we found that POTE paralog grouped into 4 different groups : POTEA with red color and POTEB, B2, B3, C, D are in orange color and POTEE, F, I, J are shown in blue shades and POTEM, KP, G, H are shown in Purple and pink color.

Furthermore, it was also observed that the ROG of all the four POTEs lay between 40 and 55 cm, and the number of hydrogen bonds formed was less compared to the remaining paralogs. Therefore, we selected only these four POTE paralogs, namely-POTEE, POTEF, POTEI, and POTEJ, for a detailed molecular dynamic simulation analysis executed at 50ns. It is evident that the complexes have been refined to the best potential, and the total energy of the complex has also been stabilized, with all the structures having a good RMSD score with fewer clash scores and are well-fitting the Ramachandran plot criterion. Table 4 summarizes the best refined POTE targets, including preliminary criteria such as - RMSD scores, clash core, accuracy score (refinement of the backbone), Ramachandran plot score, and MolProbity scores. Root mean square deviation (RMSD) describes the various hinges present in the structure during the molecular dynamic simulation (MDS) that comprise the refined structure’s stability and confirms whether the simulation has been equilibrated. The RMSD scores were recorded to be 0.3, indicating a few minor changes after the POTE paralogs’ refinement. The accuracy score defines the improvement of the backbone structure of the initial structure, represents that POTEE (accuracy= 0.9853), POTEF (accuracy= 0.9813), and POTEI (accuracy= 0.9851) have refined better when compared to POTEJ as its accuracy score is only 0.9739. Out of these four, POTEE is much stable as it has fewer steric hindrances and clashes, and accuracy scores (Table 4). The MolProbity score gives the optimal physical correctness of the best refines structure. Typically, MolProbity scores for tertiary structures fall in the range of 1–2 angstrom (A). Our results showcase that paralogs POTEF and POTEJ (MolProbity score= 1.85) structures have good physical correctness compared to the rest.

**Table 4.**
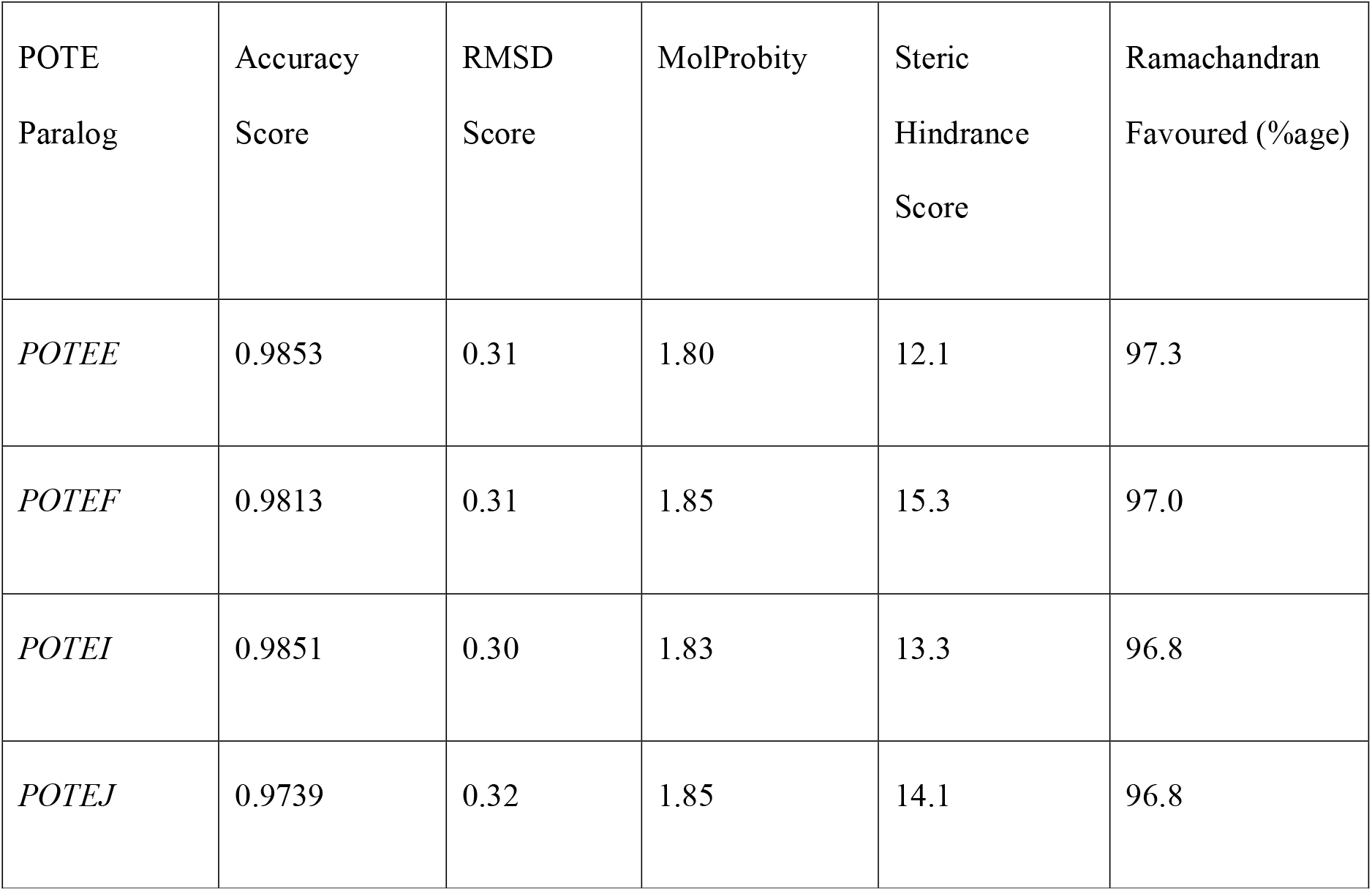
Molecular dynamics simulation (50ns) detailed results of POTEE, POTEF, POTEI and POTEJ.

### Molecular mechanics/generalized Born surface area (MMGBSA) & Electrostatic Computation

Group III POTE paralogs, POTEE, POTEF, POTEI, and POTEJ, were subjected to MM-GBSA and various other essential electrostatic calculations that define the overall stability and energy of the thermodynamic system invariant pH environment. Electrostatics is a crucial factor in understanding how biomolecules interact with one another under various molecular environments. The Adaptive Poisson–Boltzmann Solver (APBS) software was developed to solve the equations of continuum electrostatics for large biomolecule complexes to understand the chemical, biological, and biomedical applications^6^. It was observed that POTEE had an APBS range in between-590.718 to 508.187, POTEF recorded an APBS range in between-565.402 to 503.512, POTEI, on the other hand, had an APBS range in between-575.287 to 504.991, and POTEJ ranged from -561.651 to 499.492. The molecular mechanics generalized Born surface area continuum solvation (MM-GBSA) calculations suggest that POTEE and POTEF are much more robust and electrostatically stable than POTEI and POTEJ. Fig 7 below displays the MM-GBSA calculations in the form of an APBS map as visualized in PyMol software. Table 5 summarizes the necessary electrostatics computations for the four POTE paralogs.

**Table 5.** MMGBSA and other essential electrostatics calculated for POTEE, POTEF, POTEI and POTEJ.

| POTE<br>paralog | System<br>Surface Area<br>(Å<sup>2</sup>) | Generalized Born Self<br>Energy<br>(GBSE)(kJ) | Coulomb<br>Energy (kJ) | Electrostatic<br>Solvation<br>Energy<br>(kJ/mol) | Total<br>Energy<br>(kJ/mol) | APBS Map |
| --- | --- | --- | --- | --- | --- | --- |
| *POTEE* | 3.39090e+04 | -<br>15055.071<br>461 | -97816.405699 | -<br>3693.103361 | -<br>99474.9702<br>52 | -<br>590.718 to 508.<br>187 |
| *POTEF* | 3.32948e+04 | -<br>14831.959<br>332 | -97843.869365 | -3425.638721 | -<br>99271.8188<br>74 | 565.402 to 503.<br>512 |
| *POTEI* | 3.35568e+04 | -<br>14920.890<br>240 | -<br>97415.190820 | -3566.078112 | -<br>98967.8590<br>69 | -<br>575.287 to 504.<br>991 |
| *POTEJ* | 3.30252e+04 | -<br>14695.354<br>653 | -97299.238959 | -3328.072766 | -<br>98645.7973<br>24 | -<br>561.651 to 499.<br>492 |

### POTE protein expression and subcellular localization

Subcellular localization is predicted for all POTE paralogs using various computational tools. It was observed that DeepLoc, MDLoc, and Hum_mPLoc software give approximately the same prediction, but WoLF PSORT predicted different localization. WoLF PSORT software predicts subcellular localization such as the nucleus, mitochondria, cytosol, plasma membrane, extracellular, peroxisome, Golgi, etc., with competitive accuracy but also provides detailed information relevant to protein localization. The data was analyzed, and the result depicted that POTE paralogs are majorly localized in Cytoplasm and Cell membrane, as shown in Supplemental Tables 2, 3, and 4.

### Interaction network

Using the STRING database, we retrieved the interaction between POTE paralogs with a high confidence score, i.e., 0.70, and analyzed that paralogs POTEE, POTEF, POTEI, and POTEJ make one cluster Fig 8. A significant number of genes and POTEs families were connected to functional networks involving cancer-relevant pathways such as cellular growth and proliferation, immune cell trafficking, cell cycle, cancer-related to the reproductive system, etc.

### Function Prediction

ProFunc was employed for predicting POTE paralog’s main functionality. The results are astounding as we discern that POTE paralogs have succumbed themselves according to their nearest neighbor with an attributable course of evolutionary time. Similar structures of paralogs have shared functions. This analysis’s crux is simple, ProFunc assay suggests that all the POTE paralogs are mainly involved in binding like protein, nucleotide, and ATP binding. Moreover, they are also engaged in enzyme regulation, catalytic activity, transporter, and transferase activity, respectively, as shown in Supplemental Table 5.

### Molecular Docking on POTE Proteins against Different Anticancer Drugs

We performed a docking study on 14 POTE paralogs with a list of selected compounds from ovarian, prostate, and testicular anticancer drugs. Doxorubicin showed the highest binding energy of -15.945 kcal/mol for the active site of POTEF among 15 ovarian cancer bioactive compounds, which were taken for the docking study. Notably, one of the quaternary amines of the ligand sites is well in the POTEF receptors active site where the metal interaction with Asp887 is observed, and the hydroxyl establishes an H-bond with Glu907. In contrast, the arene part of the ligand, which has hydroxyl and acidic groups, sits on the middle of the POTEF receptor, where the cation−π interaction with Arg762 is observed Fig 9a.

Mitoxantrone showed the highest binding energy of -15.0988 kcal/mol for the active site of POTEKP. In this protein-ligand interaction, one of the quaternary ammonium ions interacts with the amino acid residue of AspD187 of the POTEKP receptor through acceptor interaction. The hydroxyl group, which is 2 carbon atoms away from the amine group, establishes an H-bond with two amino acid residues Lysb284 and Lysd191, and another hydroxyl group is located on the arene group of the ligand concatenate through H-bond with AsnB280 residue as shown in the Fig 9b. Whereas, the docking results on the testicular anticancer compound, etoposide, showed the highest binding energy of -19.831 kcal/mol for the active site of POTEM among all the three classes of anticancer compounds. The maximum binding interactions between amino acid residues with ligand atoms as Lys205 establish H-bond with the phosphate group’s oxygen atom, and the oxygen atom of the anisole group appended benzene rings. Arg207 shows multiple H-bond interactions with two oxygen atoms of a phosphate group and anisole’s oxygen atom. Simultaneously, the Gln215 interacts through H-bond with two oxygen atoms, one from phosphate and another from the anisole group. The oxygen atom, which is a part of the six-member ring, interacts with POTEM receptors active site where the H-bond interaction is observed with Arg303, and another oxygen atom of the five-member ring establishes an H-bond with Asn269 residue as shown in Fig 9c. Thus, the Etoposide ligand has shown high binding energy with POTEM protein compared with all other anticancer compounds with POTE proteins. A list of bioactive drug compounds from ovarian, prostate, and testicular cancers with the highest binding energy averse to all POTE proteins are shown in Supplemental Tables 6 and 7.

## Discussion

In the post-genomic era, protein sequences, which deposited, have increased at an exponential rate. The latest UniProtKB shows ∼ 99,261,416 protein sequence entries in the repository. However, protein structures present in the protein data bank (PDB) (RCSB) are ∼125,799. The time, labor, and cost involved in the protein structure determinations are augmenting the sequence-structure gap. Structures are essential for function annotation and pursuing structure-based drug discovery^5,7,8^. Computational methods based on ab initio and homology methods can accelerate structure generation and can be used to partly alleviate the dilemma^9^. Hence, we targeted an exhaustive exploratory *in-silico* analysis to know POTE paralogs’ evolutionary status and behavior.

POTE paralogs have had a fate of dilatory rate of evolution, hinting at their high conservation of amino acids. Our study reveals that POTE paralogs are very analogous to one another yet very different. They have gone through an adaptive divergence. Most POTE paralogs diverge from many species, not just primates. POTE proteins are orthologous to many different species such as *D. melanogaster, C. elegans*, fission yeast, *D. discoidum*, yeast, and zebrafish, hinting that the POTE sequences might have undergone an adaptive divergence in the attributable course of evolution. Furthermore, this divergence is not in all the POTE paralogs, but only POTEE, POTEF, POTEI, POTEJ, POTEKP, and POTEA, wherein we get to know that POTEA and POTEKP are outliers and do not fall in any of the common clusters which have been formed in our clustering algorithms. Phylogenetic analysis of the POTE paralogs indicates that POTEE, POTEF, POTEI, POTEJ, POTEA, and POTEKP are clubbed together, hinting at their common divergence from other species during evolution, while POTEB, POTEB2, POTEB3, POTEC, POTED, POTEG, POTEH, POTEM comes from only primates.

The evolutionary analysis, which was done using MEGA 5.02, discerns that all the POTE paralogs’ aggregate evolutionary divergence is 0.66, again referring to its dilatory evolution rate. Most of the residues in POTE protein sequences are conserved, referring to a low rate of alterations/ mutations. Common amino acids present in all the POTE paralogs are mainly: Serine, Arginine, Lysine, and Valine, respectively, which may vary in their composition in each paralog. Structure developments showcase each paralog’s varying structures, but every paralog has a helical and supercoiled structure as its primary protein skeleton. Evaluations of every model were generated to discern that the Swiss Model structures are better than Phyre2 and MODELLER. The refinement provided a successful 0.5 Å of resolution. The graphical study of the radius of gyration, accessible surface area, hydrogen bond, and the energy potential shows every POTE paralog structure’s compactness. The explicit MD simulation, MM-GBSA, and APBS electrostatics suggest that only POTEE and POTEF targets have absolute high affinities with minimal energetic exploitation. Both of these paralogs – POTEE and POTEF are highly stable as a system with equilibrated thermodynamic properties and Generalized Born self-energy (GBSE). The MM-GBSA estimates also indicate that POTEE and POTEF have a higher entropic contribution than POTEI and POTEJ.

We also ascertained that the tertiary structures of POTEE, POTEF, POTEI, POTEJ, and POTEKP (pseudogene) have beta strands along with helices and loops. However, POTEA, POTEB, POTEB2, POTEB3, POTEC, POTED, POTEG, POTEH, and POTEM had only helices. This ardently provides evidence to our evolutionary analysis that some of the POTE paralogs have adaptively diverged and differed in sequence, structure, and functioning compared to their counterparts. Adaptive divergence leads to new forms resulting from the adaptation to a new environmental condition^10,11,12^. Thus, new forms of POTE paralogs have emerged with time from ancestral origins, giving rise to more robust gene and protein structures. We would also like to correlate our study with^5,13,14^, where the authors suggest that group 3 POTE members are not cancer-testis antigens functioning only in normal tissues. This differing nature of group 3 POTE paralogs is for sure to ponder upon. Our study also suggests that group 3 POTE paralogs are very ardently compared to the rest of the members. Henceforth, we propose that this differing nature of group 3 POTE members POTEE, POTEF, POTEI, POTEJ, and POTEKP (non-functional) is only their adaptive divergence.

Further, we have identified the sub-cellular localization of POTE paralogs, as predicted by WoLF PSORT, Hum_mPLoc, DeepLoc, discerns that POTEE, POTEF, POTEI, POTEJ, POTEKP, and POTEM are localized in the cytoplasm, while others are located in the extracellular region. Our study on interaction network analysis discerns that POTEE, POTEF, POTEI, and POTEJ have a high confidence score of 0.70 with only one interactor protein, yet again corroborating our phylogenetic studies, which attributable to the advent of subdivisions in already existing group 3.

Function prediction and its analysis suggest that all the POTE paralogs, which are predicted and refined, are mainly involved in biological functions such as protein, nucleotide, and ATP binding. Moreover, they are also engaged in enzyme regulation, catalytic activity, transporter, and transferase activity. POTE family encodes six or seven ankyrin repeats in the middle of the molecule, spectrin-like structure at the carboxy-terminus, and three cysteine-rich repeats at the amino-terminus. Ankyrin repeats have been identified in numerous functionally diverse proteins and are involved in protein-protein interactions in many functional pathways within the cell ^15,16^. In accord, POTE expression in the testis is primarily confined to spermatids, which are caspase-3 positive^17^. Ingenuity pathway analysis of both microarray and RNA-Seq data also suggests that POTEs might be associated with cancer pathogenesis and connected to functional networks involving cancer-relevant pathways such as cellular growth and proliferation.

Further, the molecular docking results show a higher affinity of POTE members for different anticancer drugs. Etoposide and doxorubicin showed the highest binding energy of -19.831 kcal/mol and -15.0988 kcal/mol, respectively, for the active site of POTEM and POTEF proteins. However, the results are preliminary and surely need experimental confirmation, which will be conducted soon via molecular biology studies. However, contemplating all these structural aspects and Glide score, the POTE family might be the first choice that could be exploited to design as an anticancer therapy in the future. To correlate our research findings, we designed a circos plot that encapsulates all the stated analogies stated in this paper. The circus plot has been represented in Fig 10.

Our study reveals that POTE paralogs have gone through an adaptive divergence. Most of the POTE paralogs diverge from many species, and not just primates. The evolutionary analysis discerns that all the POTE paralogs’ total evolutionary divergence is 0.66, again referring to its dilatory evolution rate. POTE paralogs are mainly involved in binding like protein, nucleotide, and ATP binding. Moreover, they are also engaged in enzyme regulation, catalytic activity, transporter, and transferase activity. Of interest, cytoplasmic/nuclear/membrane-localized POTEs are more evident in advanced cancer cases. Taken together, further investigations are required to understand better the roles of POTEs members in cancer progression and malignancy and to determine how dysregulation of these genes/proteins may alter responses to therapeutic. Using virtual screening methodology, this study may also help understand and design the novel anticancer compounds concerning POTE proteins.

## Materials & Methods Data Sources

All the 14 POTE paralogs members have been considered, i.e. POTEA (NM_001002920; Q6S8J7), POTEB (NM_001277304; A0A0A6YYL3), POTEB2 (NM_001277303; H3BUK9), POTEB3 (NM_207355; A0JP26), POTEC (NM_001137671; B2RU33), POTED (NM_174981; Q86YR6), POTEE (NM_001083538; Q6S8J3), POTEF (NM_001099771; A5A3E0), POTEG (NM_001005356; Q6S5H5), POTEH (NM_001136213; Q6S545), POTEI (NM_001277406; P0CG38); POTEJ (NM_001277083; P0CG39), POTEM (NM_001145442; A6NI47) and POTEKP (AY014272; Q9BYX7) respectively. The above protein paralogs FASTA sequences were retrieved from HUGO Gene Nomenclature Committee (HGNC)^18^.

### Sequence Similarity & Alignment

As the protein sequence data of these paralogs is quite large and highly divergent, we have executed our study with leaves of 14 sequences (POTEA, POTEB, POTEB2, POTEB3, POTEC, POTED, POTEE, POTEF, POTEG, POTEH, POTEI, POTEJ, POTEM, and POTEKP) could keep genetic divergence small. The UniProt (https://www.uniprot.org/) accession numbers of the 14 POTE member sequences are shown in **Table 1**. For deducing the sequence similarity of the paralogs, we employed bioinformatics tools such as Simple Modular Architecture Research Tool – Basic Local Alignment Search Tool (SMART BLAST) (https://blast.ncbi.nlm.nih.gov/smartblast/smartBlast.cgi?CMD=Web), which tends to provide a breviloquent graphical summary of the entire proteins based on their evolutionary tracts^19^ and Position-Specific Iterated PSI-BLAST (https://www.ebi.ac.UK/Tools/sss/psiblast/) iteratively searches many protein databases like NR (non-redundant) and utilizes a profile Position-Specific Scoring Matrix (PSSM) for the same^20^. After deriving the sequence similarity, we moved to a sequence alignment of the POTE paralogs that were executed based on the multiple sequence alignment (MSA) analogy^21^. Multiple sequence alignments (MSA) (https://www.ebi.ac.uk/Tools/msa/) are pivotal in various sequence analysis methods and are generally calculated using heuristic methods. The protein sequences were aligned by using CLUSTALW^22^. Multiple Sequence Comparison by Log Expectation (MUSCLE)^23^ and CLUSTAL Omega ^24^ with default settings.

### Phylogenetic Analysis

The phylogeny of POTE protein sequences was developed based on the multiple alignments of amino acid by employing Un-weighted Pair Group Method with Arithmetic Mean (UPGMA), Neighbour Joining (NJ), Minimum Evolution (ME), Maximum Parsimony (MP)^21^, and Maximum Likelihood (ML) in MEGA 5.2^25^. UPGMA is an agglomerative clustering method that shows the phenotypic similarities between operational taxonomic units (OTU) by showing an ancestral root. It assumes that evolution rates are more or less constant among different lineages^26^. Neighbor-joining (NJ) attempts to correct the UPGMA method for its inappropriate assumption about constant evolutionary rates throughout the lineage; thus, it gives a rootless phylogenetic tree. NJ is similar to the UPGMA method because the distant pairs of nodes are linked, and their common ancestral node is added to the tree, and their nodes are pruned from the tree accordingly^27^. Minimum evolution (ME) is a distance-based phylogenetic method where the trees are calculated from the pair-wise distances between the sequences rather than from the fit of individual nucleotide sites to a tree^28^. The trio, UPGMA, NJ, and ME are distance-based methods of the phylogeny.

Maximum Parsimony (MP) and Maximum Likelihood are two cladistics methods of generating phylogenetic trees for a commonly set species or reproductively isolated populations of a single species. Maximum parsimony (MP) searches for a tree that needs only a few evolutionary changes to explain the differences observed among the OTUs (Biology 1971). Maximum Likelihood (ML) creates all possible trees containing the set of organisms considered and then uses the statistics to evaluate the most likely tree for a small number of populations^29,30^.

Furthermore, we have also discerned each of the POTE paralogs’ amino acid compositions and the number of amino acid substitutions in each paralogue. Some informative, conserved, and Mark-Parsim sites have also been identified, which can help POTE family members’ structural analysis. The phylogenetic analysis also led to the estimation of POTE’s average evolutionary divergence rate, which is essential in deducing its motion of divergence. The disparity index test was also determined using Monte Carlo replications. Supplemental Figure S1 represents the computational and wet lab workflow used to execute this research.

### Secondary Structure Prediction

Secondary structures of all the POTE Paralogs were predicted and produced using PSIPRED software. To know the helix, Beta, and loop exact position of amino acid of all POTE paralogs, we had made the PSI-blast-based secondary structure Prediction (PSIPRED) (http://bioinf.cs.ucl.ac.uk/psipred/)^31^.

### Homology and Threading Strategies for Structure Prediction

Structures of all the POTE paralogs were produced using homology and protein threading approaches of B.I. software such as the Swiss Model, MODELLER, and Phyre2^32,33,34^ respectively. Swiss Model and MODELLER are based on the sequence homology of the proteins, which are template-based. Phyre2 is a remote homology recognition threading strategy that employs Hidden Markov Models (HMMs) or only profiles to build precise tertiary structures of proteins.

### Quality Assessment of Protein models

The retrieved protein models were then subjected to evaluations using Protein Quality (ProQ), Protein Structure Analysis (ProSA), and RAMPAGE^35,36^. This was done to get the best optimal and stable tertiary structures, which can be further analyzed using MD simulations.

### Molecular Dynamic Simulations (MDS)

We first executed a refinement analysis for all the 14 POTE paralogs at 10ns using the GalaxyRefine^37^ online tool wherein the paralogs are stabilized using the AMBER force field ff94^38^. After retrieving the refinement results, based on the ROG and RMSF results, we selected group III POTE paralogs-POTEE, POTEF, POTEI, and POTEJ for a detailed molecular dynamic simulation (MDS) using QwikMD toolkit^39^ of Visual molecular dynamics (VMD)^40^. Generalized Born Molecular Mechanics (GBMM) was deployed to retrieve the approximate results in explicit solvent. NVT dynamics were deployed, which hold an amount of substance (N), volume (V), and temperature (T) constants. The Noose-hover temperature was set to 300 K, and the entire simulation was executed at 50 ns. The visualization of the refined structures was done in PyMol (https://pymol.org/2/).

### Molecular mechanics/generalized Born surface area (MMGBSA) & Electrostatics Computation

The Molecular Mechanics-/generalized Born surface area (MM-GBSA) approach was also deployed to estimate the binding free energy (delta G) for complexes over simulation time^41^. This was executed using the APBS plugin available in VMD software (https://pymolwiki.org/index.php/APBS_Electrostatics_Plugin). Electrostatics were computed using Blues software^42^.

### Sub-cellular localization analysis of POTE proteins

To study POTE paralogs’ functions, it is essential to know its subcellular localization as a protein can be localized either in the outer membrane, inner membrane, periplasm, extracellular space, or cytoplasm. Hence, before proceeding with the functional analysis, we checked for the sub-cellular localization using four different software, i.e., WoLFPSORT^43^, Hum_mPLoc 3.0^44^, DeepLoc^45^, and MDLoc^46^. WoLF PSORT software uses amino acid sequence and some sorting signal motifs of targeted protein to predict its subcellular localization of the protein. It displays information about detected sorting signals. Hum_mPloc 3.0 is also based on amino acid sequence and predicts 12 human subcellular localizations. DeepLoc software predicts the subcellular localization based on a neural network that processes the entire protein sequence and an attention mechanism identifying protein regions important for the subcellular localization. This online resource can differentiate between different localizations: Nucleus, Cytoplasm, Extracellular, Mitochondrion, Cell membrane, Endoplasmic reticulum, Golgi apparatus, Lysosome/Vacuole, and Peroxisome. MDLoc predicts the multiple locations for proteins using inter-dependencies among locations. It is based on an iterative process and uses the DBMLoc dataset for predicting subcellular localization. We give POTE protein sequences input in all the software, which is retrieved from the UNIPROT database.

### Functional Prediction using ProFunc

Functional prediction and analysis were executed by employing ProFunc^47^ on the optimal selected tertiary structures on both predicted and refined models obtained after an exhaustive validation. ProFunc provides insight into proteins’ functional capacity and various other aspects such as domain and clefts of the proteins, binding capacity, biological, cellular, and metabolic processes.

### Protein-Protein Interaction (PPI) Network

Further, we tried to assess the importance of the most highly connected proteins. For this, we analyzed their clusters using the bioinformatics database STRING^13^ were constructed, and thus, protein interaction networks were retrieved. This database derives high throughput experimental data (>=0.700) from a wide range of sources, analyses the co-expression of genes computationally, uses a scoring framework, and outputs a single confidence score per prediction. This confidence score is a measure of the predicted interactions’ reliability, and a high score indicates that the predicted interactions are also replicated in the KEGG database^48^.

### Molecular Docking on POTE Proteins against Different Anticancer Drugs

All the molecular docking study’s computational procedures were carried out with the Molecular Operating Environment (MOE). POTE protein receptors were initially prepared with the default 3D protonation procedure in MOE^43^. The drug compounds were downloaded from NCI (https://www.cancer.gov/about-cancer/treatment/drugs#D) and then converted from name to 2D structure using ChemAxon tool ((http://www.chemaxon.com) followed by 3D structure generation. Docking was performed using all default parameters with Triangle Matcher, Rigid Receptor, initial scoring method London dG retaining 30 poses, and final scoring method used was GBVI/WSA with 5 poses. POTE protein structures were imported into MOE after removing water molecules. All hydrogen atoms were then added to the structure with their standard geometry, followed by their energy minimization using default parameters with Forcefield value Amber 10: EHT and RMS gradient of 0.1[kcal/mol. Each POTE receptor’s binding site was identified through the MOE Site Finder program, which uses a geometric approach to calculate putative binding sites in a protein, starting from its tridimensional structure. Active sites were identified, and dummy atoms were created around the resulting alpha sphere centers. The backbone and residues were kept fixed, and energy minimization was performed.

## Acknowledgments

I thank Prof. Adam R Karpf, Professor, Fred & Pamela Buffett Cancer Center, University of Nebraska Medical Center, the USA, for his constrictive scientific input. SQ thanks to DST-INSPIRE for fellowship. Thanks to Neetu for her contribution in project work. SQ & BJ equally contributed.

