## Supplemental Figure for "BESFA: Bioinformatics based Evolutionary, Structural & Functional Analysis of Prostrate, Placenta, Ovary, Testis, and Embryo (POTE) Paralogs"

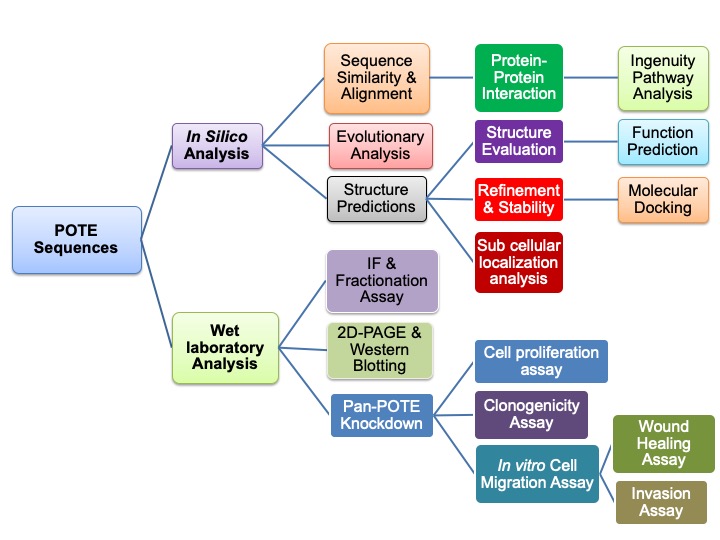

**Supplemental Figure S1: Summary of the computational and wet lab workflow**

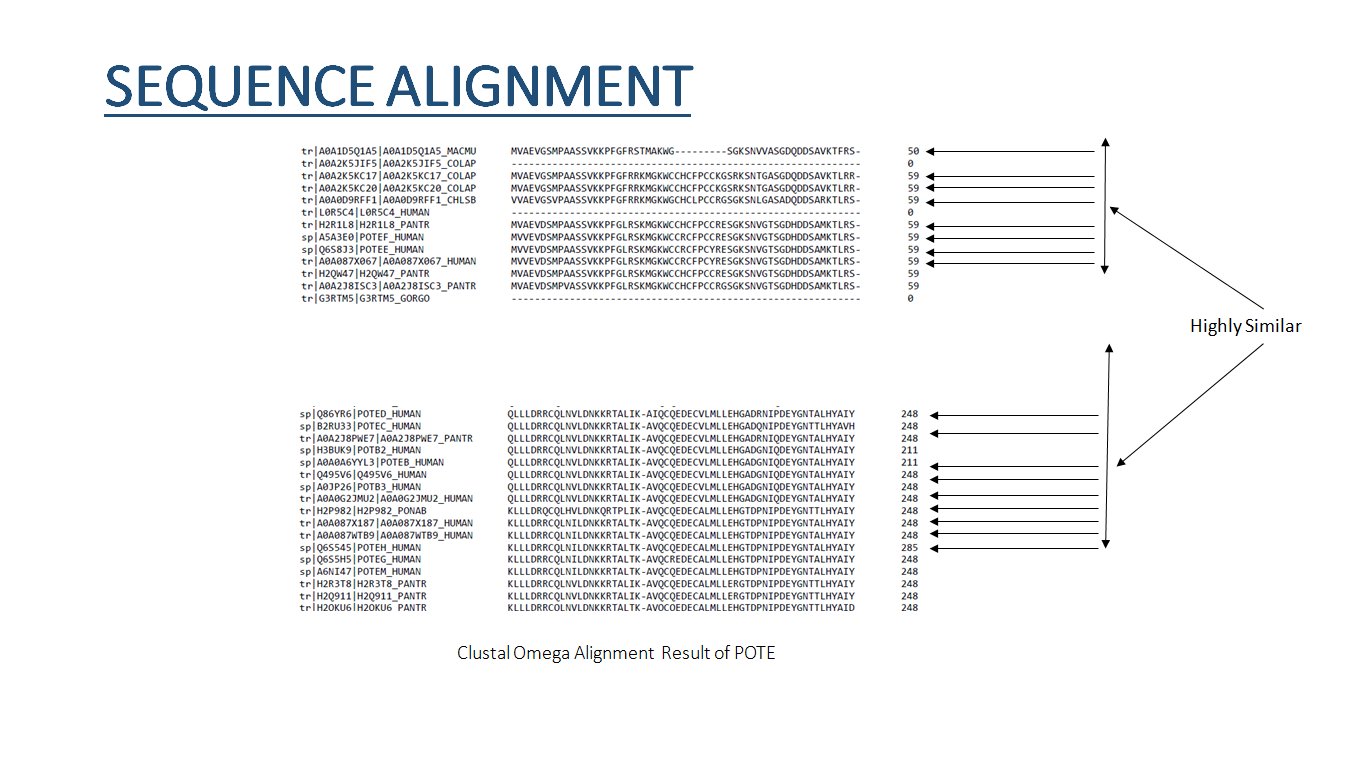

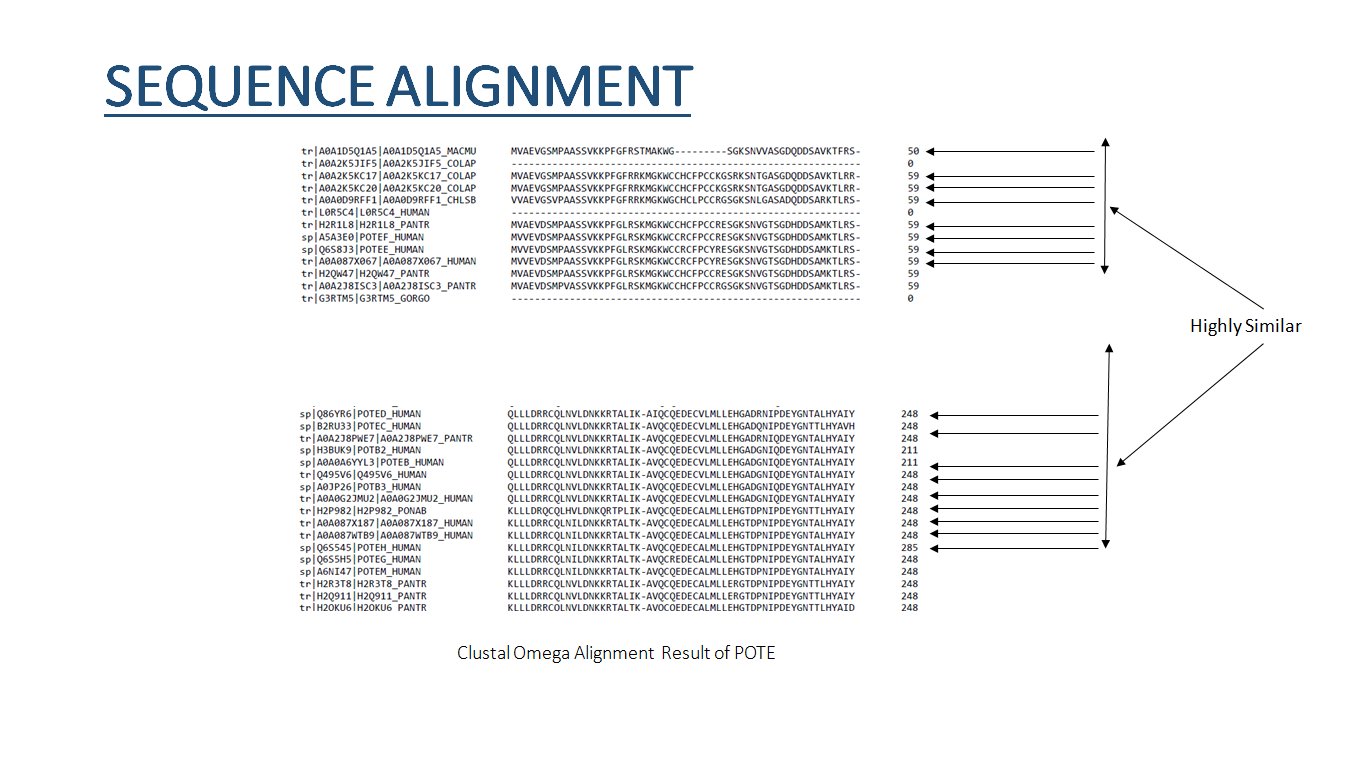

**(a)**

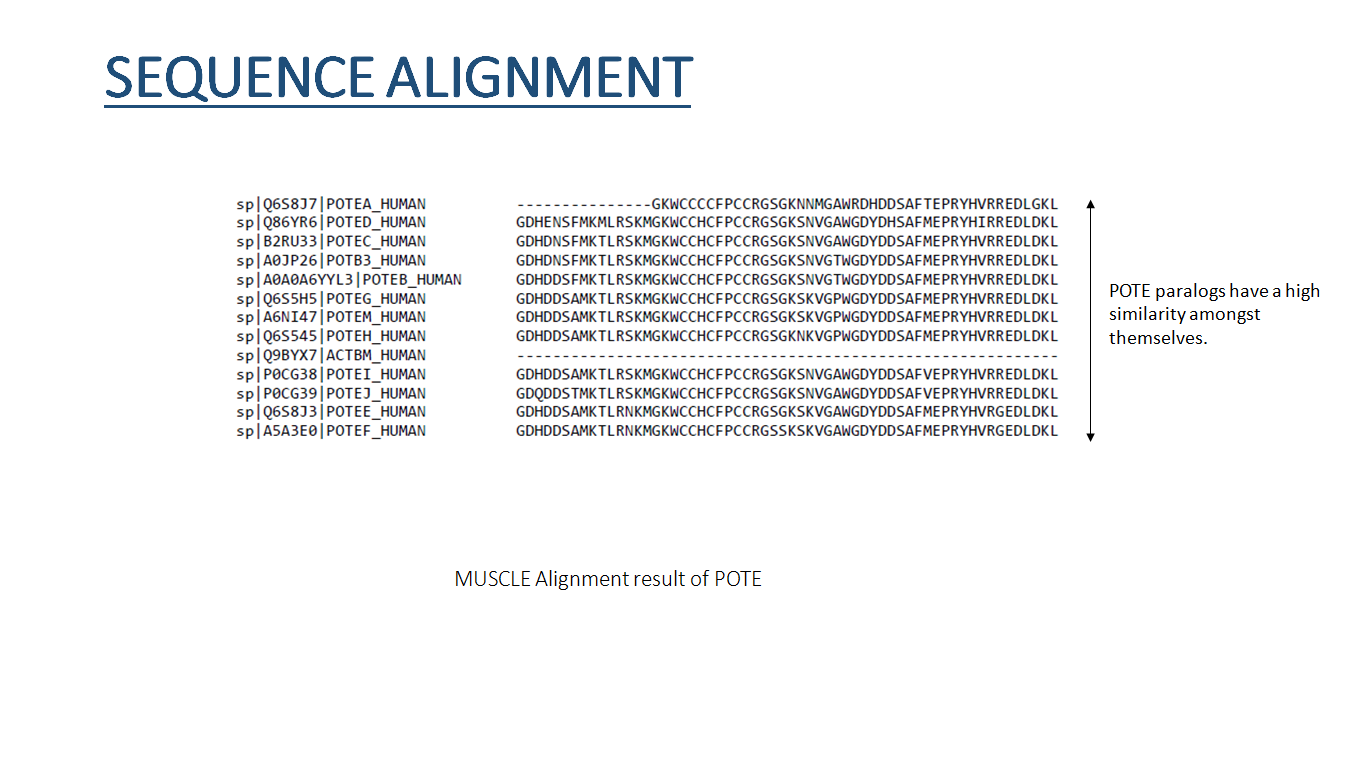

**(b)**

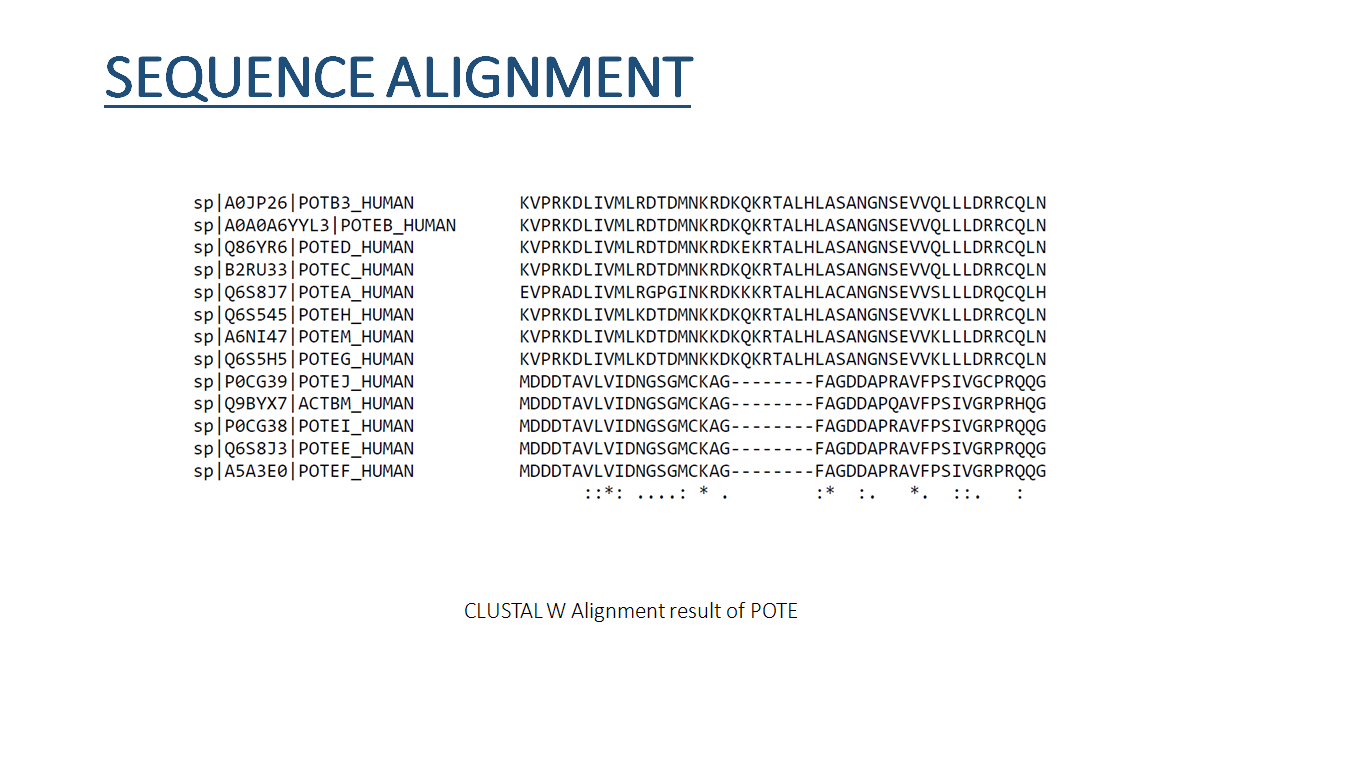

**(c)**

**Supplemental Figure S2: Multiple sequence alignment of POTE paralogs by a) CLUSTAL Omega, b) MUSCLE and c) CLUSTAL W.**

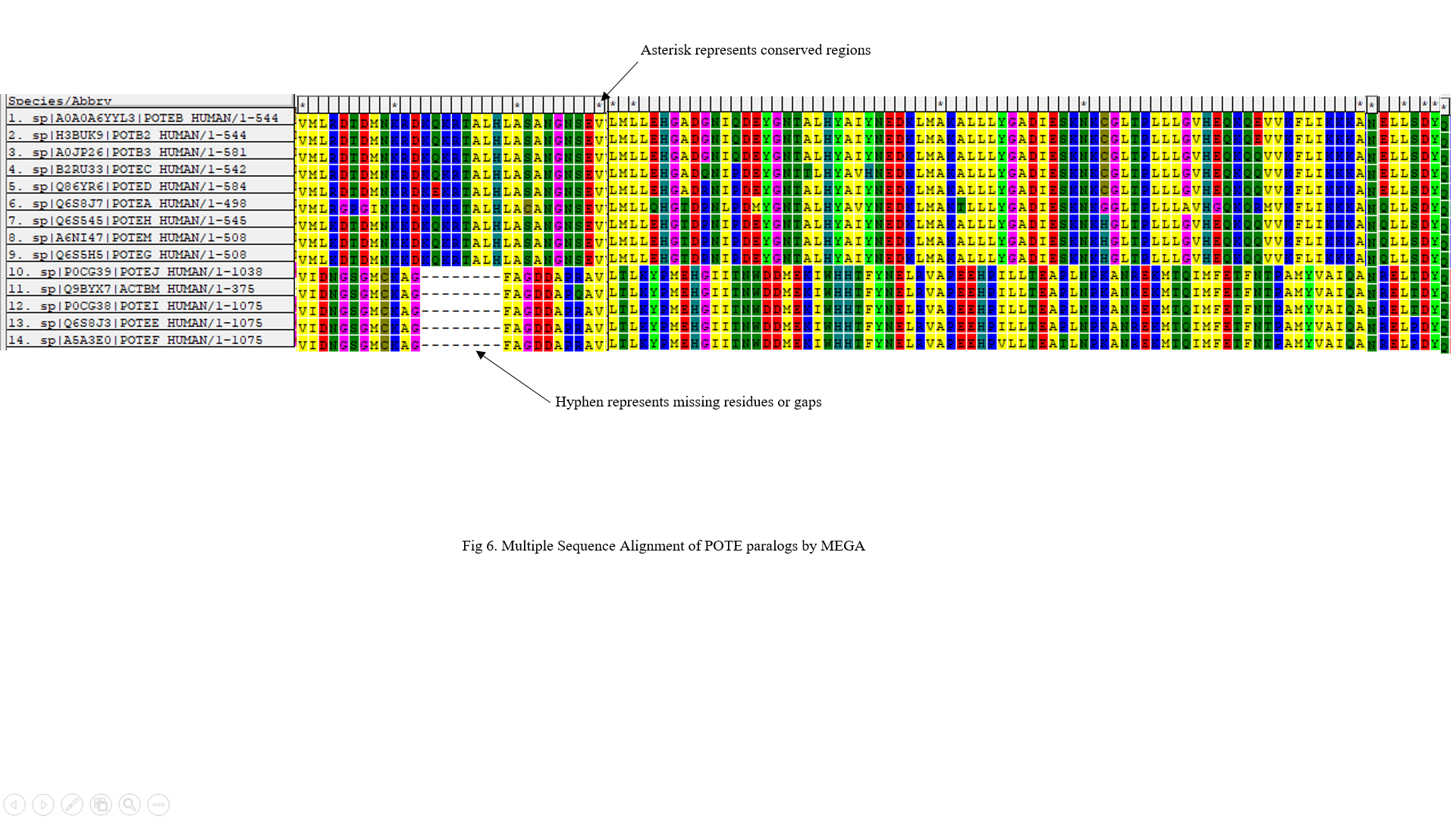

**(a)**

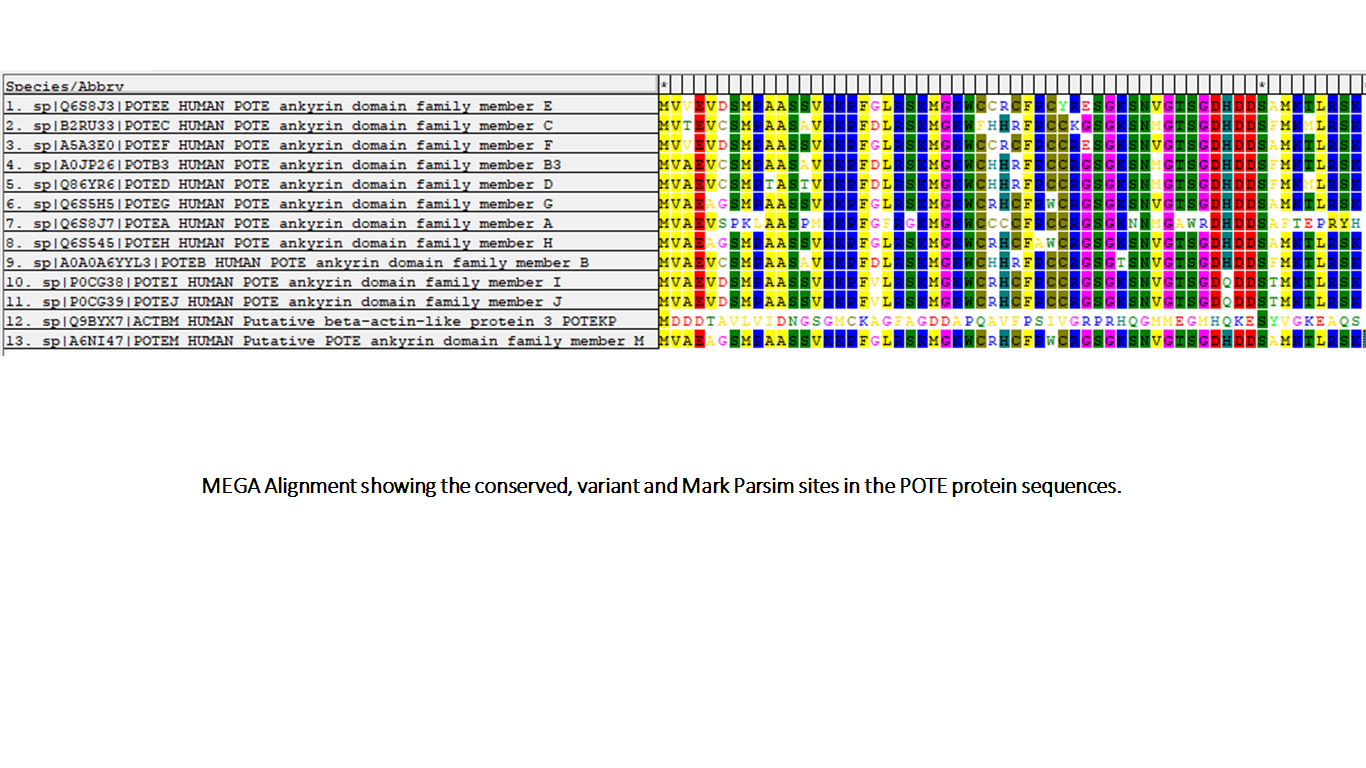

**(b)**

**Supplemental Figure S3: (a) Multiple sequence alignment of the POTE paralogs, (b) Conserved and Mark Parsim Sites discerned using MEGA.**

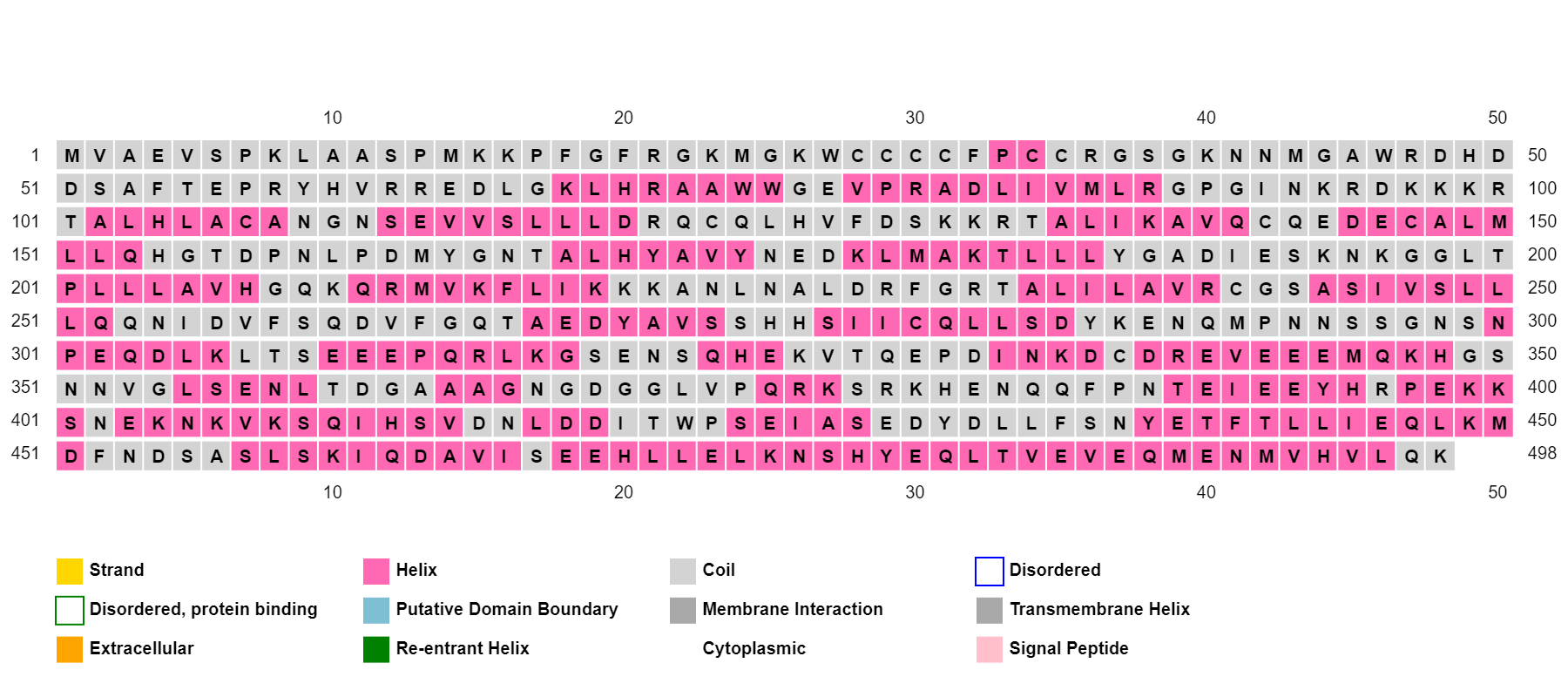

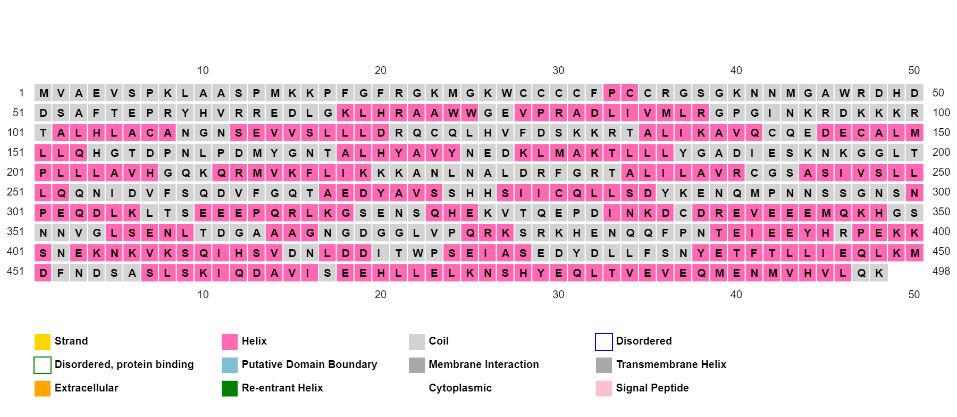
**POTEA**
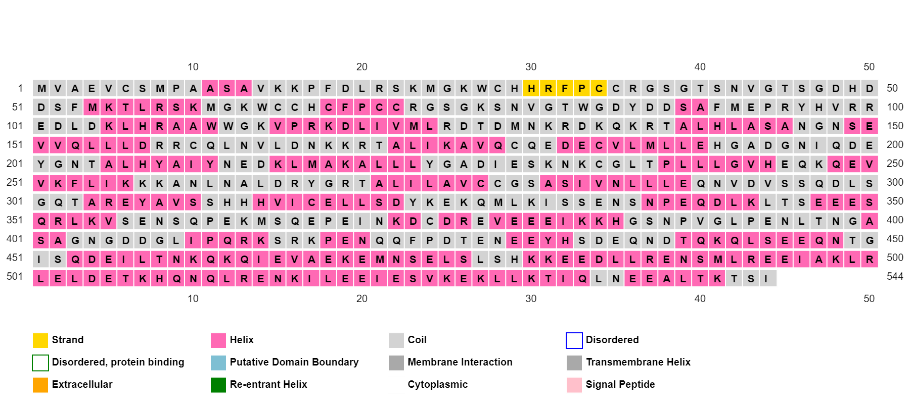

**POTEB**

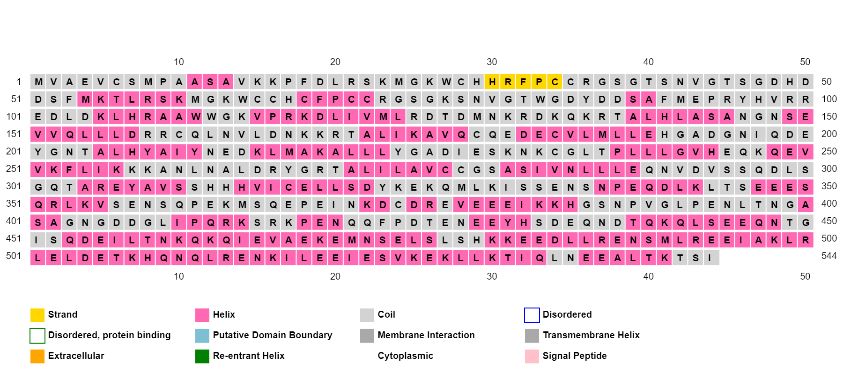

**POTEB2**

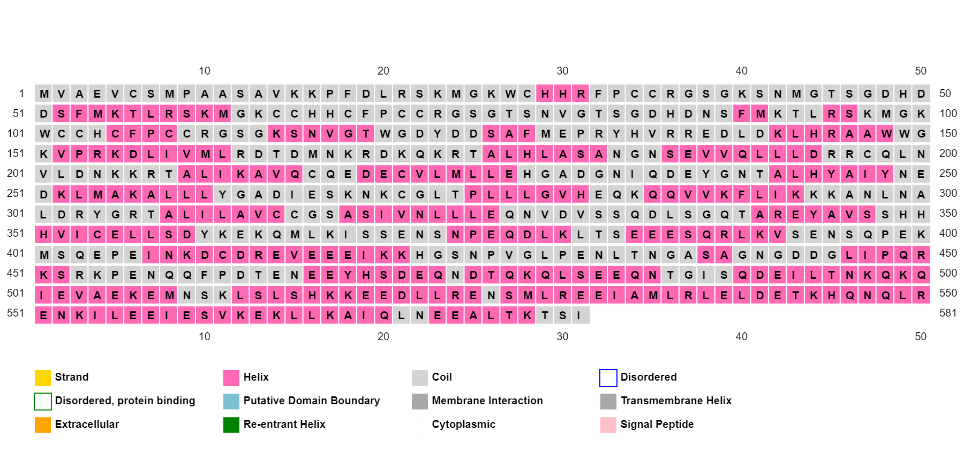

**POTEB3**

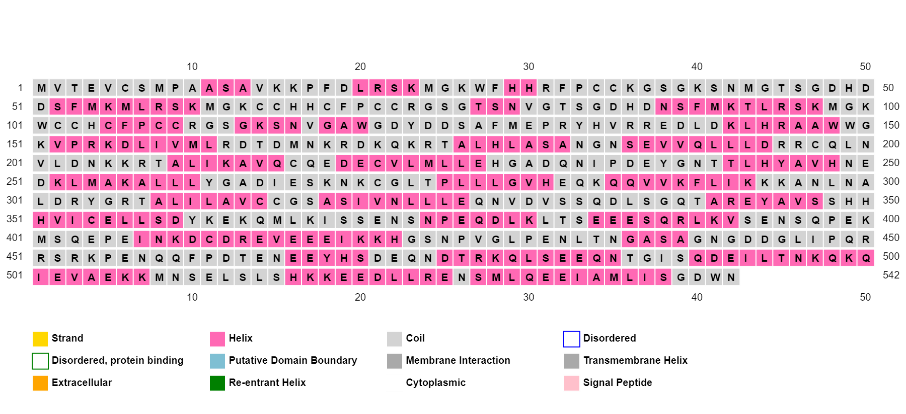

**POTEC**

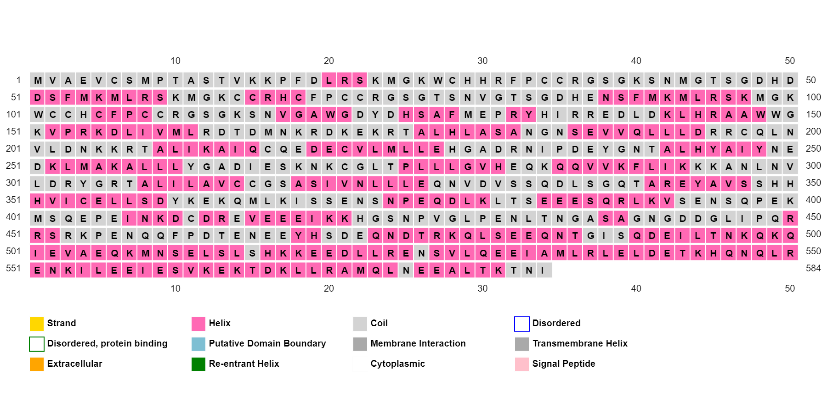

**POTED**

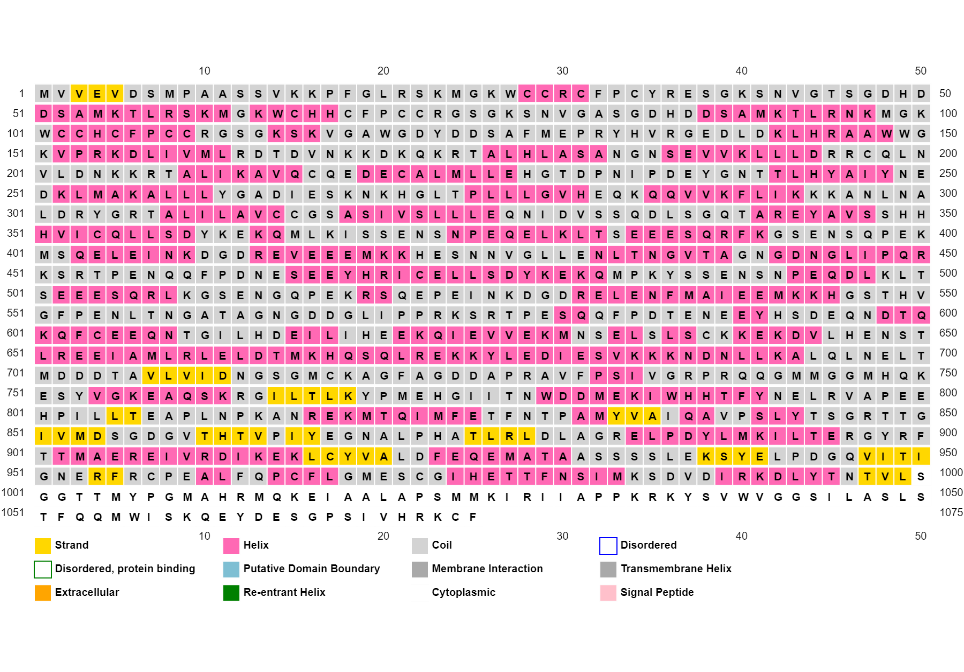

**POTEE**

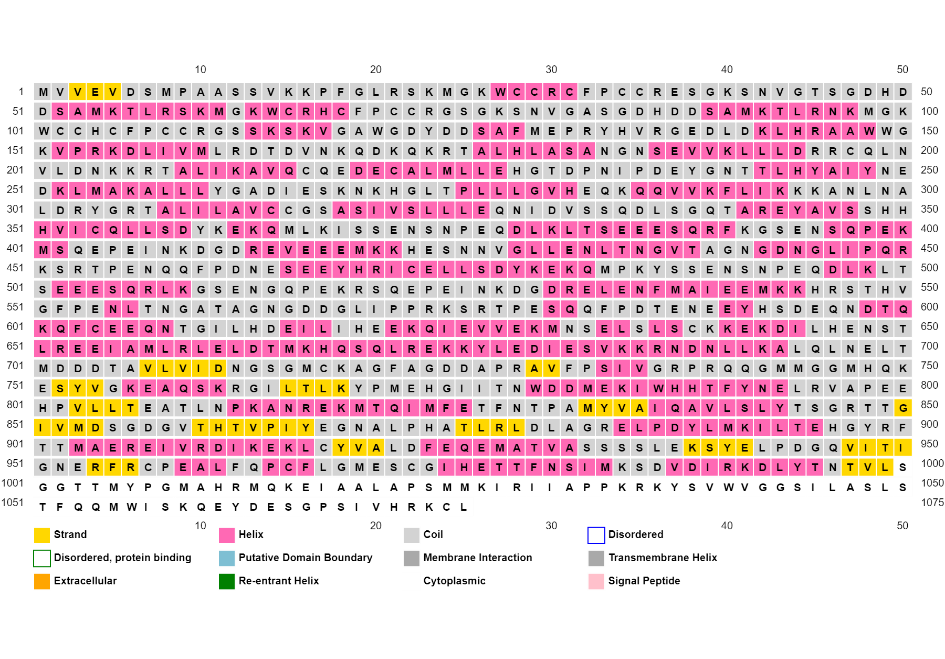

**POTEF**

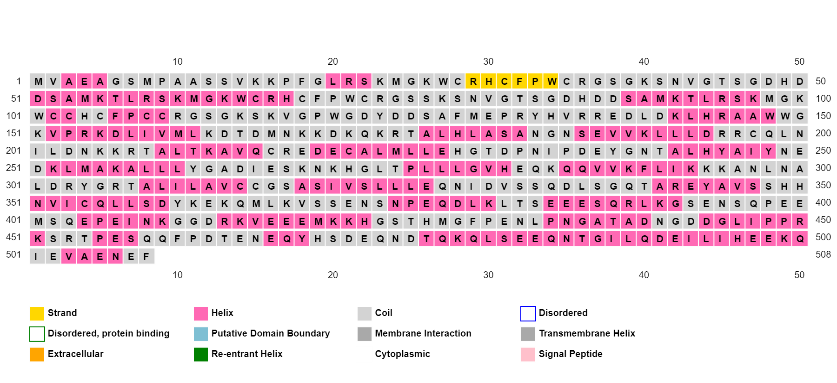

**POTEG**

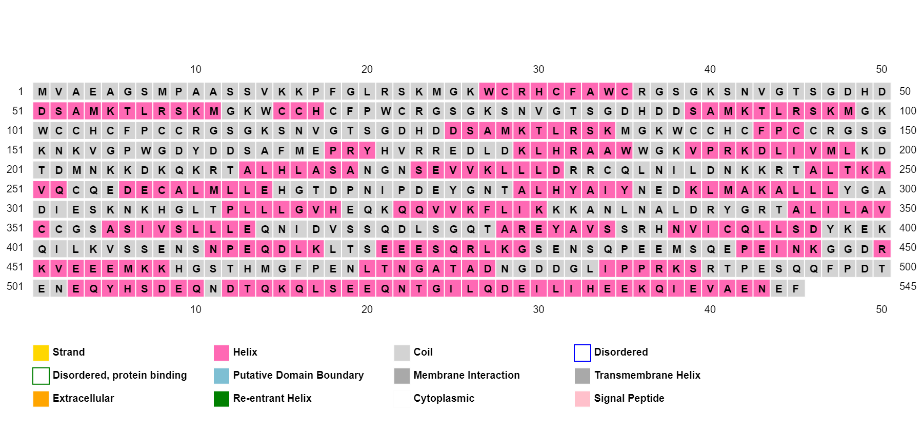

**POTEH**

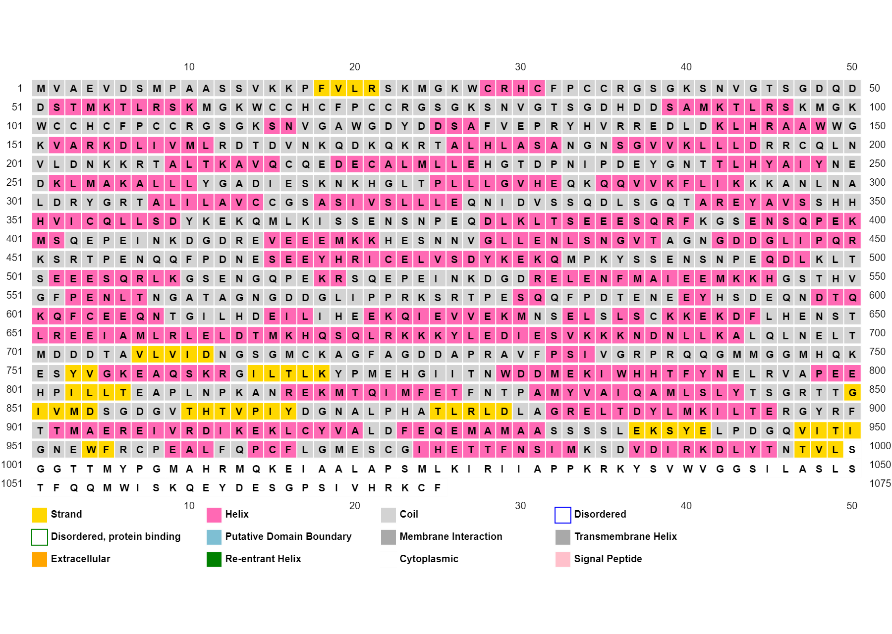

**POTEI**

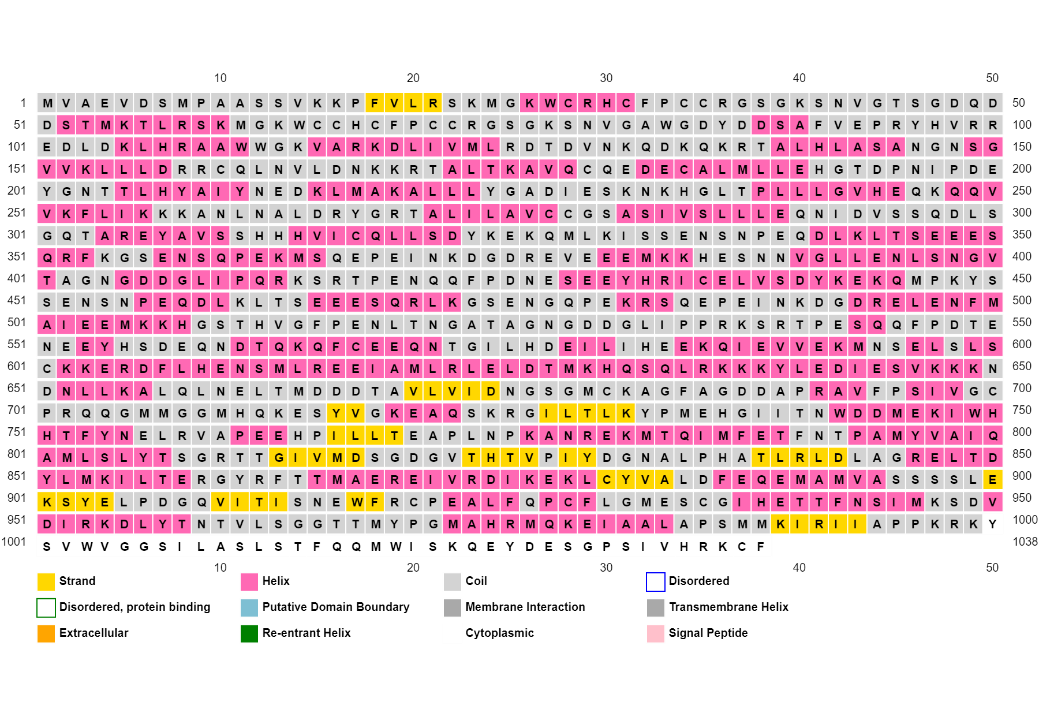

**POTEJ**

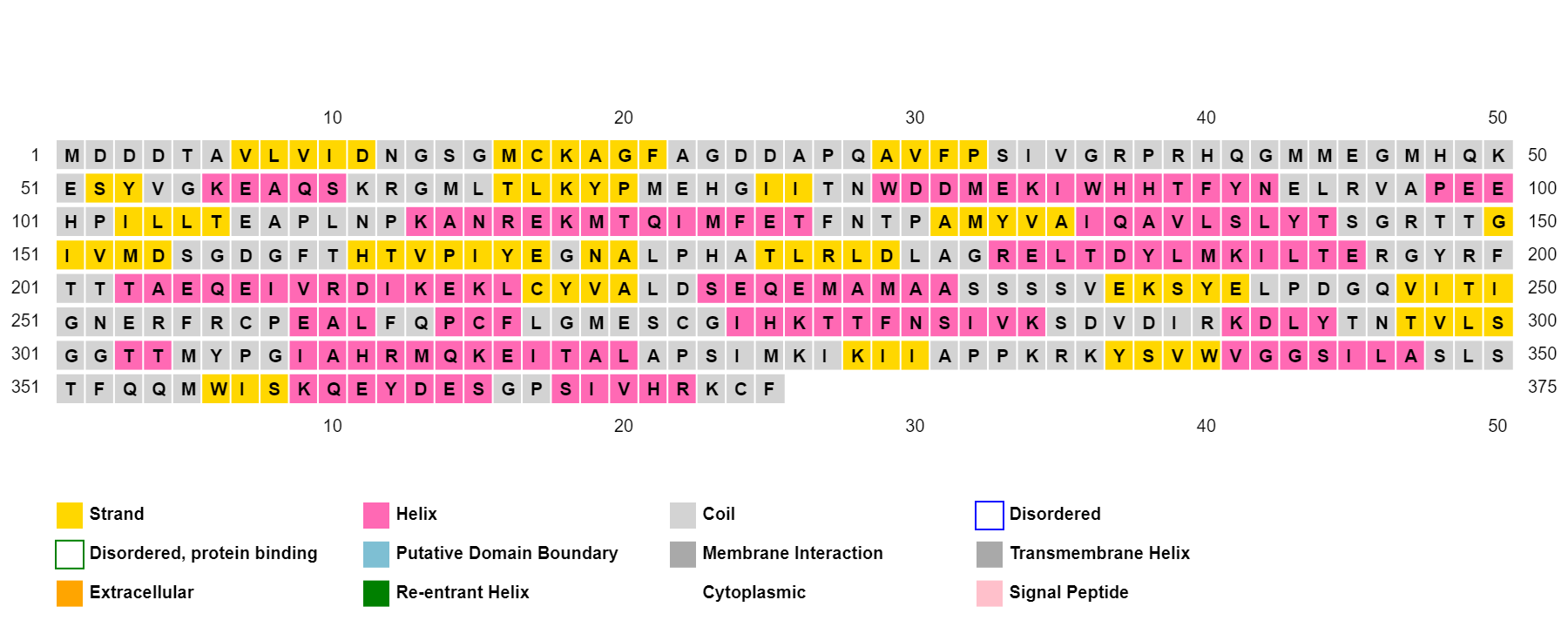

**POTEKP**

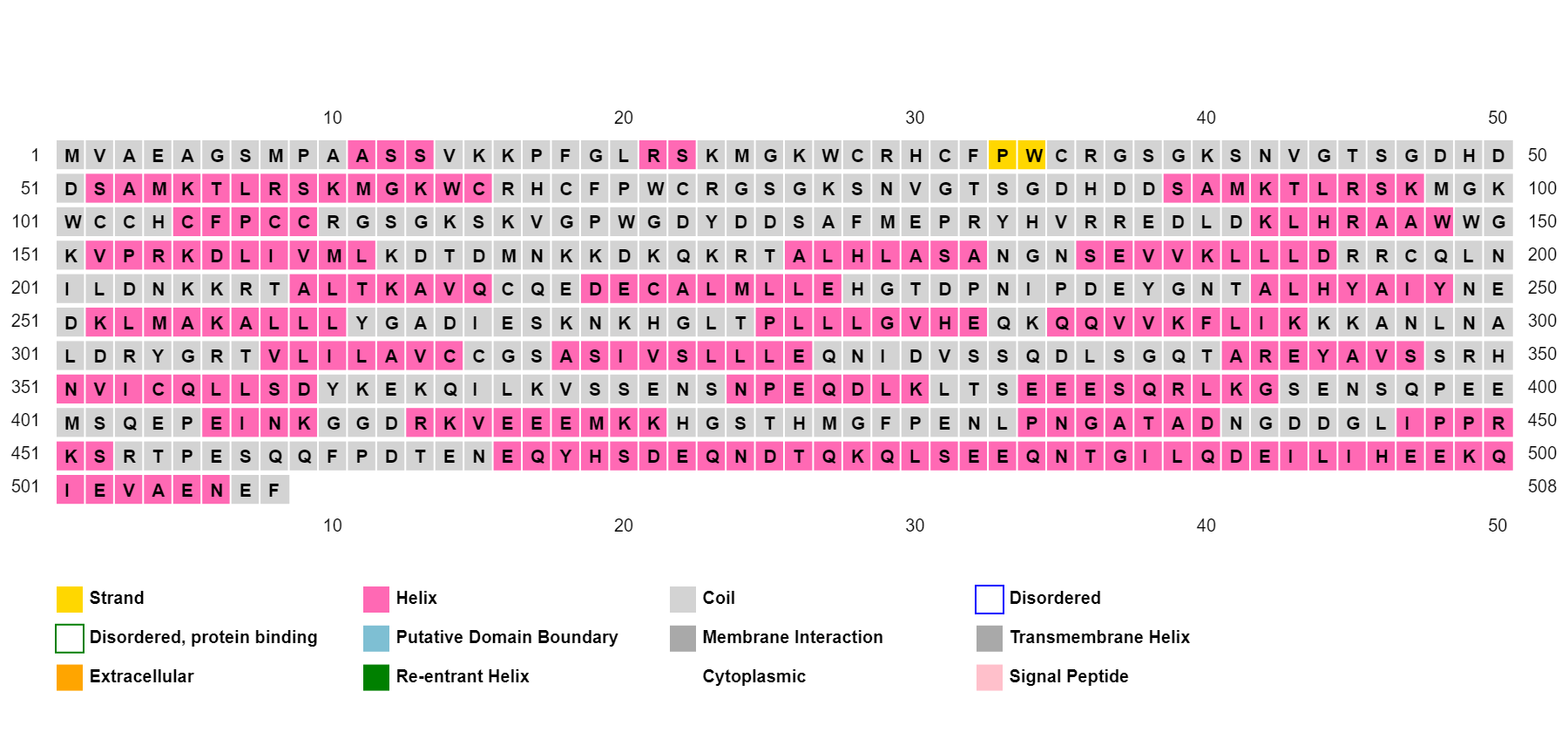

**POTEM**

**Supplemental Figure S4: Secondary structure predicted using PSIPRED of all POTE paralogs respectively.**

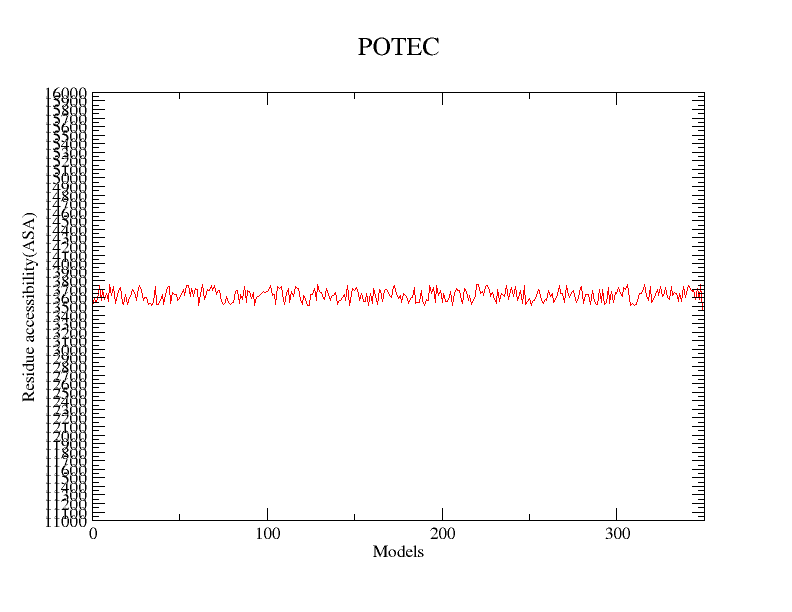

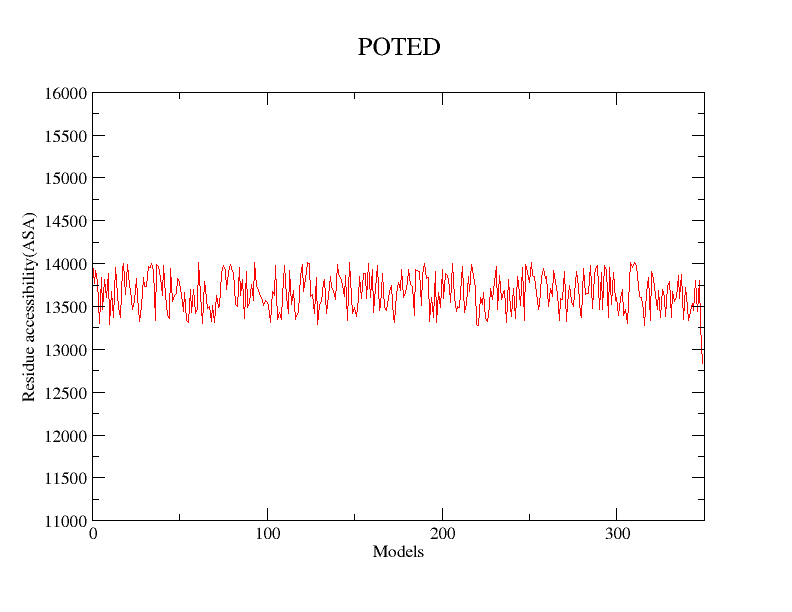

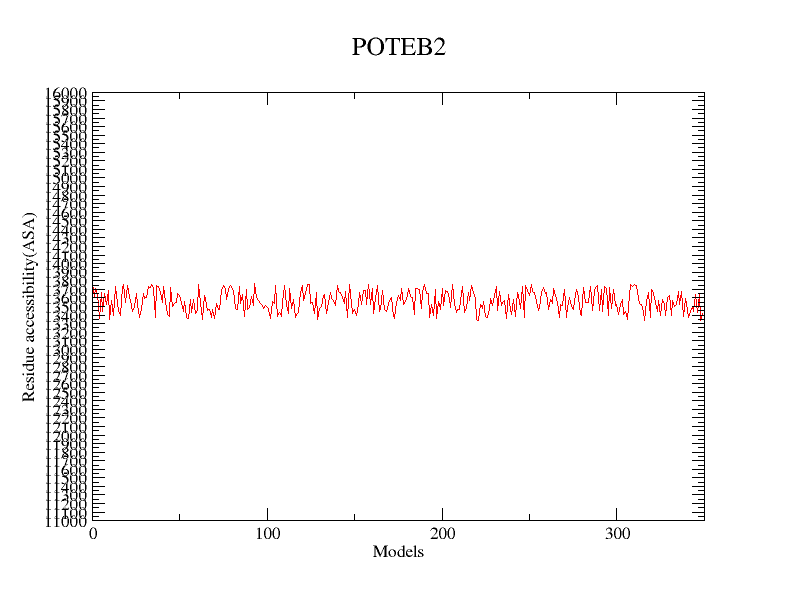

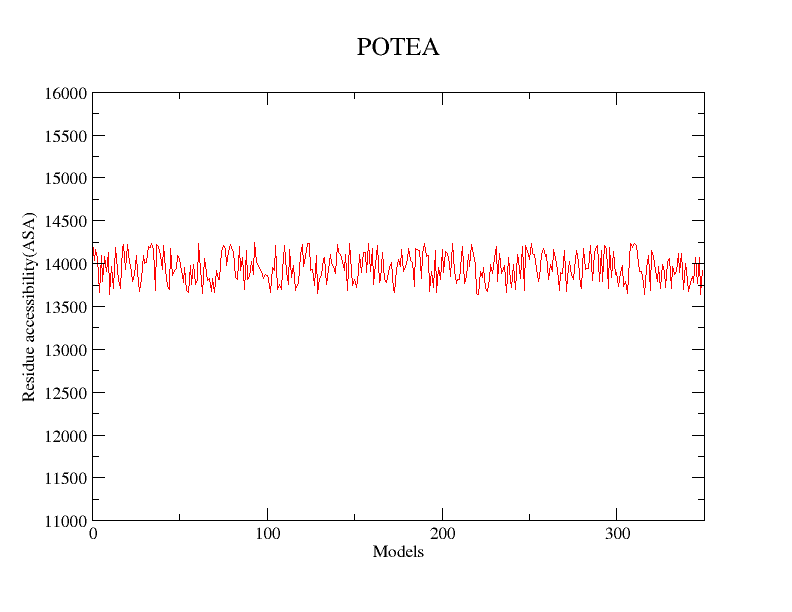

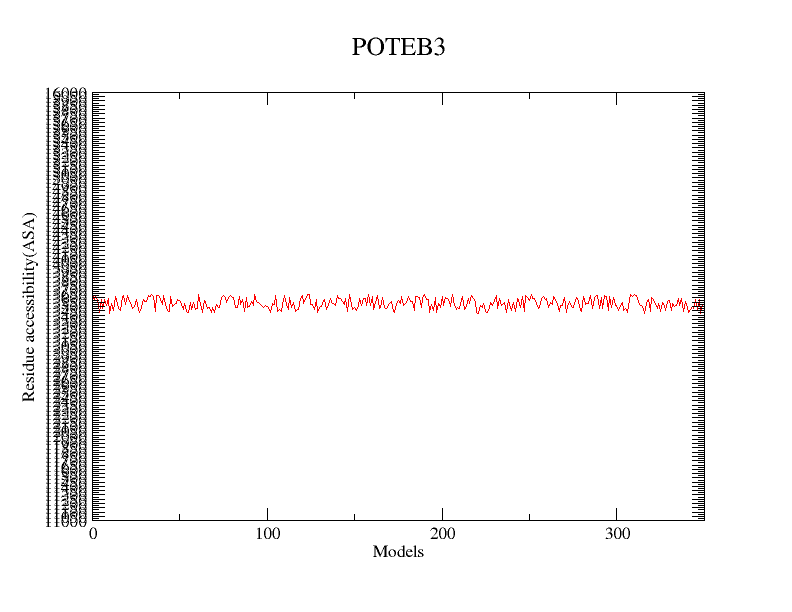

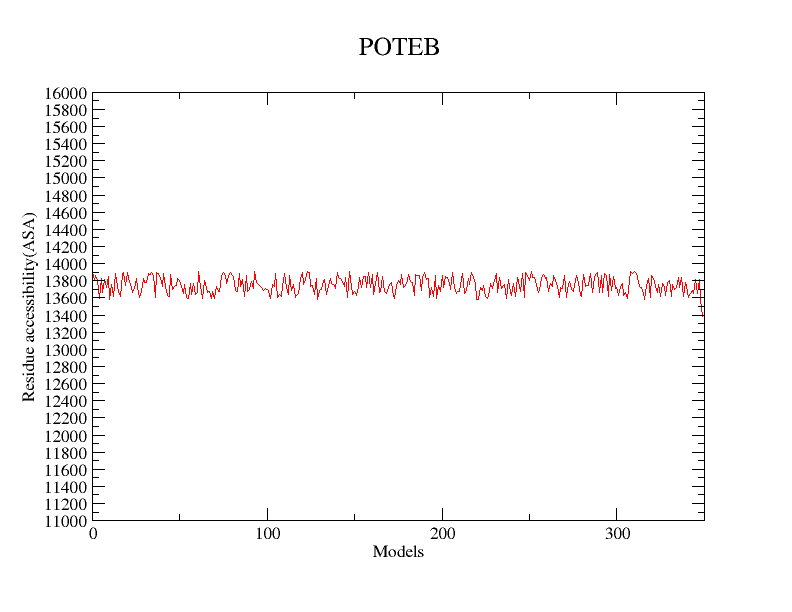

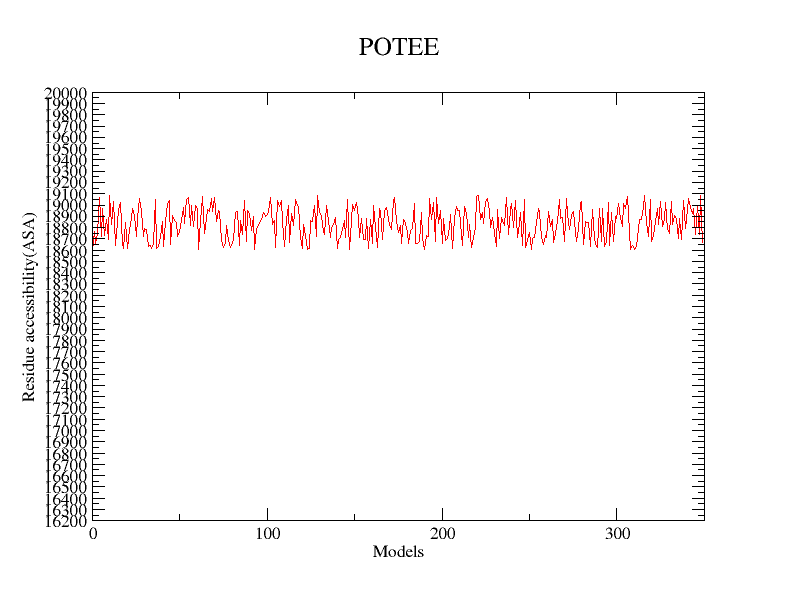

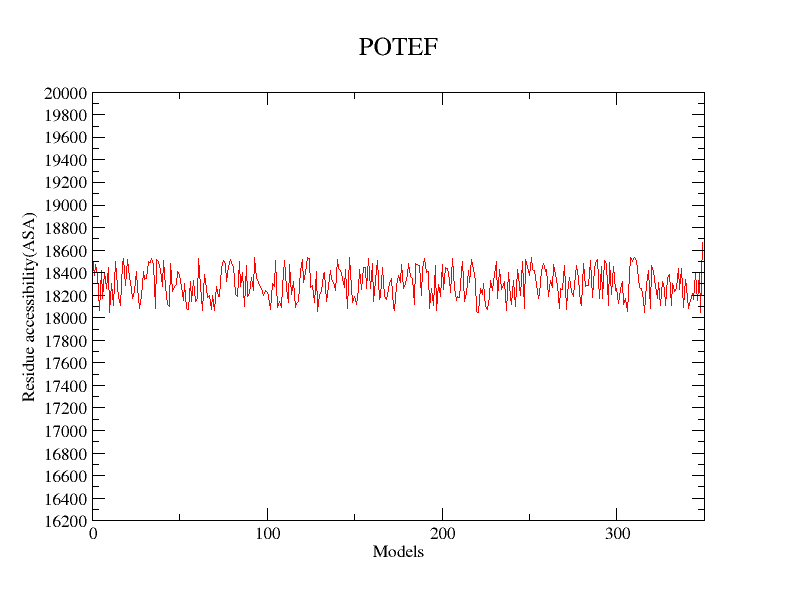

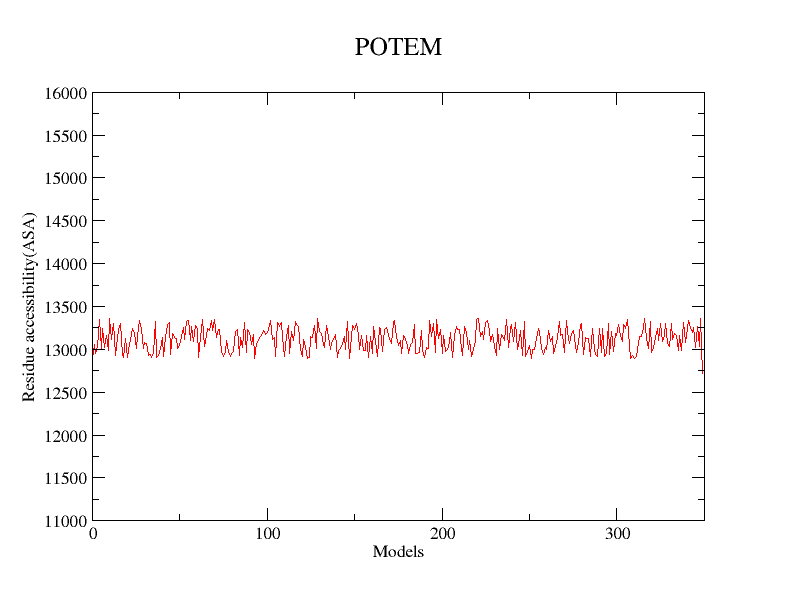

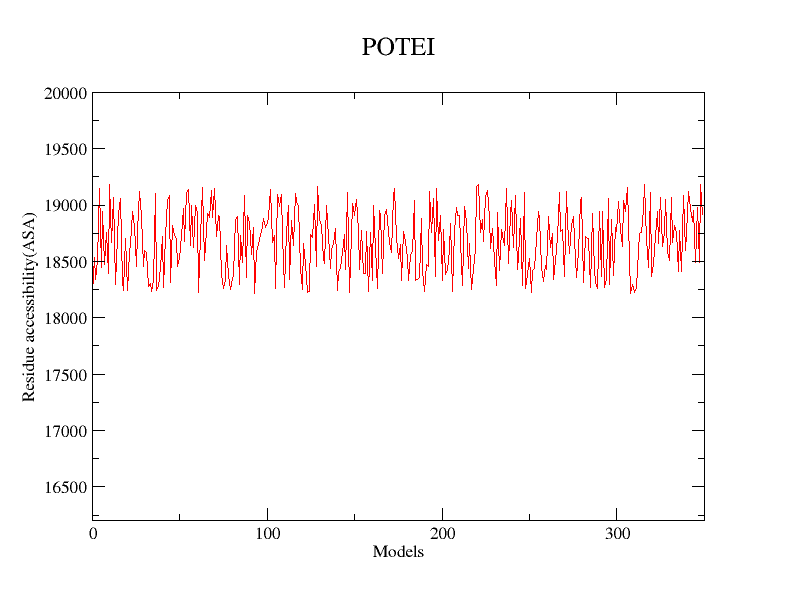

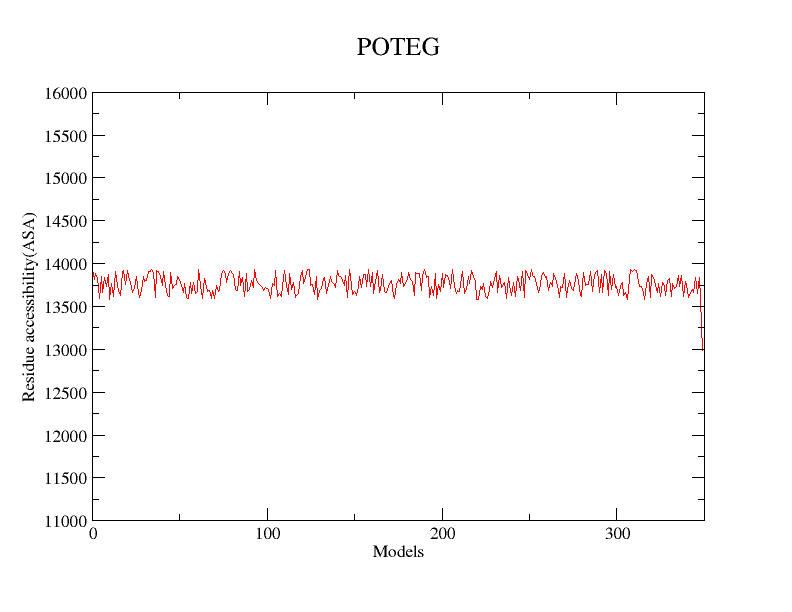

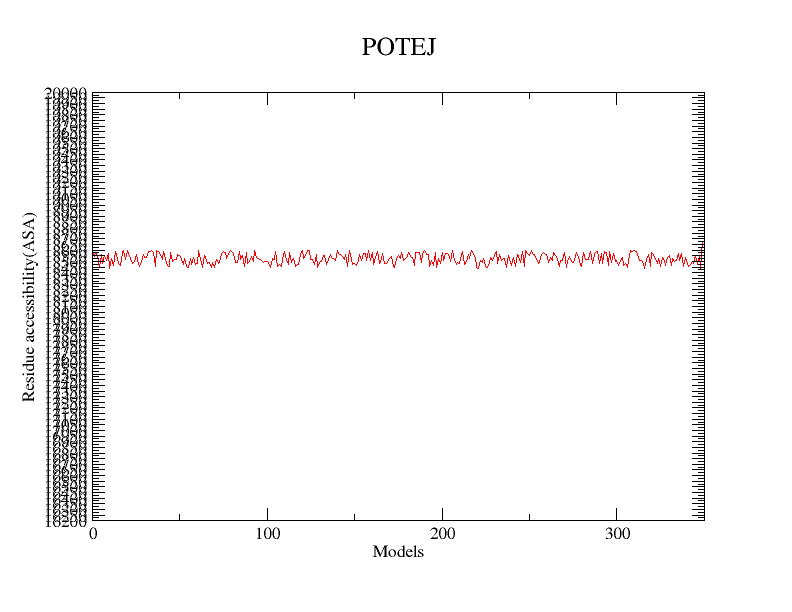

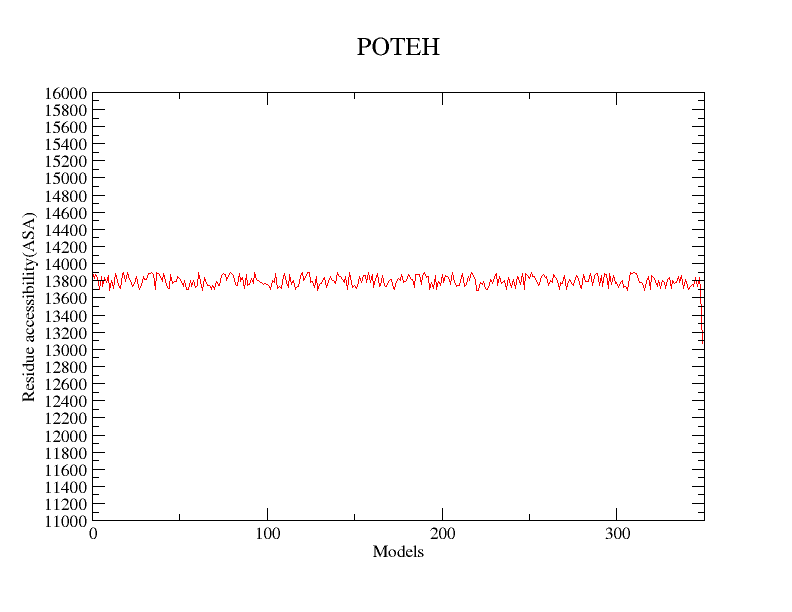

**Supplemental Figure S5: Residue accessibility vs Models (30 ns). Graph showing the trend of ASA in all POTE paralogs.**
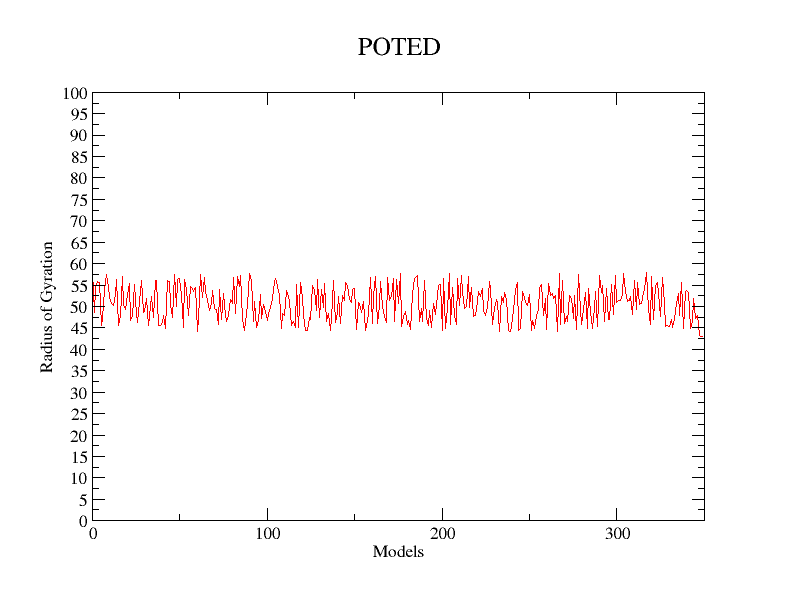

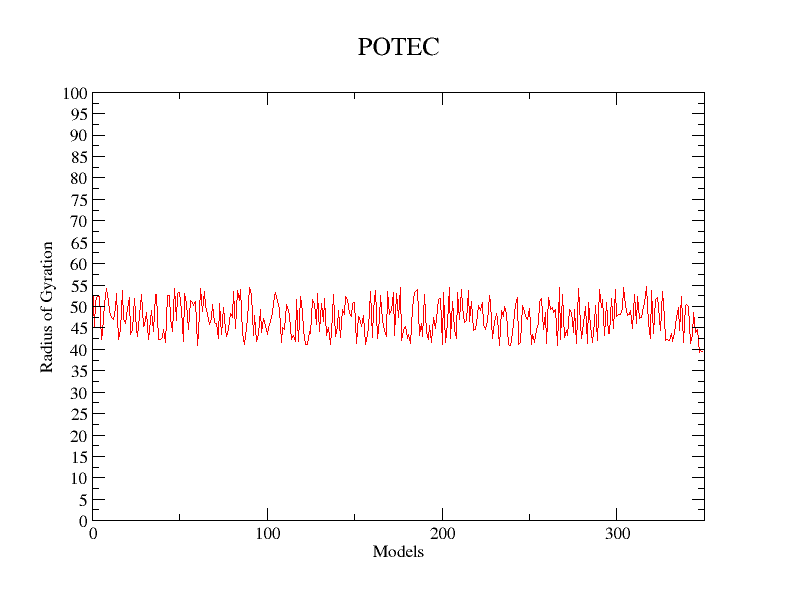

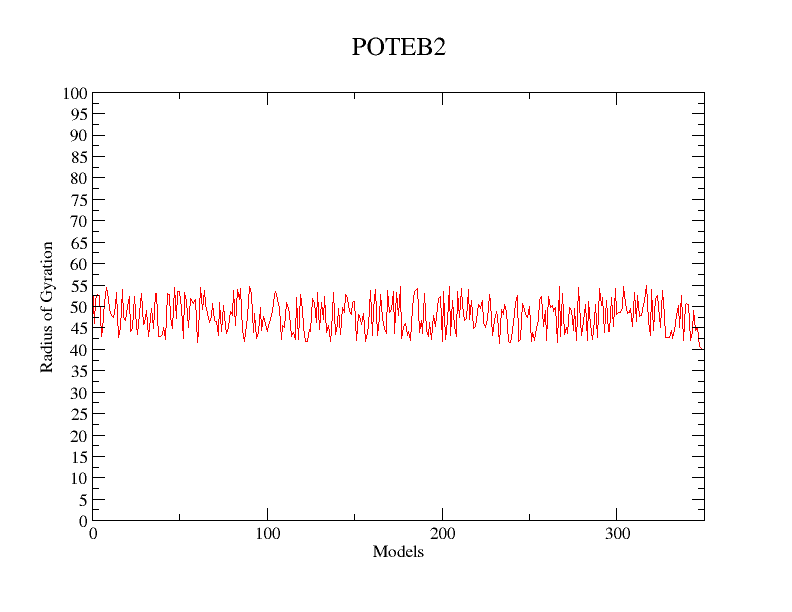

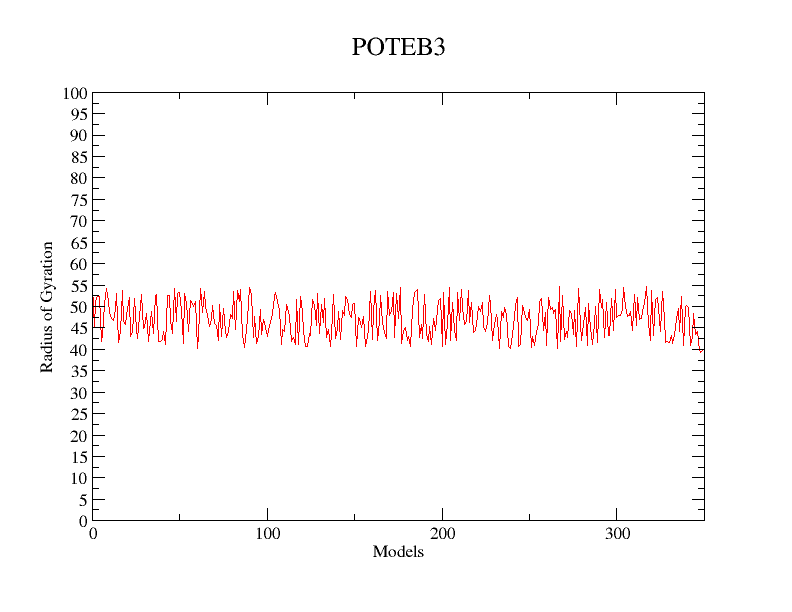

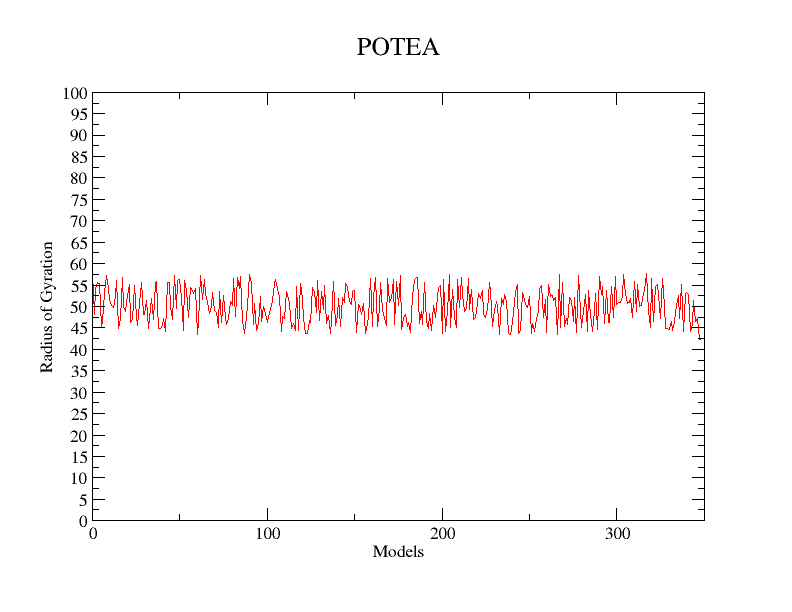

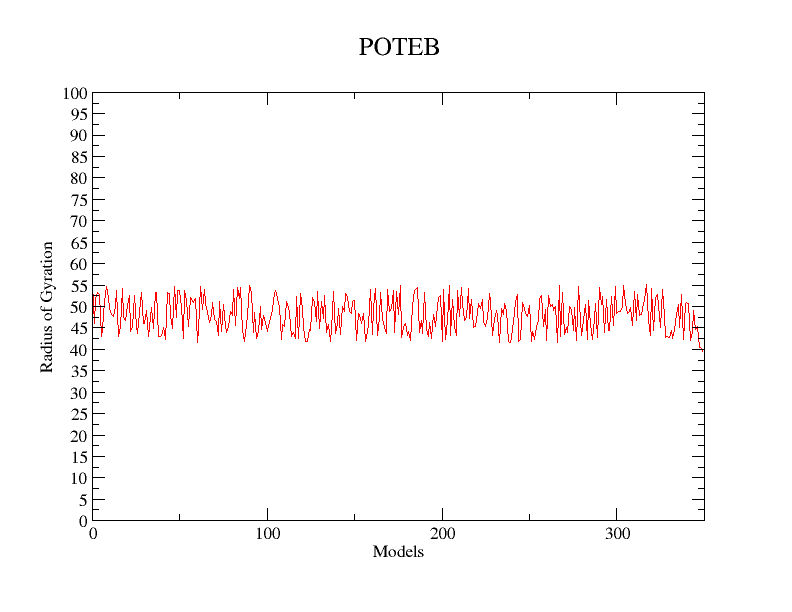

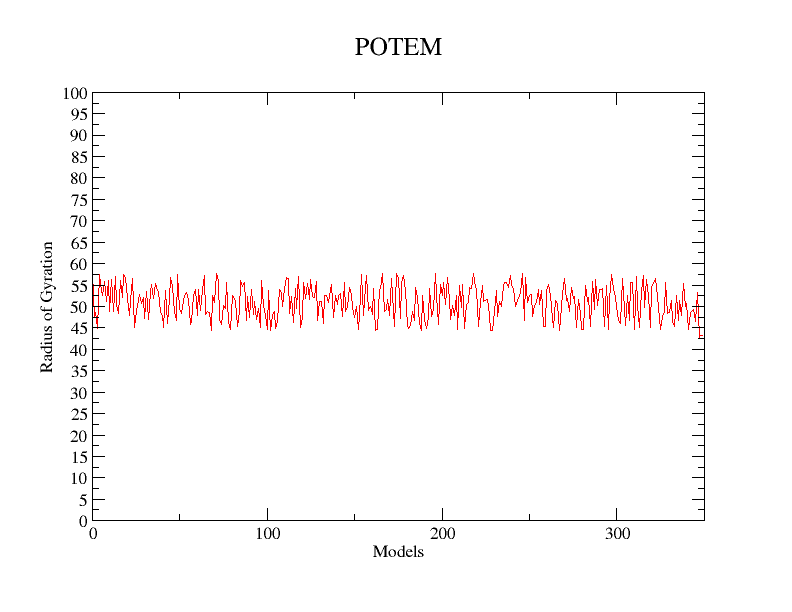

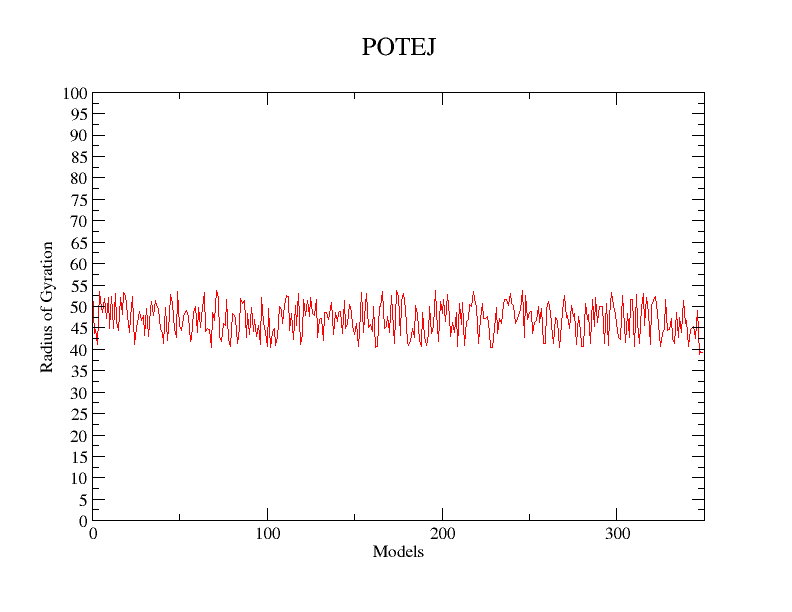

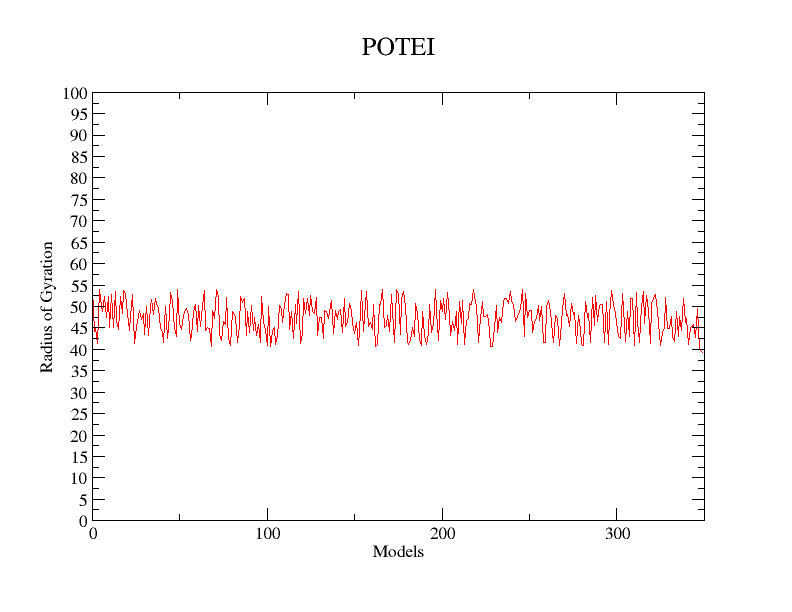

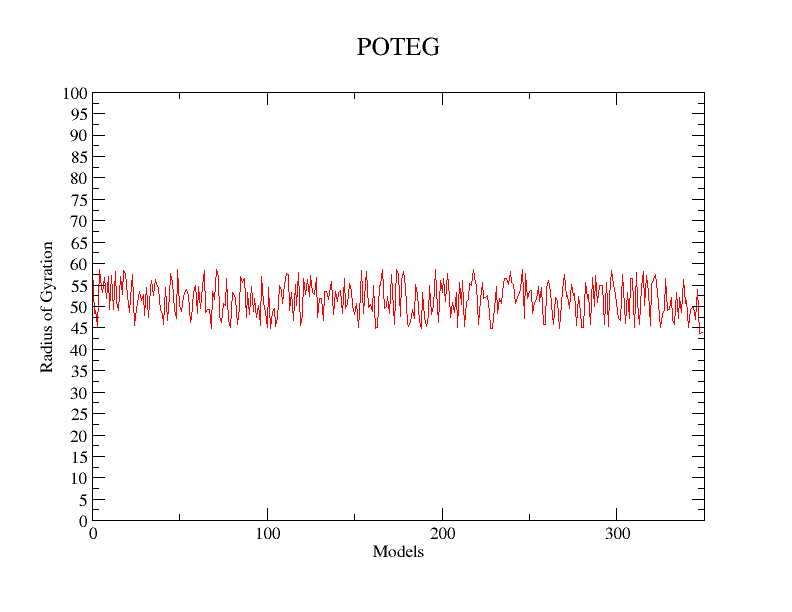

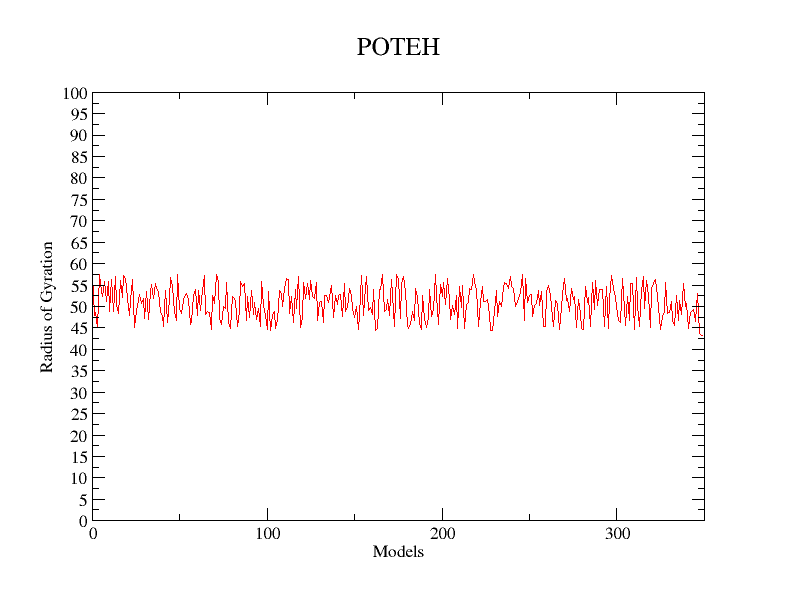

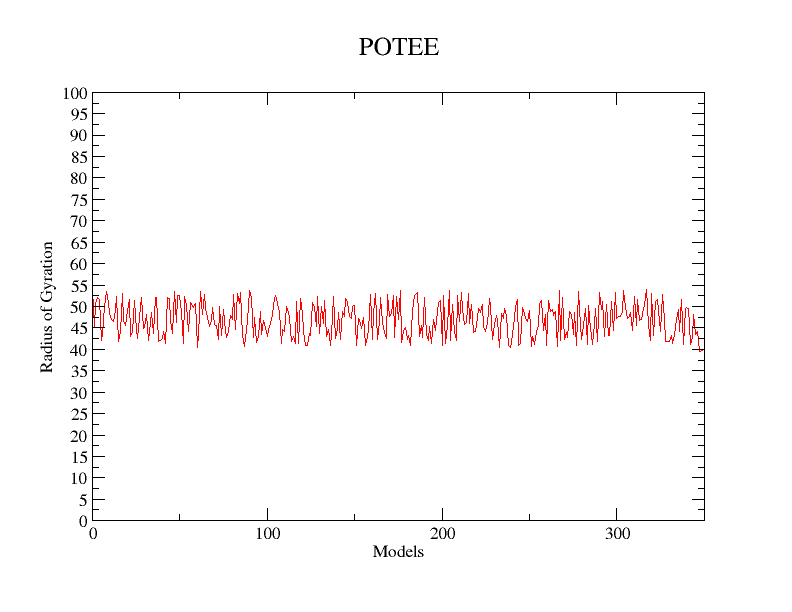

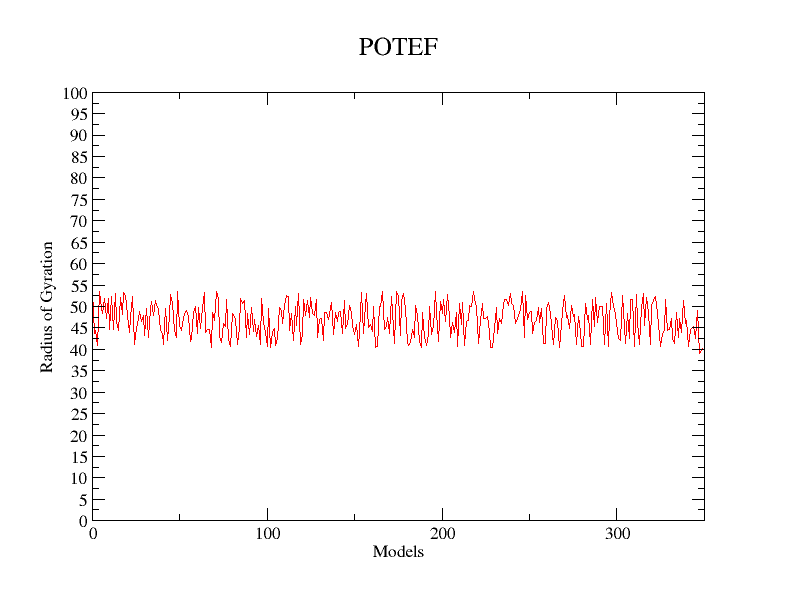

**Supplemental Figure S6: Radius of Gyration vs Models (30 ns)**. **Graph showing the trend of radius of gyration in all POTE paralogs.**
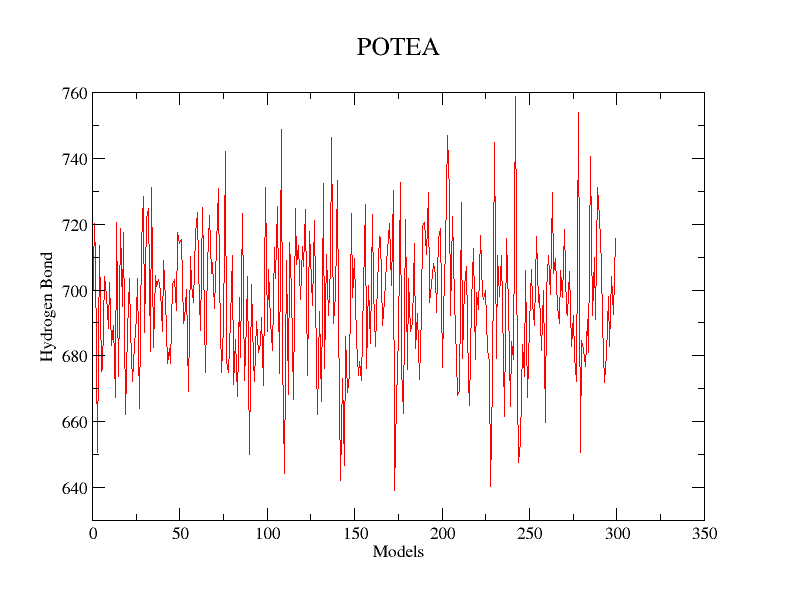

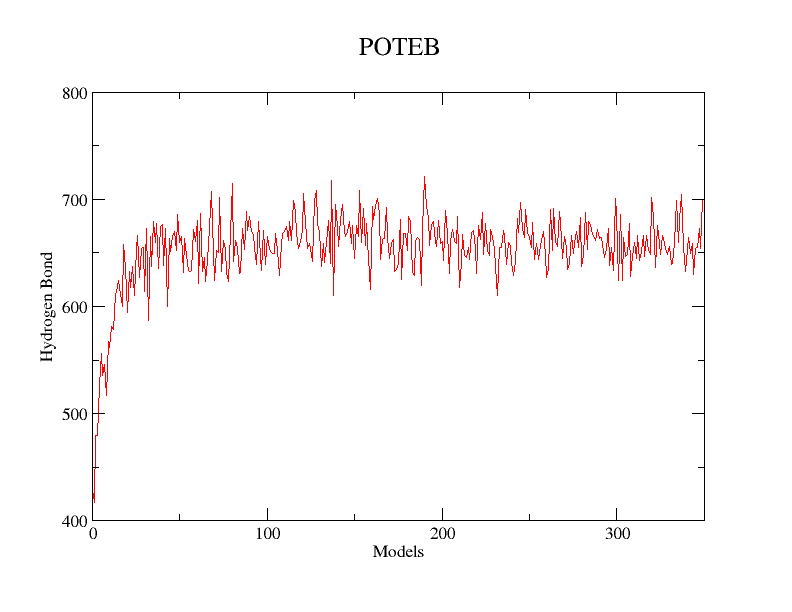

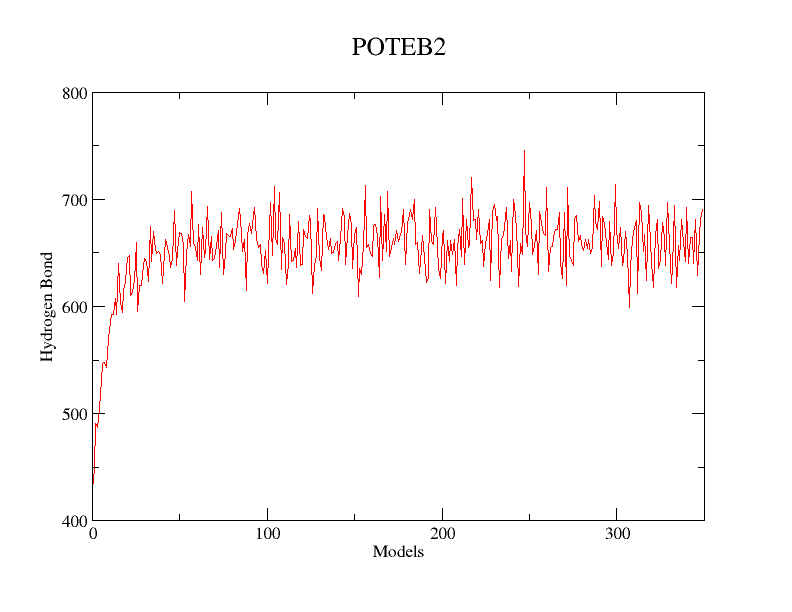

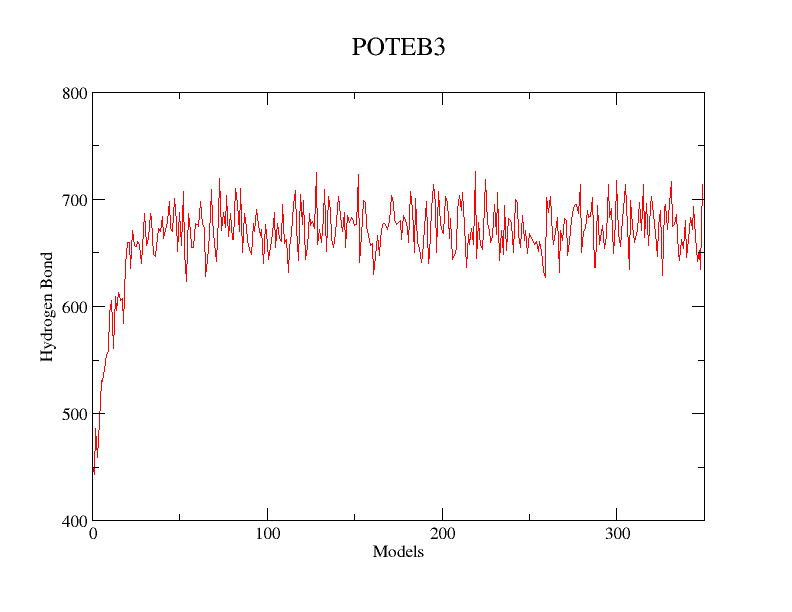

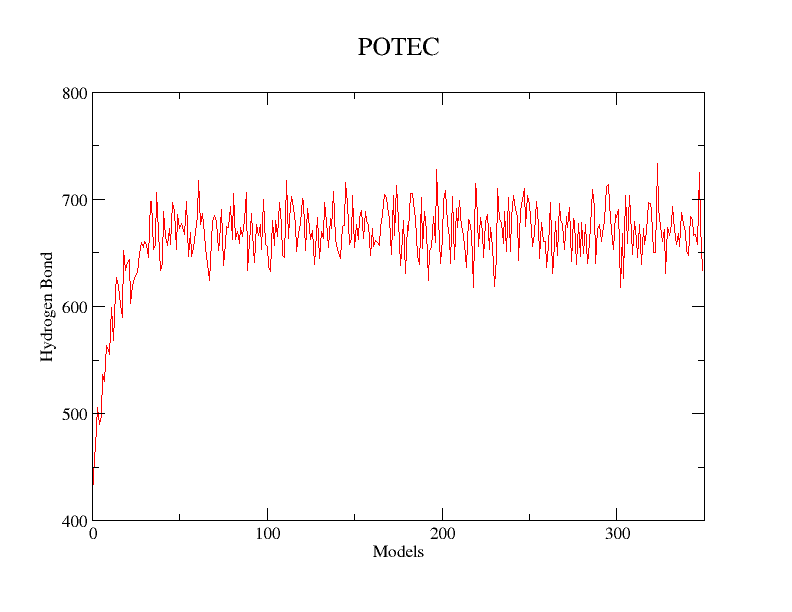

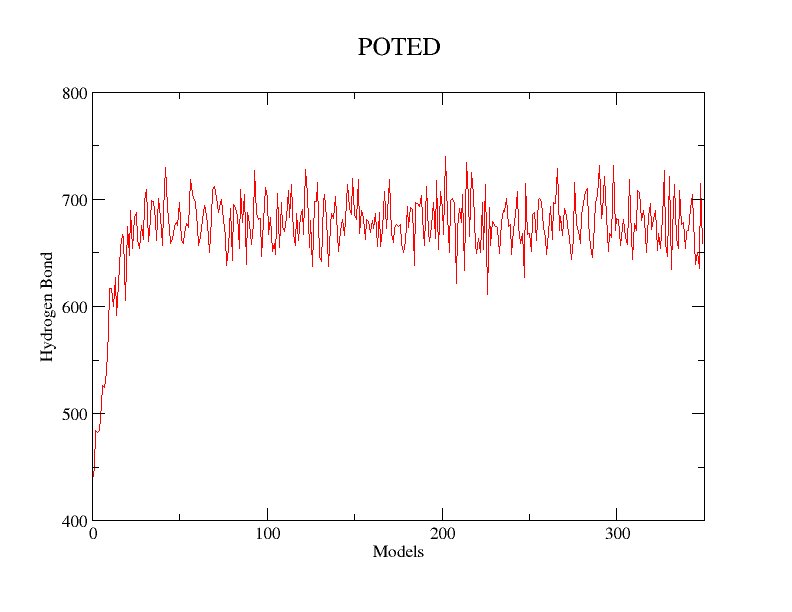

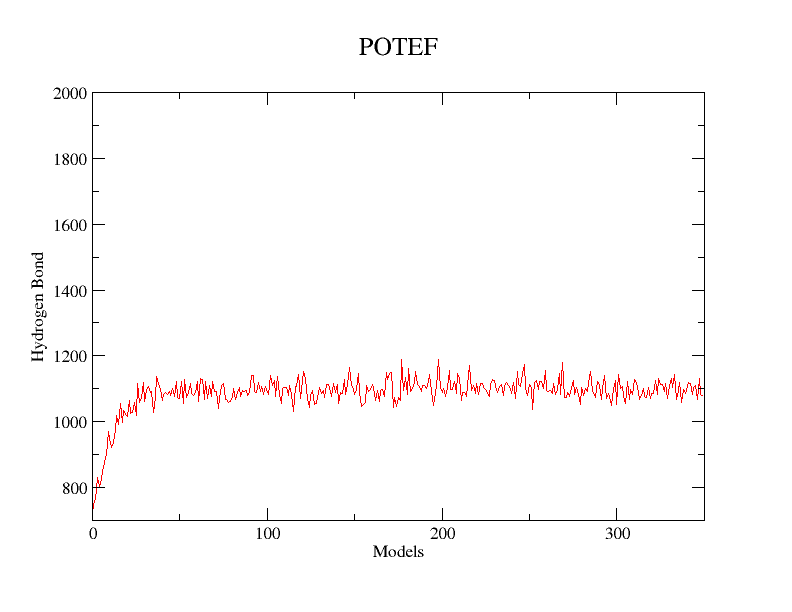

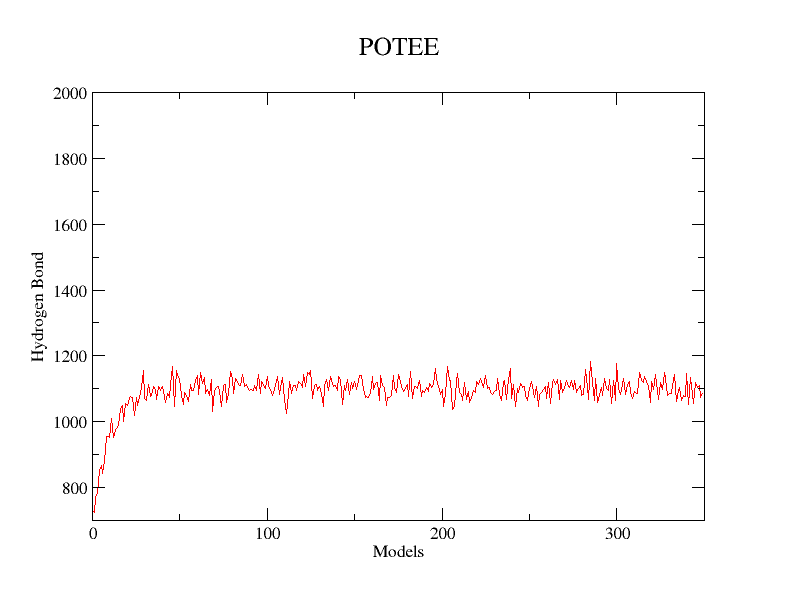

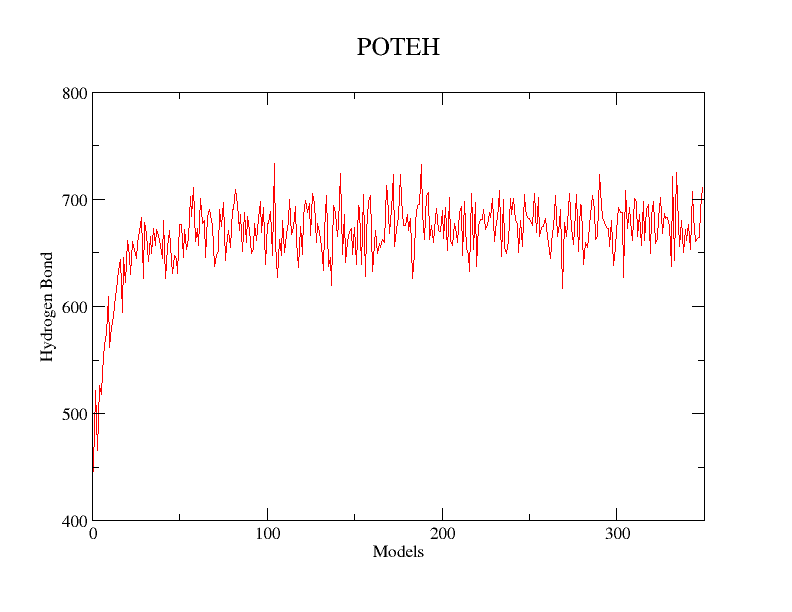

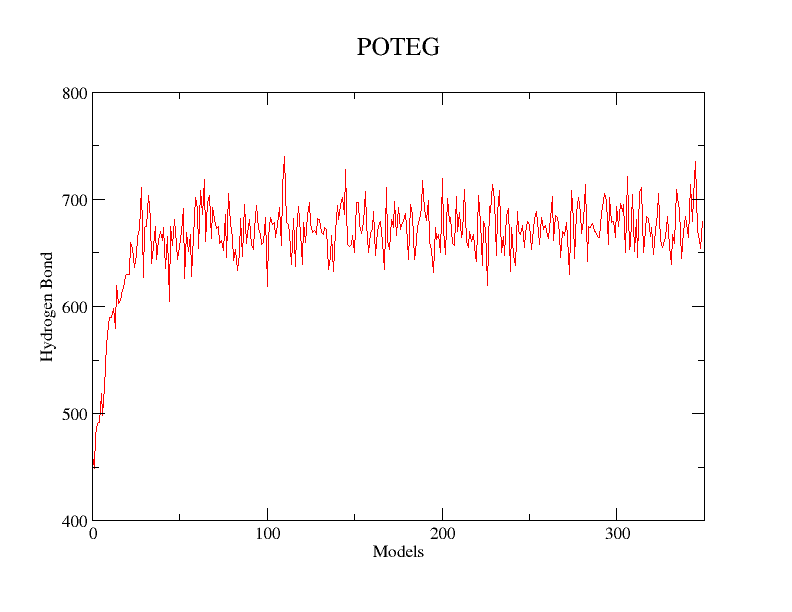

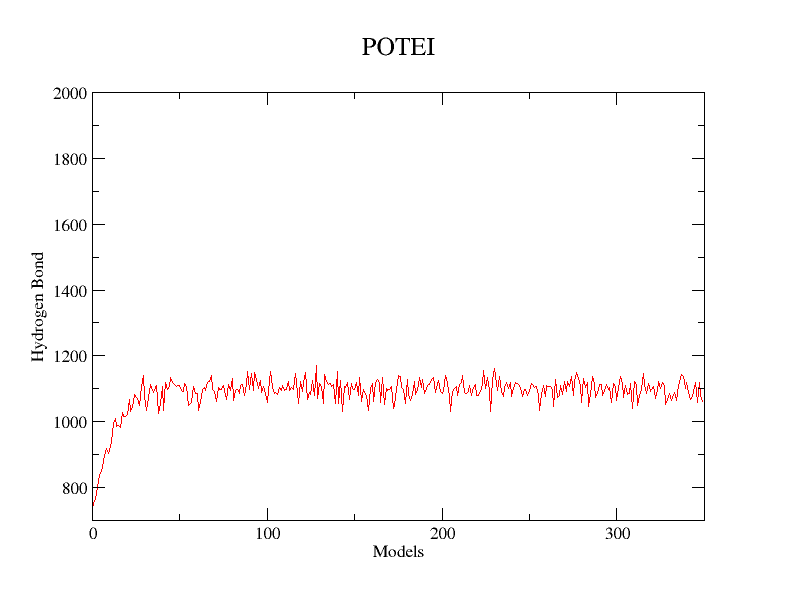

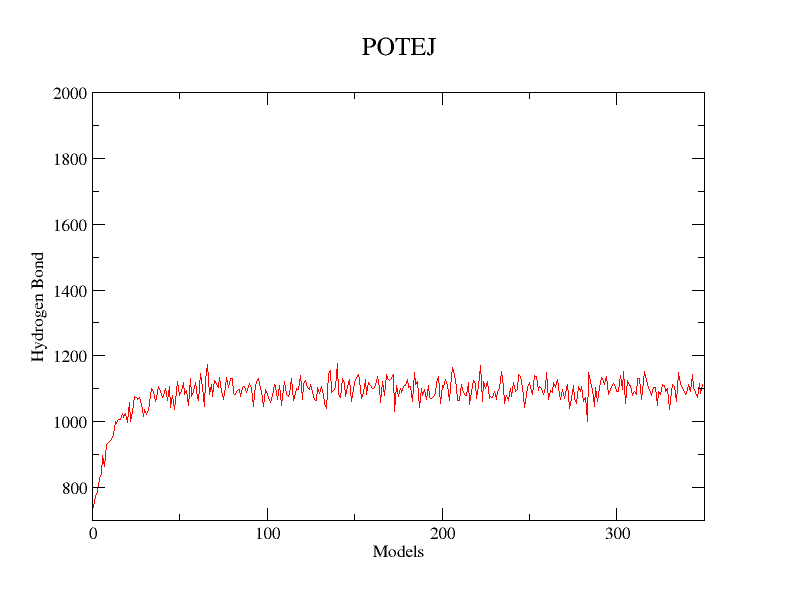

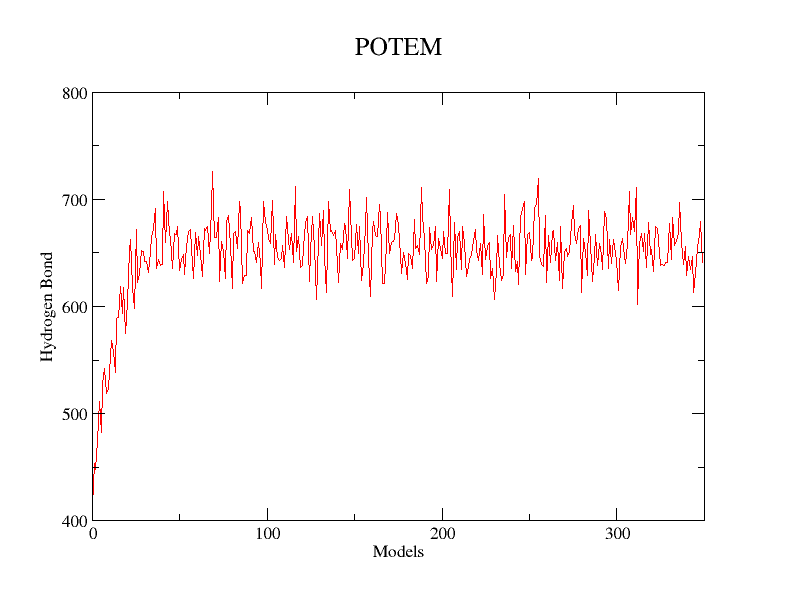

**Supplemental Figure S7: Hydrogen bonds vs Models (30ns). Graph showing the trend of hydrogen bond in all POTE paralogs**.
