## Supplemental Table for "BESFA: Bioinformatics based Evolutionary, Structural & Functional Analysis of Prostrate, Placenta, Ovary, Testis, and Embryo (POTE) Paralogs"

**Supplementary Table 1: Template for POTE paralog with E-value, Query Coverage & Sequence Similarity.**

| **Accession Number** | **Query Coverage** | **E- Value** | **Identity** |
| --- | --- | --- | --- |
| **6CXI A** | **34%** | **0.0** | **91%** |
| **3U4L A** | **34%** | **0.0** | **92%** |
| **2OAN A** | **34%** | **0.0** | **92%** |
| **4EFH A** | **34%** | **0.0** | **89%** |
| **5JHL A** | **34%** | **0.0** | **91%** |
| **5NW4 V** | **34%** | **0.0** | **91%** |
| **3BYH A** | **34%** | **0.0** | **92%** |
| **1D4X A** | **34%** | **0.0** | **90%** |
| **2BTF A** | **34%** | **0.0** | **92%** |
| **3J82 A** | **34%** | **0.0** | **92%** |
| **2HF3 A** | **34%** | **0.0** | **90%** |
| **4JHD A** | **34%** | **0.0** | **90%** |
| **4JHD B** | **34%** | **0.0** | **90%** |
| **3EKS A** | **34%** | **0.0** | **90%** |

**Supplementary Table 2:** **Sub-cellular localization of POTE paralogs.**

| **POTE Family** | **WoLF PSORT** | **Hum-m PLoc** | **DeepLoc** |
| --- | --- | --- | --- |
| POTE A | Mitochondria | Cytoplasm | Cell membrane |
| POTE B | Nucleus | Extracellular | Cell membrane |
| POTE B2 | Nucleus | Extracellular | Cell membrane |
| POTE B3 | Nucleus | Extracellular | Cell membrane |
| POTE C | Nucleus | Extracellular | Cell membrane |
| POTE D | Plasma | Extracellular | Cell membrane |
| POTE E | Plasma | Cytoplasm | Cytoplasm |
| POTE F | Plasma | Cytoplasm | Cytoplasm |
| POTE G | Plasma | Extracellular | Cytoplasm |
| POTE H | Nucleus | Extracellular | Cell membrane |
| POTE I | Plasma | Cytoplasm | Cytoplasm |
| POTE J | Plasma | Cytoplasm | Cytoplasm |
| POTE KP | Cytoplasm | Cytoplasm | Cytoplasm |
| POTE M | Plasma | Cytoplasm | Cytoplasm |

**Supplementary Table 3: Data of HUM_MPLOC**

| Centrosome | -1.24 | -1.40 | -1.40 | -1.32 | -1.29 | -1.33 | -1.68 | -1.74 | -1.26 | -1.35 | -1.74 | -1.69 | -1.25 | -1.82 |
| --- | --- | --- | --- | --- | --- | --- | --- | --- | --- | --- | --- | --- | --- | --- |
| CytoPlasm | -0.24 | -0.64 | -0.64 | -0.66 | -0.31 | -0.54 | 1.86 | 1.98 | -0.27 | -0.19 | 1.82 | 1.85 | -0.09 | 1.90 |
| Cytosol | -1.49 | -1.42 | -1.42 | -1.36 | -1.16 | -1.28 | -0.07 | -0.18 | -1.21 | -1.28 | -0.25 | -0.19 | -1.23 | -0.53 |
| Endosome | -1.23 | -1.35 | -1.35 | -1.10 | -1.33 | -1.16 | -2.64 | -2.65 | -1.47 | -1.70 | -2.71 | -2.41 | -1.64 | -3.37 |
| ER | -1.61 | -1.77 | -1.77 | -1.75 | -1.83 | -1.83 | -1.41 | -1.51 | -1.93 | -2.07 | -1.35 | -1.48 | -2.03 | -1.47 |
| Extrac | -0.46 | -0.12 | -0.12 | 0.50 | 0.15 | 0.40 | -3.59 | -3.55 | 0.30 | -0.16 | -3.82 | -3.09 | -0.10 | -4.19 |
| Golgi | -1.60 | -1.86 | -1.86 | -2.30 | -2.07 | -2.33 | -2.12 | -2.03 | -2.15 | -1.95 | -2.00 | -2.02 | -2.29 | -1.91 |
| Lysosome | -2.69 | -2.22 | -2.22 | -1.93 | -2.15 | -1.67 | -2.92 | -3.02 | -2.25 | -2.54 | -2.82 | -2.78 | -2.28 | -2.69 |
| Mitochondria | -3.37 | -3.09 | -3.09 | -3.40 | -3.17 | -3.37 | -2.72 | -2.88 | -3.17 | -3.06 | -2.71 | -2.95 | -3.18 | -2.83 |
| Nucleus | -0.87 | -1.16 | -1.16 | -1.04 | -1.04 | -1.11 | -0.71 | -0.88 | -1.02 | -0.86 | -0.70 | -0.91 | -0.91 | -0.76 |
| Peroxisome | -1.24 | -1.40 | -1.40 | -1.65 | -1.63 | -1.58 | -1.38 | -1.37 | -1.58 | -1.61 | -1.33 | -1.46 | -1.61 | -1.17 |
| Plasma | -1.48 | -1.53 | -1.53 | -1.78 | -1.64 | -1.64 | -1.66 | -1.57 | -1.46 | -1.55 | -1.71 | -1.44 | -1.59 | -1.42 |
| POTE PARALOG | POTEA | POTEB | POTEB2 | POTEB3 | POTEC | POTED | POTEE | POTEF | POTEG | POTEH | POTEI | POTEJ | POTEM | POTEKP |
| S.NO. | 1 | 2 | 3 | 4 | 5 | 6 | 7 | 8 | 9 | 10 | 11 | 12 | 13 | 14 |

**Supplementary Table 4: Data of Wolf_PSOT**

| **S.No** | **POTE** | **Mit** | **nucl** | **E.R._Mit** | **cyto_nuc** | **Cytop** | **Golgi** | **Peroxy-some** | **plasma** | **Extr** | **cysk** |
| --- | --- | --- | --- | --- | --- | --- | --- | --- | --- | --- | --- |
| 1. | POTEA | 12.5 | 6.5 | 7 | 6 | 4.5 | 5 | 2 | 1 |  |  |
| 2. | POTEB | 4.5 | 6.5 | 3 | 6.5 | 5.5 | 5 | 4 | 4 | 2 |  |
| 3. | POTEB2 | 4.5 | 6.5 | 3 | 6.5 | 5.5 | 5 | 4 | 4 | 2 |  |
| 4. | POTEB3 | 4.5 | 6.5 | 3 | 5 | 2.5 | 5 | 1 | 6 | 6 |  |
| 5. | POTEC | 5.5 | 10 | 3.5 | 8 | 4 |  |  | 7 | 5 |  |
| 6. | POTED | 4.5 | 10 | 3 | 8 | 4 |  | 1 | 12 |  |  |
| 7. | POTEE |  | 11.5 |  | 7.5 | 2.5 |  | 1 | 14 |  | 3 |
| 8. | POTEF |  | 12 |  |  | 2 |  | 1 | 14 |  | 3 |
| 9. | POTEG | 4.5 | 11 | 3 | 7.5 | 2 |  |  | 14 |  |  |
| 10. | POTEH | 3 | 15 |  | 9.5 | 2 |  |  | 12 |  |  |
| 11. | POTEI |  | 12 |  |  | 3 |  |  | 14 |  | 3 |
| 12. | POTEJ |  | 10.5 |  | 7.5 | 3.5 |  |  | 15 |  | 3 |
| 13. | POTEKP | 1 | 8.5 |  | 17 | 20.5 |  | 2 |  |  |  |
| 14. | POTEM | 4.5 | 11 | 3 | 7.5 | 2 |  |  | 14 |  |  |

**Mit=mitochondria; nucl = nucleus; E.R.= endoplasmic reticulum; Cytop = cytoplasm;**

**Supplementary Table 5:** **Function Prediction of predicted structure.**

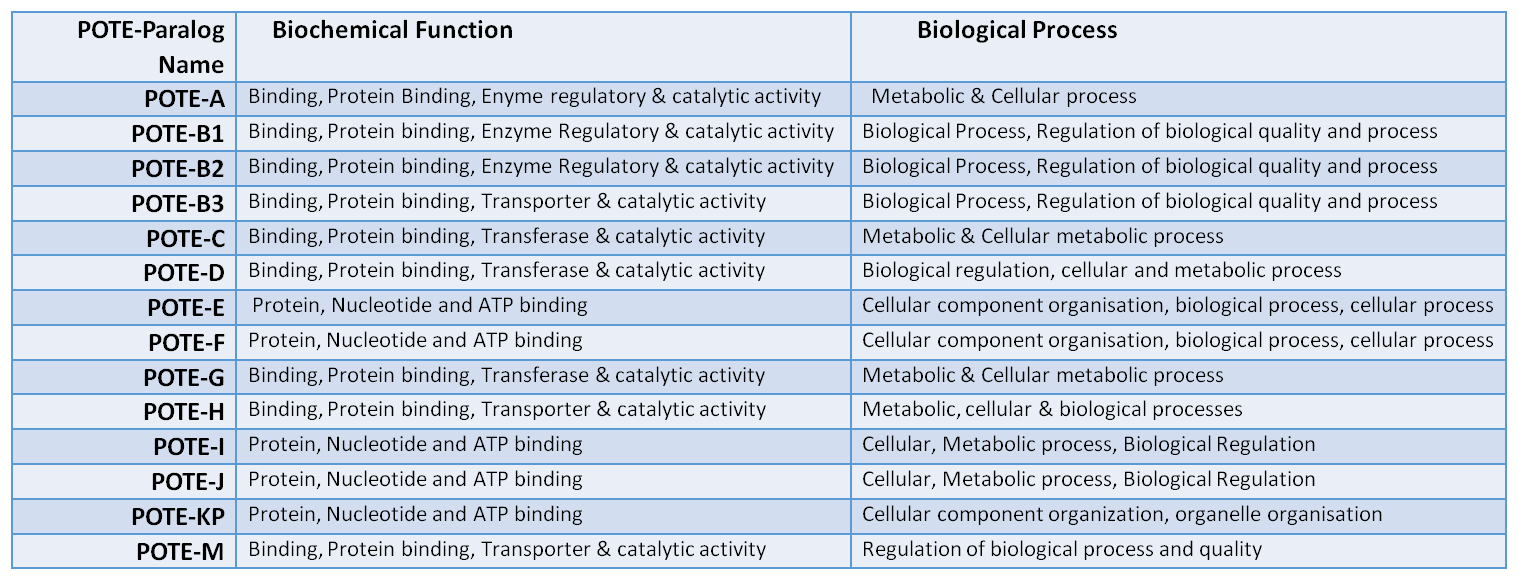

**Supplementary Table 6: List of selected anticancer drug compounds with higher binding energy.**

| Sl. No. | Cancer Type | Ligand Id | Structure |
| --- | --- | --- | --- |
| 1 | Ovarian | 1 | 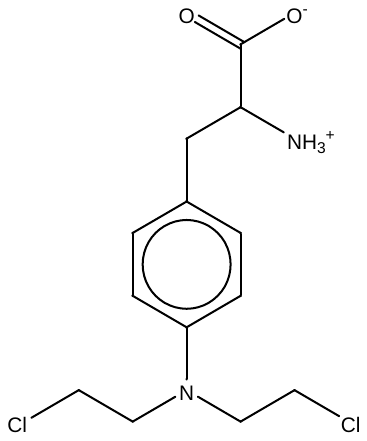 |
| 2 | Testicular | 2 | 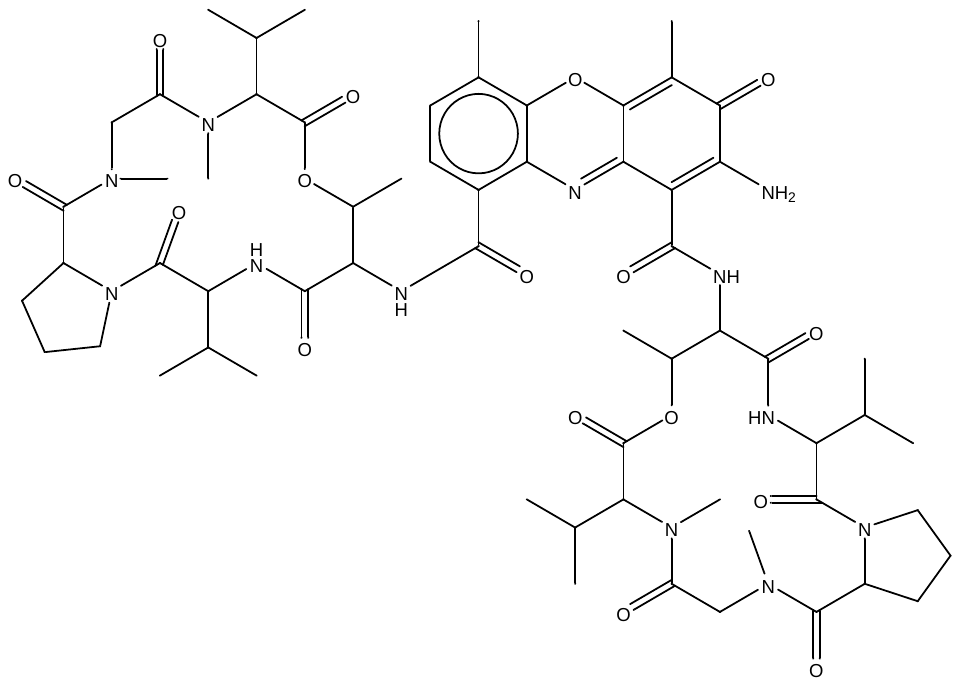 |
| 3 | Testicular | 3 | 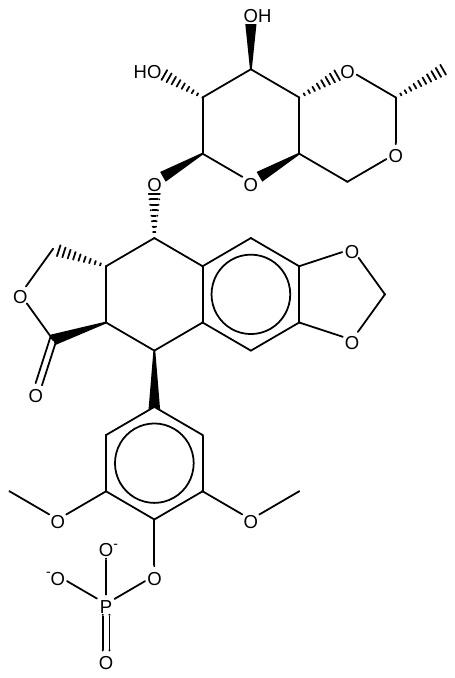 |
| 4 | Prostate | 3 | 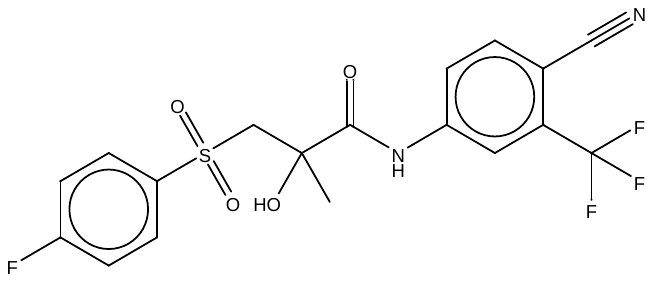 |
| 5 | Testicular | 4 | 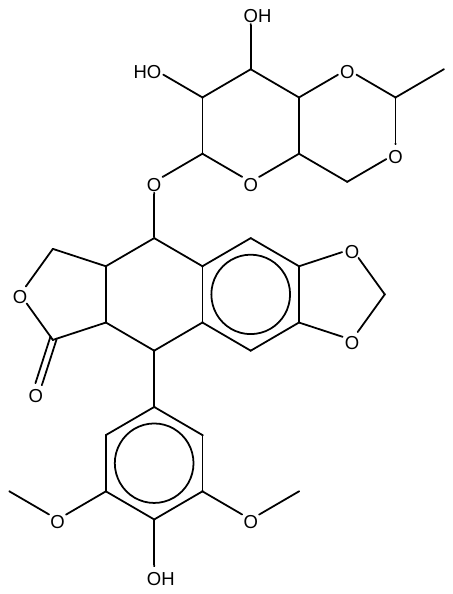 |
| 6 | Ovarian | 4 | 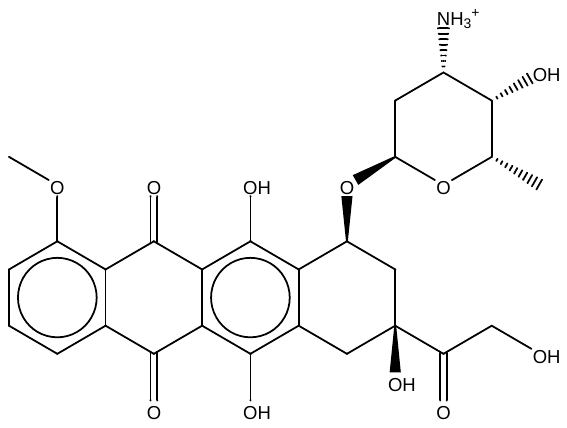 |
| 7 | Ovarian | 5 | 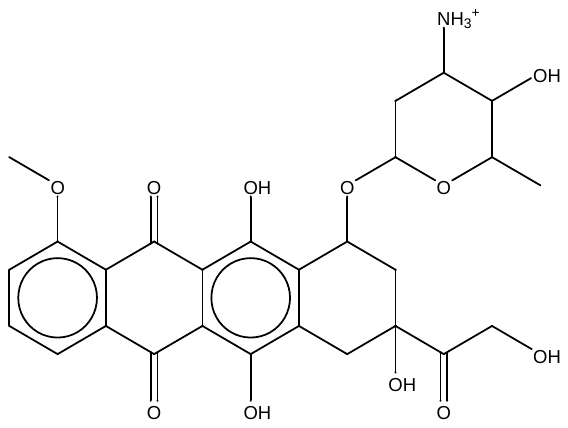 |
| 8 | Prostate | 6 | 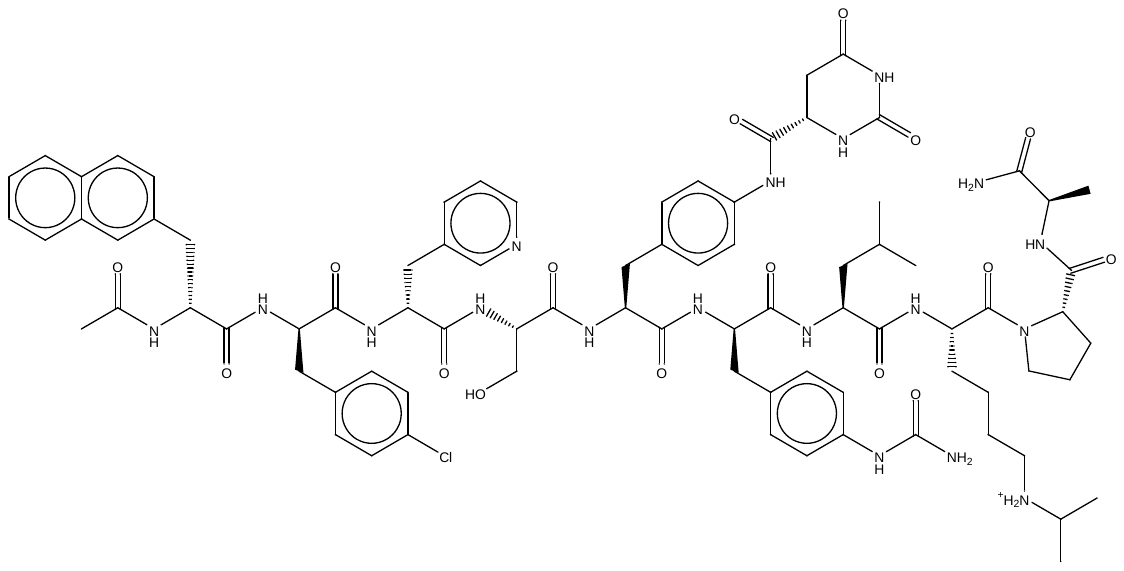 |
| 9 | Prostate | 8 | 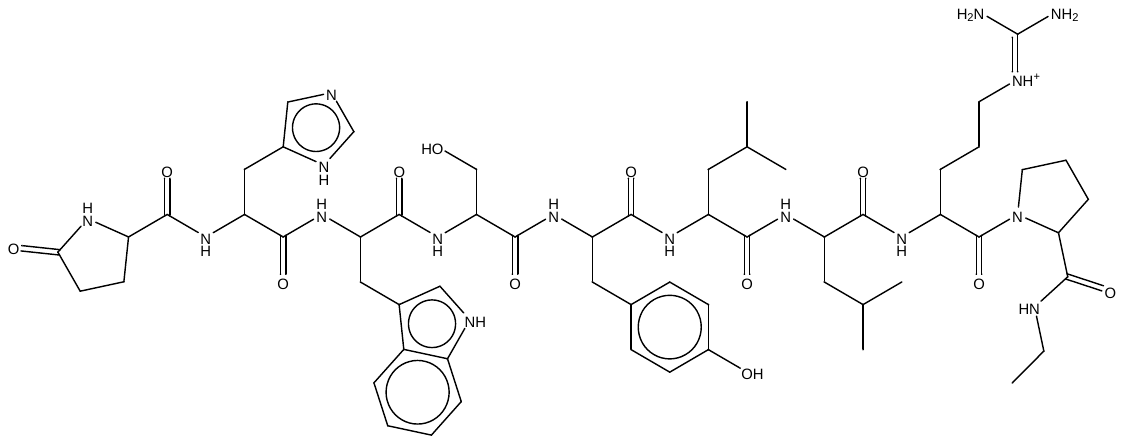 |
| 10 | Prostate | 10 | 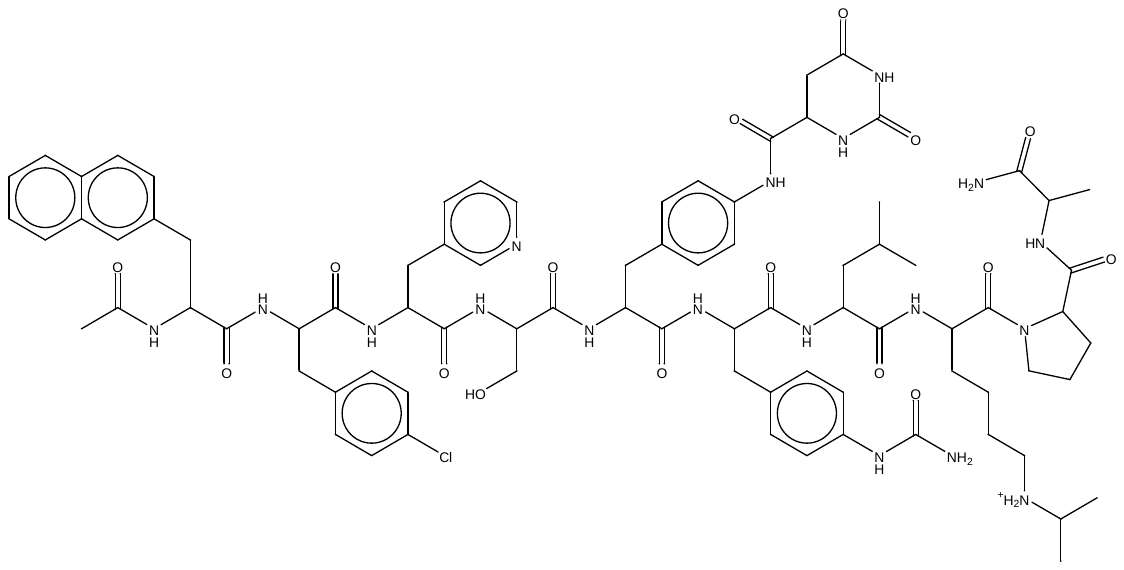 |
| 11 | Ovarian | 12 | 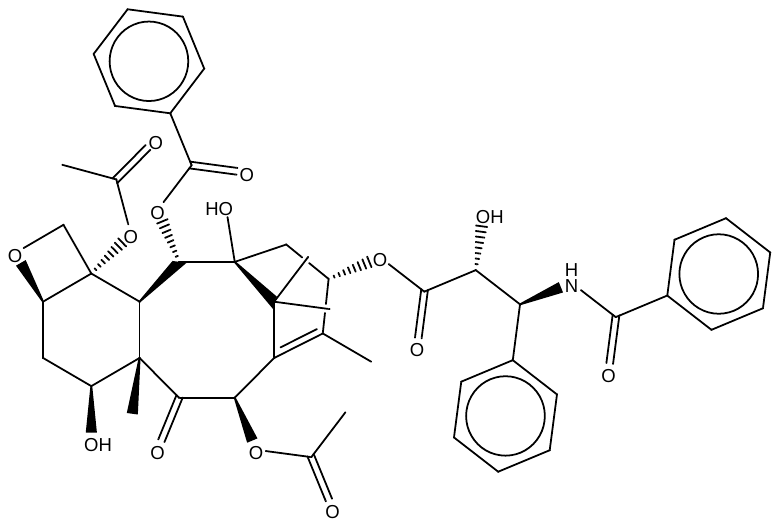 |
| 12 | Ovarian | 14 | 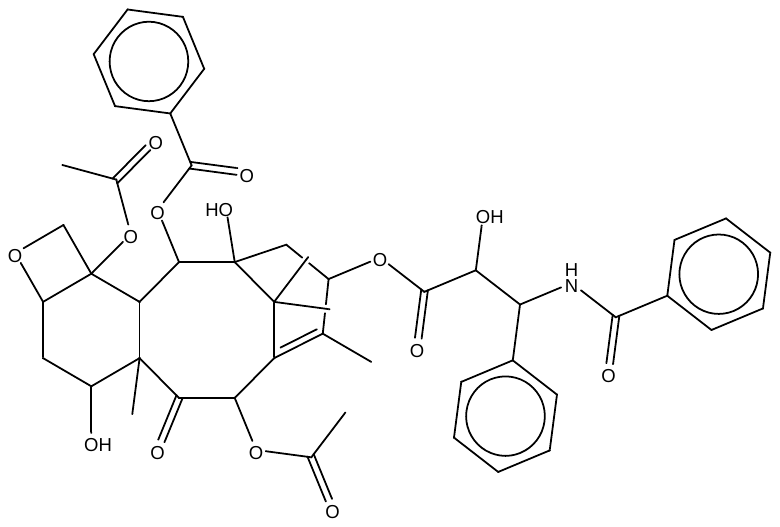 |
| 13 | Prostate | 15 | 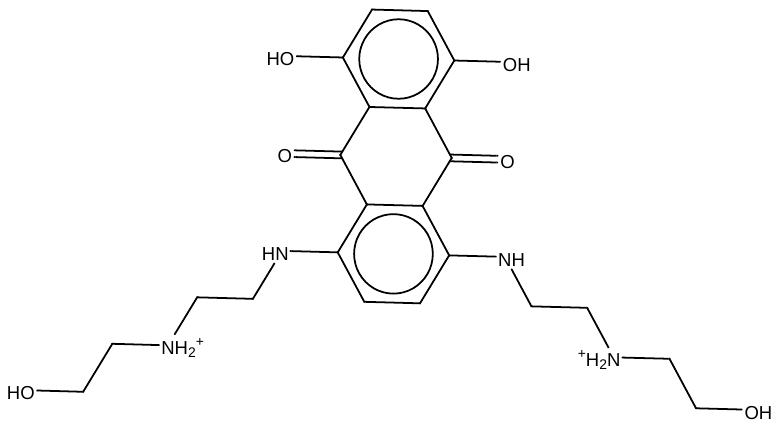 |

**Supplementary Table 7:** **List of selected drugs with binding energy, POTE proteins and cancer type.**

| Sl. No. | Protein Name | Cancer Type | Ligand Id | Binding Energy (kcal/mol) |
| --- | --- | --- | --- | --- |
| 1 | POTEA | Ovarian | 4 | -11.1231 |
| 2 | POTEA | Prostate | 10 | -11.6997 |
| 3 | POTEA | Testicular | 1 | -10.2206 |
| 4 | POTEB | Ovarian | 5 | -14.6565 |
| 5 | POTEB | Prostate | 10 | -11.6997 |
| 6 | POTEB | Testicular | 2 | -11.2449 |
| 7 | POTEB2 | Ovarian | 5 | -15.1491 |
| 8 | POTEB2 | Prostate | 10 | -13.3722 |
| 9 | POTEB2 | Testicular | 1 | -12.1947 |
| 10 | POTEB3 | Ovarian | 12 | -8.37356 |
| 11 | POTEB3 | Prostate | 6 | -12.4264 |
| 12 | POTEB3 | Testicular | 2 | -9.93672 |
| 13 | POTEC | Ovarian | 5 | -14.5191 |
| 14 | POTEC | Prostate | 8 | -11.1837 |
| 15 | POTEC | Testicular | 4 | -15.0707 |
| 16 | POTED | Ovarian | 1 | -11.8637 |
| 17 | POTED | Prostate | 15 | -12.2074 |
| 18 | POTED | Testicular | 3 | -13.1265 |
| 19 | POTEE | Ovarian | 12 | -8.29924 |
| 20 | POTEE | Prostate | 15 | -14.222 |
| 21 | POTEE | Testicular | 3 | -14.2351 |
| 22 | POTEF | Ovarian | 5 | -15.9455 |
| 23 | POTEF | Prostate | 10 | -11.3934 |
| 24 | POTEF | Testicular | 3 | -14.0163 |
| 25 | POTEG | Ovarian | 5 | -12.0845 |
| 26 | POTEG | Prostate | 3 | -11.9003 |
| 27 | POTEG | Testicular | 3 | -15.8356 |
| 28 | POTEH | Ovarian | 12 | -7.73076 |
| 29 | POTEH | Prostate | 15 | -12.5518 |
| 30 | POTEH | Testicular | 3 | -13.4496 |
| 31 | POTEI | Ovarian | 5 | -13.2094 |
| 32 | POTEI | Prostate | 10 | -12.2155 |
| 33 | POTEI | Testicular | 4 | -16.0921 |
| 34 | POTEJ | Ovarian | 4 | -15.6959 |
| 35 | POTEJ | Prostate | 15 | -14.1459 |
| 36 | POTEJ | Testicular | 4 | -13.913 |
| 37 | POTEK | Ovarian | 14 | -9.67262 |
| 38 | POTEK | Prostate | 15 | -15.0036 |
| 39 | POTEK | Testicular | 1 | -15.5574 |
| 40 | POTEM | Ovarian | 4 | -13.9724 |
| 41 | POTEM | Prostate | 6 | -10.2239 |
| 42 | POTEM | Testicular | 3 | -19.8317 |
